# Modular to cyclic TCA governs hematopoiesis in *Drosophila*

**DOI:** 10.64898/2025.12.04.692477

**Authors:** Ajay Tomar, Shaon Chakrabarti, Tina Mukherjee

## Abstract

TCA cycle is well known for its role in bioenergetics and also a hub for metabolic exchange, here we identify a developmentally programmed, non-uniform TCA architecture that is required for hematopoietic state transitions in the *Drosophila* lymph gland. Early blood progenitors operate a modular TCA configuration, in which a *CS/mAcon1*-derived citrate node preserves progenitor identity, while a *Gdh/Kdh* mediated α-ketoglutarate to succinyl-CoA branch, not succinate, supports proliferative potential. As cells exit the progenitor state and initiate lineage commitment, the cells undergo metabolic reorganization and engage a fully cyclic, *Pdh*- and *Idh*-dependent oxidative TCA cycle whose cataplerotic activity keeps the levels of TCA metabolites in check as their excess otherwise induces differentiation. Blocking these metabolic transitions disrupts progenitor homeostasis, compromises proliferative competency of transitional progenitors, and distorts balanced differentiation. These findings establish a developmentally regulated alternative to the traditional TCA cycle and reveals that appropriate engagement of modular versus cyclic TCA modes is required for orchestrating hematopoietic fate transitions. This work highlights TCA topology and not flux alone functions as a core determinant of stemness, proliferation, and lineage progression during hematopoiesis.

**Significance Statement:** Hematopoietic development requires precise metabolic control to maintain progenitors while enabling their transition into differentiated lineages. We show that the *Drosophila* lymph gland achieves this through a developmentally programmed switch in TCA-cycle configuration where early progenitors rely on a modular TCA architecture that independently sustains identity and proliferation, whereas later stages require a fully cyclic, oxidative TCA mode to maintain homeostasis and balanced differentiation. Disrupting either phase perturbs hematopoietic organization, demonstrating that TCA topology itself regulates cell-state transitions. These findings reveal TCA architecture as the determinant of hematopoietic fate and provides a conceptual framework relevant to hematopoiesis and also stem cell biology.

## Introduction

The tricarboxylic acid (TCA) cycle is one of the most versatile and deeply conserved metabolic circuits in biology. Beyond ATP generation, the TCA cycle integrates carbon from multiple sources, produces biosynthetic precursors, controls redox balance, and generates metabolites that modulate chromatin, signaling, lipid homeostasis, stress adaptation, and cellular fate decisions. Each intermediate: citrate, α-ketoglutarate (α-KG), succinate, fumarate, malate, and oxaloacetate, participates in distinct biochemical and regulatory processes, positioning the TCA cycle as a multifunctional metabolic and signaling hub essential for growth and development (1–10). Across metazoan development, however, the TCA cycle does not operate as a uniform circular loop. Instead, it exhibits context-dependent rewiring in embryos, stem cells, and differentiating tissues. Early mammalian embryos and pluripotent stem cells rely on truncated or reductive TCA flux to support rapid biosynthesis: proliferating tissues employ citrate-acetyl CoA branching; differentiated cells often establish a fully cyclic oxidative loop; while immune cells selectively upregulate discrete nodes such as succinate or itaconate-producing branches (3, 11–19). These variant, non-canonical or modular modes of TCA operation reveal that the cycle is not fixed but a flexible metabolic architecture, assembled and disassembled according to developmental and physiological demand. This complexity complicates the assumption that the TCA cycle functions as a closed circular loop in all contexts, its structure is instead dynamic, modular, and state-dependent. Finally, as the TCA cycle also integrates carbon and signals from diverse nutrient sources, this adds an additional regulatory and dynamic dimension, positioning it as a central integrator of systemic and physiological state changes. Consequently, shifts in TCA-flux can modulate cellular responses to align with organismal needs.

Hematopoiesis, the process of generating and renewing blood cells, is a classic example of state-dependent metabolic specialization. Blood progenitors and stem cells require TCA derived metabolites for identity, redox regulation, and lineage transitions. Citrate-acetyl CoA couple growth with chromatin remodeling (3, 20–24); α-KG regulates dioxygenase activity and differentiation competence (25–27); succinate and fumarate tune hypoxia, inflammatory priming, and myeloid fate (2, 26, 28–39); and mitochondrial ROS derived from TCA flux calibrate differentiation thresholds (8, 40–45). Yet, how the TCA cycle is strategically deployed across the continuum of blood-forming cells, from quiescent stem/progenitors to proliferative intermediates to differentiated lineages, remains unresolved. These uncertainties are intensified by the fact that the TCA cycle can operate either as a fully connected oxidative loop or in semi-disconnected “modules.” For instance, in proliferating or differentiating cells, the citrate branch, α-KG branch, or succinate/fumarate segments can function independently or with asymmetric flux. Given that hematopoiesis demands fundamentally different metabolic states at different stages, from quiescence, expansion, priming to differentiation, it is unclear how the hematopoietic tissue employs the TCA cycle to modulate blood development. An understanding of how hematopoietic tissues leverage TCA activity for proliferation and differentiation while avoiding the excess oxidative burden that disrupts homeostasis, this optimization *in vivo* remains poorly explored.

*Drosophila* larval lymph gland (LG), is the primary hematopoietic organ and site for definitive hematopoiesis. The organ contains multipotent stem-like blood progenitor cells, that reside in the inner Medullary Zone (MZ). As these progenitors transition into differentiation, the transitioning differentiating progenitors are spatially localized in the Intermediate Zone (IZ) of the LG (46). Once matured, the differentiated cells reside in the outer Cortical Zone (CZ) (47, 48). The myeloid-like blood progenitor cells of *Drosophila* LG, differentiate into 3 mature lineages, plasmatocytes, crystal cells and lamellocytes (49–56). Plasmatocytes, are phagocytic and akin to macrophages (57–60); crystal cells, which mediate melanization and wound healing (61–65); and lamellocytes, which are stress-induced immune cells responsible for encapsulating large pathogens such as parasitic wasp eggs(66, 67). Thus the overall spatial organization of blood development in the LG, mirrors that of the vertebrate, from hematopoietic stem cell (HSC) to (common myeloid progenitor (CMP) to myeloid differentiation continuum.

MZ cells are multipotent, stem-like progenitors that, like vertebrate CMPs, maintain elevated Reactive oxygen species (ROS) buffered by high antioxidant capacity (42), restrained mitochondrial activity(68, 69), and metabolite-sensitive identity circuits (70). As cells transition toward the IZ and ultimately the CZ, they progressively increase mitochondrial activity and oxidative metabolism, recapitulating the metabolic intensification characteristic of vertebrate myeloid differentiation (68, 71–77). This architecture alludes to spatially resolved map of metabolic states across blood formation. The metabolic sensitivity of the progenitor cells to sense and responds strongly to systemic, nutritional, and sensory cues further accentuates the strong reliance of metabolic inputs in governing progenitor development and homeostasis. Especially, our recent work has shown the importance of GABA-shunt activity in progenitor cells and that it restricts TCA flux. This is necessary to promote lymph gland growth (68) and maintain anti-oxidant synthesis (69). The TCA enzyme succinate dehydrogenase (SDH/Complex II) produces ROS that primes progenitors for differentiation (42), yet excessive TCA-derived ROS limits growth, revealing a narrow physiological window for TCA-driven redox (68, 69). Restraining *Pdh* activity gates pyruvate entry into the TCA cycle and regulates growth and differentiation onset. These studies collectively demonstrate that TCA activity is essential for blood development, yet both insufficient and excessive flux disrupt hematopoietic balance, underscoring that TCA regulation must be developmentally, spatially, and systemically optimized.

In this study, we systematically interrogate the TCA cycle across the spatial and temporal landscape of *Drosophila* hematopoiesis, perturbing each enzymatic node within defined blood progenitor and differentiating populations. This framework allows us to resolve how individual TCA segments contribute to progenitor maintenance, proliferative expansion, and differentiation, and to determine whether the cycle operates in modular or cyclic configurations during development. Our findings reveal a developmentally programmed transition, from early, functionally discrete citrate and α-KG producing modules to a later, fully integrated oxidative loop, that optimizes metabolic output necessary to preserve progenitor homeostasis and overall differentiation. This modular to cyclic reorganization within the zones, provides a metabolic architecture that simultaneously supports lymph gland growth and preserves homeostatic blood formation. The transitioning TCA topology across different zones in the LG, illuminates the systems changing developmental hematopoietic demands, thereby shaping overall blood development. Overall, the work demonstrates that hematopoiesis relies not on fixed metabolic circuits, but on dynamic reconfiguration of TCA to coordinate distinct phases of hematopoiesis: from proliferation to balanced lineage output at an organ level control.

## Results

### Progenitor compartment: TCA activity sustains progenitor maintenance and lymph gland growth

To define how the tricarboxylic acid (TCA) cycle contributes to blood progenitor development, we systematically perturbed individual TCA enzymes in the *Drosophila* larval lymph gland (LG) using the GAL4-UAS system (Fig. 1A). The lymph gland is organized into distinct hematopoietic zones, each representing a progressive stage of blood differentiation (Fig. 1B). The medullary zone (MZ) at the core harbors the multipotent stem-like blood progenitors, the intermediate zone (IZ) contains transitioning cells that co-express progenitor and early differentiation markers, while the outer cortical zone (CZ) is composed of terminally differentiated hemocytes. Within the MZ, a smaller population of core progenitors marked by *Tep4* expression defines the most stem-like subset (78, 79). To selectively modulate these distinct cell populations, we used a set of spatially restricted/overlapping GAL4 drivers: *domeMESO-GAL4,UAS-GFP* (pan-MZ progenitors), *Tep4-GAL4,UAS-mCherry* (core progenitors), *CHIZ-GAL4* (IZ), and *HmlΔ-GAL4,UAS-2xEGFP* (CZ) (Fig. 1B). These drivers do not label the PSC niche (51, 80–84), allowing zone-specific modulation of metabolic genes without confounding niche-derived effects. Changes in hematopoietic composition were visualized by the reporter fluorescence and assessed by immunostaining for lineage markers: Peroxidasin, *Hemolectin*, for differentiating blood cells; NimC1 (P1) for plasmatocytes; ProPOA for crystal cells; and Myospheroid (Mys) for lamellocytes.

**Figure 1:**
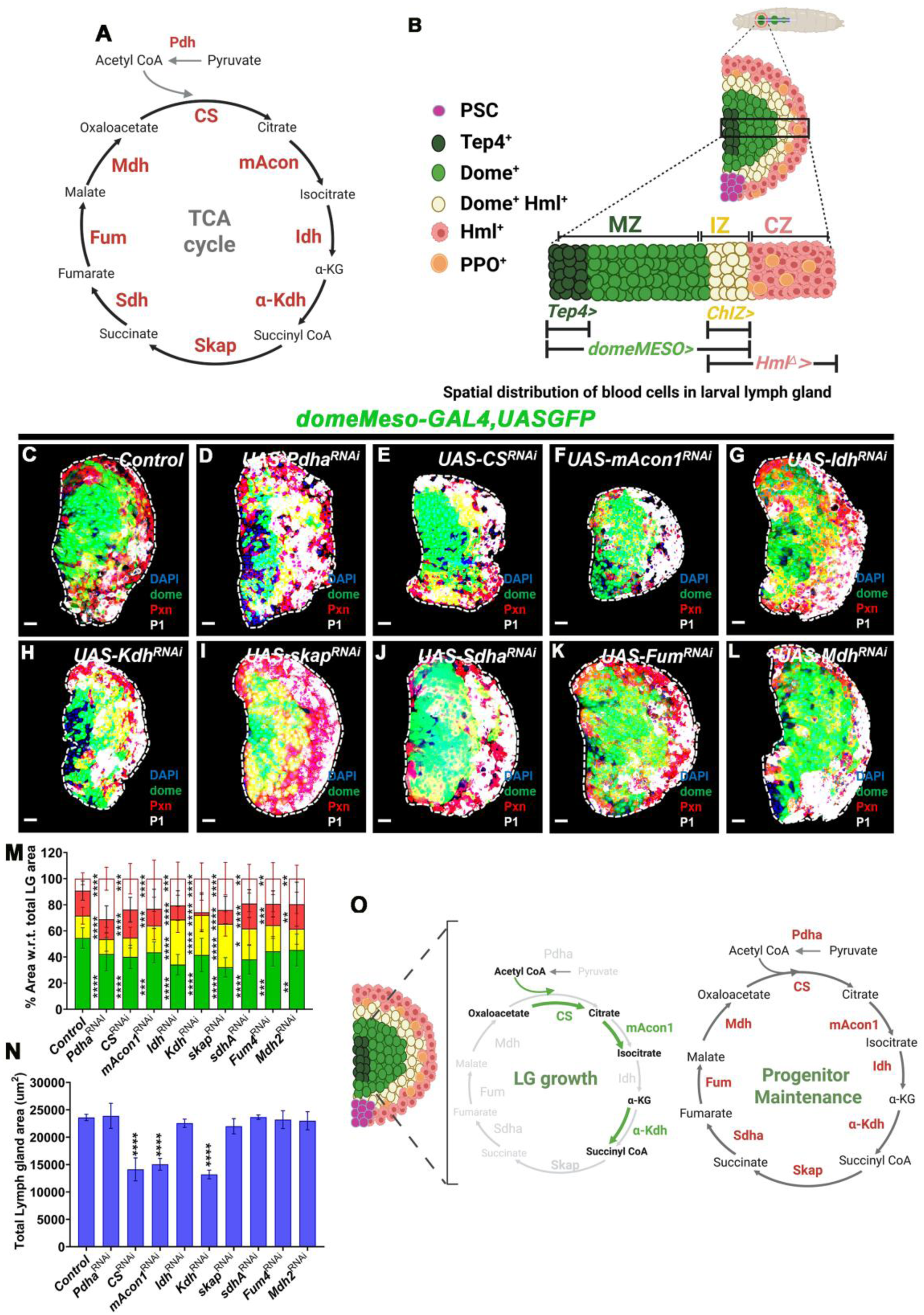
Enzymes of the TCA cycle control medullary zone blood progenitor homeostasis and lymph gland growth. **A)** Schematic representation of the TCA cycle depicting TCA enzymatic steps and respective metabolic conversions involved. **(B)** Diagrammatic Illustration of a 3^rd^ instar *Drosophila* larval lymph gland (top) and the inner spatial organization within each primary lobe of the hematopoietic (bottom). The primary lobe is divided into distinct regions along the medio-lateral axis: The Medullary Zone (MZ, dark and light green), containing hematopoietic progenitors, that are core progenitors (Tep4^+^; dark green) and distal progenitors (Dome^+^; light green); the Intermediate Zone (IZ, yellow), containing partially differentiated cells (Dome^+^Hml^+^; yellow); and the Cortical Zone (CZ, red), composed of differentiating plasmatocytes (Hml^+^; red), and crystal cells (PPO^+^; orange). The Posterior Signaling Center (PSC, magenta) provides niche signals regulating progenitor maintenance. The transgenic drivers used to modulate each respective zone are: *Tep4>* (*C*ore progenitor driver), *domeMESO>* (pan progenitor driver)*, ChIZ>* (intermediate progenitor driver), and *Hml>* (differentiating cell driver). **(C-L)** Representative images of wandering 3^rd^ instar *Drosophila* larval LG marked for respective zones (MZ: Dome^+^, green, IZ: Dome^+^Pxn^+^, yellow, CZ: Pxn^+^, red and P1^+^, white) from **(C)** control (domeMeso-Gal4,UAS-GFP/+), **(D)** *Pdha^RNAi^* (*domeMeso-Gal4,UAS-GFP;UAS-Pdha^RNAi^*), **(E)** *CS^RNAi^ (domeMeso-Gal4,UAS-GFP;UAS-CS^RNAi^*), **(F)** *mAcon1^RNAi^ (domeMeso-Gal4,UAS-GFP;UAS-mAcon1^RNAi^*), **(G)** *Idh^RNAi^* (*domeMeso-Gal4,UAS-GFP;UAS-Idh^RNAi^*), **(H)** *kdh^RNAi^ (domeMeso-Gal4,UAS-GFP;UAS-kdh^RNAi^*), **(I)** *skap^RNAi^ (domeMeso-Gal4,UAS-GFP;UAS-skap^RNAi^*), **(J)** *Sdh^RNAi^ (domeMeso-Gal4,UAS-GFP;UAS-Sdh^RNAi^*), **(K)** *Fum^RNAi^ (domeMeso-Gal4,UAS-GFP;UAS-Fum^RNAi^*), and **(L)** *Mdh^RNAi^ (domeMeso-Gal4,UAS-GFP;UAS-Mdh^RNAi^*), showing reduction in MZ (green) area with a concomitant increase in differentiation population (red, yellow and white) upon loss of TCA enzymatic activity. Compared to **(C)** control. **(E-F** and **H)** also show smaller LG size. **(M)** Quantification of respective zones as relative percent area with respect to total lymph gland area: *domeMeso>GFP/+* (control, n=37), *domeMeso>GFP/Pdha^RNAi^* (n=13, MZ: p<0.0001; IZ: p=0.9488; CZ: Pxn^+^ area p=0.0077, P1 area: p<0.0001), *domeMeso>GFP/CS^RNAi^* (n=16, MZ: p<0.0001; IZ: p=0.9847; CZ: p=0.0077, P1 area: p=0.0004), *domeMeso>GFP/mAcon1^RNAi^* (n=27, MZ: p=0.0002; IZ: p=0.7423; CZ: p>0.9999, P1 area: p<0.0001), *domeMeso>GFP/Idh^RNAi^* (n=35, MZ: p<0.0001; IZ: p<0.0001; CZ: p=0.9927, P1 area: p=0.0004), *domeMeso>GFP/Kdh^RNAi^* (n=35, MZ: p<0.0001; IZ: p<0.0001; CZ: p>0.9999, P1 area: p<0.0001), *domeMeso>GFP/skap^RNAi^*(n=51, MZ: p<0.0001; IZ: p<0.0001; CZ: p=0.5696, P1 area: p<0.0001), *domeMeso>GFP/Sdh^RNAi^*(n=35, MZ: p<0.0001; IZ: p=0.0486; CZ: p=0.1149, P1 area: p=0.0036), *domeMeso>GFP/Fum^RNAi^*(n=42, MZ: p=0.0001; IZ: p=0.7746; CZ: p=0.2021, P1 area: p=0.0014), *domeMeso>GFP/Mdh^RNAi^* (n=39, MZ: p=0.0009; IZ: p>0.9999; CZ: p=0.5932, P1 area: p=0.0013), **(N)** Quantifications of total lymph gland area, *domeMeso>GFP/+* (control, n=79), *domeMeso>GFP/Pdha^RNAi^*(n=32, p=0.9999), *domeMeso>GFP/CS^RNAi^*(n=85, p<0.0001), *domeMeso>GFP/mAcon1^RNA^*^i^ (n=56, p<0.0001), *domeMeso>GFP/Idh^RNAi^*(n=85, p=0.5342), *domeMeso>GFP/Kdh^RNAi^* (n=70, p<0.0001), *domeMeso>GFP/skap^RNAi^* (n=66, p=0.1512), *domeMeso>GFP/Sdh^RNAi^* (n=84, p>0.9999), *domeMeso>GFP/Fum^RNAi^*(n=76, p=0.9973), *domeMeso>GFP/Mdh^RNAi^* (n=58, p=0.9843). (**O**) Model summarizing all the TCA steps involved in lymph gland growth (green arrows in the small circle) and progenitor maintenance (black arrows in the big circle). Data is presented as median plots (*p<0.05; **p<0.01; ***p<0.001; ****p<0.0001), ordinary one-way ANOVA. Scale bar: 20µm. ‘n’=lymph gland lobes. DAPI marks DNA, all comparisons for significance are with respect to control values. LG, is lymph gland. One respective LG lobe from each genetic condition is shown and outlined with white border with all other accessory tissues cleared for better demarcation.

Under homeostatic condition, the third-instar LG displays a well-defined zonal organization, where the progenitor-rich Dome-GFP⁺ MZ region devoid of any differentiation marks, (hence forth referred to as Dome^+^ only) occupies roughly 55% of the total lymph gland area, while the remaining 45% of the area corresponds to the differentiated Pxn⁺ population (Fig. 1C, M). Of the total Pxn⁺ population, they contain 10– 15% intermediate zone (IZ) cells, co-expressing Dome-GFP⁺ marker, 20% Dome-GFP^-^ Pxn+ differentiating blood cells and 10% terminally differentiated P1⁺ plasmatocytes, all located at the outer most CZ region (Fig. 1C, M). Crystal cell numbers are found to be ∼35–40 cells per LG lobe (Fig. S1A1, E). Overall, this spatially organized framework of the LG demonstrates the homeostatic distribution of blood cells from varying stages of development and maturation. Any deviation from the wild type proportion upon modulating the TCA from the respective zones, is what we set out to address.

To test the role of TCA cycle in progenitor development, we targeted the MZ compartment using *domeMESO-GAL4,UAS-GFP* and perturbed all enzymatic steps of the cycle including *Pdh*, the rate-limiting enzyme controlling pyruvate entry into the TCA (85). Across all knockdowns, we observed a striking reduction in the Dome⁺ only progenitor area (Fig. 1C-L and M). We also observed an increase in Pxn^+^ area, which is comprising of the differentiating-IZ, and differentiated-CZ cell populations (Fig. 1C–L, M). The relative proportion of IZ area, which is the Dome⁺Pxn⁺ population, across most enzymatic knock-downs remained largely unchanged with exceptions of *Isocitrate dehydrogenase (Idh)* (Fig. 1G, M)*, α-Ketoglutarate Dehydrogenase (Kdh)* (Fig. 1H, M) and *Succinyl-coenzyme A synthetase (skap)* (Fig. 1I, M), where their loss showed significant expansion of the IZ area. Specifically, across all knock-down conditions, a significant increase in terminally mature P1 blood cell population was evident (Fig. 1C–L, M). On the contrary, there was a reduction in the crystal cell numbers in specific knock-downs of *CS^RNAi^* (Fig S1A2, E), *mAcon1^RNAi^* (Fig S1 A3, E)*, Kdh^RNAi^* (Fig S1A5, E)*, skap^RNAi^* (Fig S1A6, E) and *Sdha^RNAi^*(Fig S1 A7, E) while in the others it remained comparable to controls (Fig. A4, A8, A9 and S1E). Lamellocytes were also detected under all TCA step perturbations, except for *Kdh* knockdown (Table S1). These observations implied that blocking the TCA-cycle in MZ progenitor cells led to loss of progenitor homeostasis and increased differentiation into terminal plasmatocyte lineage with reduction in crystal cell potential. Thus, indicating a role for TCA cycle in progenitor maintenance, homeostasis and differentiation dynamics.

In addition to this broad progenitor-to-differentiation transition, we identified a second, distinct phenotype, which is smaller lymph glands that appeared selectively upon MZ-specific loss of *CS* (Fig. 1E, N), *mAcon1* (Fig. 1F, N), or *Kdh* (Fig. 1H, N). We observed a 30-40% size reduction as assessed by total lymph gland area quantifications (See Methods). This unexpected observation pointed to the requirement of only specific TCA enzymatic nodes in promoting lymph gland growth. The phenotypic characterizations were validated using multiple RNA*i* lines that have been reported previously for their functional roles in other contexts (86). Overall, we observed a consistent reduction in Dome^+^ only area with a corresponding increase in Pxn^+^ population (Fig. S2A) with these lines as well. Defect in growth with *CS^RNAi^,* and *Kdh^RNAi^* (Fig. S2B) was also noticed. The P1 expansion phenotype was however of varying degrees, which alluded to strength of RNA*i* knock-down efficacies (Fig. S2A). Regardless, using multiple RNA*i*, the aforementioned phenotypes of differentiation and growth were largely recapitulated, indicating that the effects were specific and reproducible. The strongest RNA*i* lines were thereafter chosen for further analysis. Altogether, the data revealed dual facets of the TCA where LG growth was independent of the entire TCA cycle’s operation, and relied on only specific steps, while progenitor homeostasis was reliant on its cyclical mode.

To further refine the requirements observed in progenitor cells, we next used the *Tep4-GAL4,UAS-mCherry* driver to target the core progenitor subset (Fig. S2C). Under normal conditions, *Tep4*⁺ core progenitors occupy ∼35-40% of the total LG area, with ∼40% of remaining cells expressing Pxn⁺ differentiation marker and ∼20% representing Tep4⁻Pxn⁻ double-negative cells situated between these two compartments (Fig. S2C, E). Knockdown of TCA enzymes in this subset caused a robust reduction in *Tep4*⁺ progenitor areas but interestingly, this reduction was not accompanied by any Pxn expansion (Fig. S2C and E-M). This was unlike the phenotypes seen with pan-progenitor *Dome>* based knockdowns where along with reduction in progenitor population a concomitant increase in Pxn^+^ cells were also apparent (Fig. 1C-L, M). Notably, the reduction in Tep4^+^ area led to a corresponding expansion of the Tep4⁻Pxn⁻ region across all conditions and implied that loss of TCA activity in the core-progenitor cells led to their reduction in numbers but did not change the overall differentiation status in the lymph gland. Interestingly, here as well we observed growth defects, but restricted to *mAcon1* (Fig. S2D & G) and *Kdh* (Fig. S2D & I) knockdowns, while *CS* depletion in Tep4^+^ subset showed no measurable impact on lymph gland size even though it showed reduction in Tep4^+^ region (Fig. S2D & F). This contrasting data indicated zone specific requirement for CS in core progenitor cells, towards their development and maintenance, without regulating overall LG growth. Crystal cell numbers remained relatively constant at ∼35–40 cells per LG lobe (Fig. SB1, B2, B4, B6-B9 and F), except for *mAcon1^RNAi^*(Fig. S1B3, F), and *Kdh^RNAi^* (Fig. S1B5, F). Lamellocytes were also detected under all TCA perturbations (Table S1). Together, these data revealed a dynamic role of the TCA in core progenitor cells for controlling their numbers and overall lymph gland growth. Maintaining differentiation status of the lymph gland, however emerged independent of modulating the TCA in the core progenitor pool and was more distinctly seen with the larger Dome^+^ progenitor subset (Fig. 1O).

**Figure 2.**
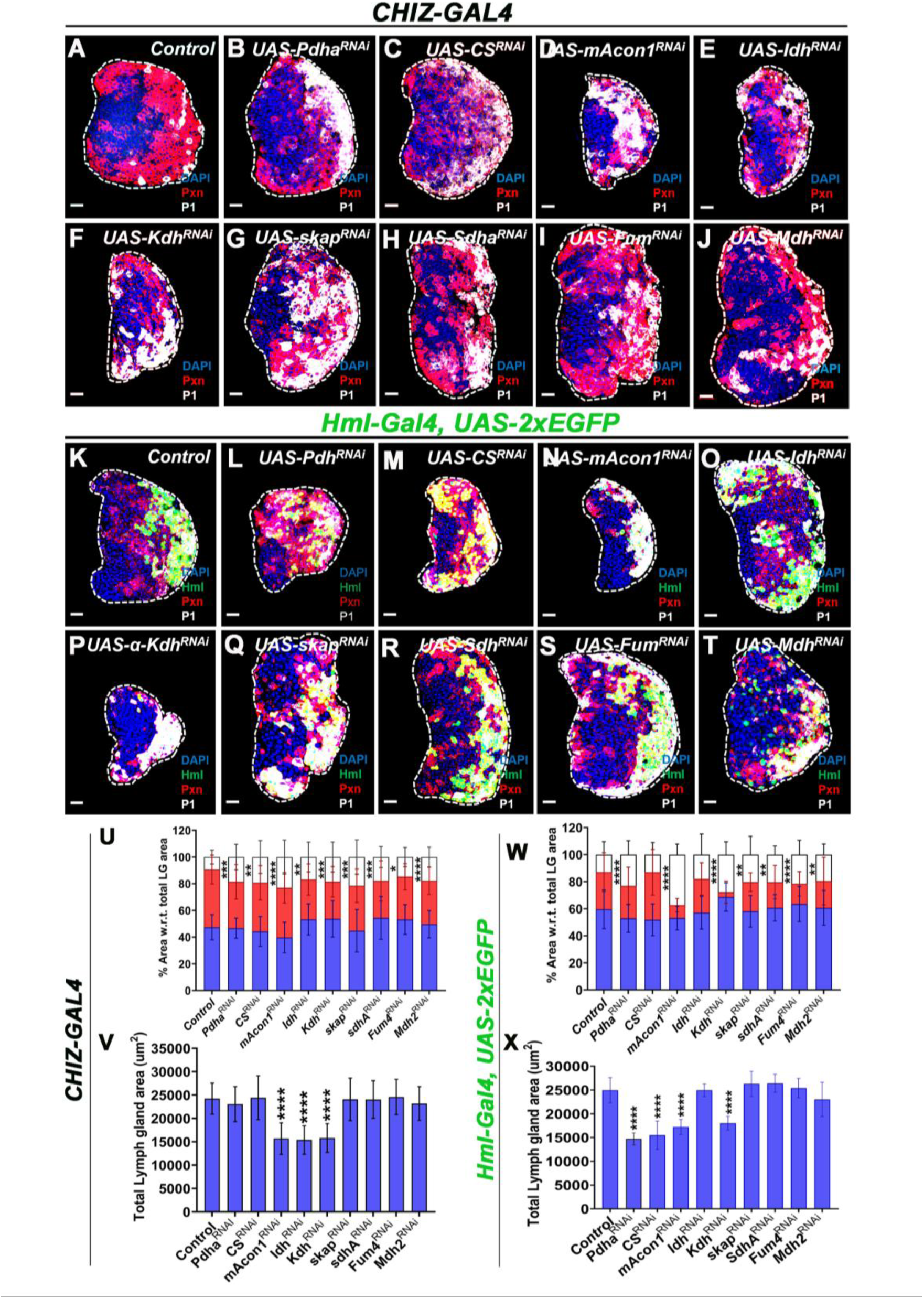
TCA cycle enzymes function in the IZ and CZ region to control overall differentiation homeostasis of the lymph gland and its growth. **(A-J)** Representative images of wandering 3^rd^ instar *Drosophila* larval LG showing differentiating blood cells marked for Pxn (red), plasmatocytes marked with P1 (white), and undifferentiated cells (blue, DAPI stained) from **(A) IZ** control (*ChIZ-GAL4/+)* **(B)** *Pdha^RNAi^ (ChIZ-GAL4;UAS-Pdha^RNAi^),* **(C)** *CS^RNAi^ (ChIZ-GAL4;UAS-CS^RNAi^),* **(D)** *mAcon1^RNAi^ (ChIZ-GAL4;UAS-mAcon1^RNAi^)*, **(E)** *Idh^RNAi^ (ChIZ-GAL4;UAS-Idh^RNAi^), (***F)** *α-Kdh^RNAi^ (ChIZ-GAL4;UAS-α-Kdh^RNAi^),* (**G)** *Skap^RNAi^ (ChIZ-GAL4;UAS-Skap^RNAi^),* (**H)** *Sdh^RNAi^ (ChIZ-GAL4;UAS-Sdh^RNAi^),* **(I)** *Fum^RNAi^ (ChIZ-GAL4;UAS-Fum^RNAi^),* and **(J)** *Mdh^RNAi^ (ChIZ-GAL4;UAS-Mdh^RNAi^)*. Compared to (**A**) *control,* (**B-J**) loss of TCA enzymatic activity in the IZ cells leads to expansion of plasmatocyte lineage (P1^+^white, see quantifications in **U**) and **(D-F)** smaller LGs in (D) *mAcon1^RNAi,^ **(E)** Idh^RNAi^* and (F) *α-Kdh^RNAi^* expressing conditions (see quantifications in **V**)**. (K-T)** Representative images of wandering 3^rd^ instar *Drosophila* larval LG showing differentiating blood cells marked for Hml (green), Pxn (red), plasmatocytes marked with P1 (white), and undifferentiated cells (blue, DAPI stained) from **(K) CZ** control *(Hml*^△^*-GAL4,UAS-2X EGFP/+)* **(L)** *Pdha^RNAi^ (Hml*^△^*-GAL4,UAS-2X EGFP;UAS-Pdha^RNAi^),* **(M)** *CS^RNAi^ (Hml*^△^*-GAL4,UAS-2XEGFP;UAS-CS^RNAi^),* **(N)** *mAcon1^RNAi^ (Hml*^△^*-GAL4,UAS-2XEGFP;UAS-mAcon1^RNAi^),* **(O)** *Idh^RNAi^ (Hml*^△^*-GAL4,UAS-2XEGFP;UAS-Idh^RNAi^),* (**P)** *α-Kdh^RNAi^ (Hml*^△^*-GAL4,UAS-2XEGFP;UAS-α-Kdh^RNAi^),* (**Q)** *Skap^RNAi^ (Hml*^△^*-GAL4,UAS-2XEGFP;UAS-Skap^RNAi^),* (**R)** *Sdh^RNAi^ (Hml*^△^*-GAL4,UAS-2XEGFP;UAS-Sdh^RNAi^),* **(S)** *Fum^RNAi^ (Hml*^△^*-GAL4,UAS-2XEGFP;UAS-Fum^RNAi^)*, and **(T)** *Mdh^RNAi^ (Hml*^△^*-GAL4,UAS-2XEGFP;UAS-Mdh^RNAi^)*. Compared to (**K**) CZ *control*, specific loss of (L) *Pdh^RNAi^***(N),** m*Acon1^RNAi^* **(P)***Kdh^RNAi^* **(Q)** *Skap^RNAi^* **(R)** *Sdh^RNAi^* **(S)** *Fum^RNAi^* and **(T)** *Mdh^RNAi^* enzymatic activities in the CZ cells leads to expansion of plasmatocyte lineage (white, see quantifications in **W**) and small lymph in (L) *Pdh^RNAi^*(M) *CS^RNAi^* (N*) mAcon1^RNAi^* and (P) *Kdh^RNAi^*. **(U)** Quantifications of respective LG areas marked with Pxn (red bar) and P1 (white bar) as relative percent area with respect to total lymph gland area: *ChIZ-GAL4/+* (*control,* n=40), *ChIZ-GAL4/Pdha^RNAi^* (n=23, blue: p>0.9999; red: p>0.9999; white: p=0.0026), *ChIZ-GAL4/CS^RNAi^* (n=23, blue: p=0.9240; red: p=0.9240; white: p=0.0009), *ChIZ-GAL4*/mAcon1^RNAi^ (n=26, blue: p=0.0766; red: p=0.0766; white: p<0.0001), *ChIZ-GAL4/Idh^RNAi^* (n=39, blue: p=0.1824; red: p=0.1824; white: p=0.0051), *ChIZ-GAL4/α-Kdh^RNAi^* (n=31, blue: p=0.1755; red: p=0.1755; white: p=0.0008),*ChIZ-GAL4/Skap^RNAi^* (n=18, blue: p=0.9837; red: p=0.9837; white: p=0.0001), *ChIZ-GAL4/Sdh^RNAi^* (n=30, blue: p=0.1063; red: p=0.1063; white: p=0.0025), *ChIZ-GAL4/Fum^RNAi^* (n=40, blue: p=0.1825; red: p=0.1825; white: p=0.0929), *ChIZ-GAL4/Mdh^RNAi^* (n=40, blue: p=0.9706; red: p=0.9706; white: p=0.0009), **(V)** Quantifications of total lymph gland areas, *ChIZ-GAL4/+* (control, n=61), *ChIZ-GAL4/Pdha^RNAi^*(n=49, p=0.4669), *ChIZ-GAL4/CS^RNAi^*(n=48, p>0.9999), *ChIZ-GAL4/mAcon1^RNAi^*(n=63, p<0.0001), *ChIZ-GAL4/Idh^RNAi^* (n=64, p<0.0001), *ChIZ-GAL4/Α-Kdh^RNAi^* (n=60, p<0.0001), *ChIZ-GAL4/Skap^RNAi^*(n=31, p>0.9999), *ChIZ-GAL4/Sdh^RNAi^*(n=51, p>0.9999), *ChIZ-GAL4/Fum^RNAi^* (n=40, p=0.9996), *ChIZ-GAL4/Mdh^RNAi^* (n=48, p=0.6187). **(W)** Quantifications of respective LG areas marked with Pxn (red bar) and P1 (white bar) as relative percent area with respect to total lymph gland area: *Hml*^△^*-GAL4,UAS-2X EGFP/+* (control, n=38), *Hml*^△^*-GAL4,UAS-2X EGFP/Pdha^RNAi^* (n=25, blue, p=0.2110; red, p=0.2110; white, p<0.0001), *Hml*^△^*-GAL4,UAS-2X EGFP/CS^RNAi^* (n=21, blue, p=0.1297; red, p=0.1297; white, p=0.7738), *Hml*^△^*-GAL4,UAS-2X EGFP/mAcon1^RNAi^* (n=14, blue, p=0.4810; red, p=0.4810; white, p<0.0001), *Hml*^△^*-GAL4,UAS-2X EGFP/Idh^RNAi^* (n=25, blue, p=0.9839; red, p=0.9839; white, p=0.0112), *Hml*^△^*-GAL4,UAS-2X EGFP/α-Kdh^RNAi^* (n=23, blue, p=0.0433; red, p=0.0433; white, p<0.0001), *Hml*^△^*-GAL4,UAS-2XEGFP/skap^RNAi^* (n=12, blue, p>0.9999; red, p>0.9999; white, p=0.0011), *Hml*^△^*-GAL4,UAS-2XEGFP/Sdha^RNAi^* (n=13, blue, p>0.9999; red, p>0.9999; white, p=0.0024), *Hml*^△^*-GAL4,UAS-2XEGFP/Fum^RNAi^*(n=29, blue, p=0.7887; red, p=0.7887; white, p<0.0001), *Hml*^△^*-GAL4,UAS-2XEGFP/Mdh^RNAi^* (n=20, blue, p>0.9999; red, p>0.9999; white, p=0.0034). **(X)** Quantifications of total lymph gland areas, *Hml*^△^*-GAL4,UAS-2XEGFP*/+ (*control*, n=63), *Hml*^△^*-GAL4,UAS-2XEGFP/Pdha^RNAi^* (n=28, p<0.0001), *Hml*^△^*-GAL4,UAS-2XEGFP/CS^RNAi^* (n=61, p<0.0001), *Hml*^△^*-GAL4,UAS-2X EGFP/mAcon1^RNAi^* (n=48, p<0.0001), *Hml*^△^*-GAL4,UAS-2XEGFP/Idh^RNAi^* (n=47, p=0.6870), *Hml*^△^*-GAL4,UAS-2X EGFP/α-Kdh^RNAi^* (n=30, p<0.0001), *Hml*△*-GAL4,UAS-2X EGFP/skap^RNAi^* (n=38, p=0.9998), *Hml*^△^*-GAL4,UAS-2XEGFP/Sdha^RNAi^*(n=53, p=0.9987), *Hml*^△^*-GAL4,UAS-2XEGFP/Fum^RNAi^* (n=45, p=0.9769), *Hml*^△^*-GAL4,UAS-2XEGFP /Mdh^RNAi^* (n=47, p=0.0743). Data is presented as bar plots. (*p<0.05; **p<0.01; p<0.001; ****p<0.0001), ordinary one-way ANOVA. Scale bar: 20µm. ‘n’=lymph gland lobes. DAPI marks DNA, differentiating population (Pxn+; Red), terminally differentiated population (P1+; white). Comparisons for significance are with respect to control values. LG lobes are outlined with white border along with removal of other accessory tissues for clear demarcation.

Thus, the perturbations across both the drivers revealed two key features of the TCA enzymes. First, TCA activity in the larger quorum of Dome^+^ progenitor cells are essential for maintaining overall progenitor homeostasis and differentiation status in the lymph gland. This includes flux through *Pdh*a, as loss of flux via *Pdha* consistently depleted progenitor cells and led to the expansion of P1 population (Fig. 1D & M). Second, specific enzymatic nodes *CS*, *Aconitase*, and *Kdh* that favored growth, alluded to TCA functioning in a modular manner to accommodate growth. This is distinct from its functioning as a cyclical loop in governing progenitor homeostasis. These results suggested that the TCA cycle, normally known for its functioning as a bioenergetics hub (87–89), in the progenitors operated dynamically to couple distinct metabolic fluxes to enable progenitor development, maintenance and lymph gland growth (Fig. 10).

### Intermediate and cortical zones show spatially distinct enzymatic requirements

To determine whether TCA activity in the downstream differentiating hematopoietic compartments regulated any aspect of blood development, we modulated TCA enzymes in the intermediary IZ progenitor cells and the differentiating blood cells of the CZ using *CHIZ-GAL4* (IZ), and *Hml^Δ^-GAL4,UAS-2xEGFP* (CZ) respectively. In these manipulations, the respective proportion of progenitor cell area was assessed by analyzing inner-core regions of the lymph gland devoid of any Pxn expression (Pxn⁻, respective progenitor area) and regions marked by Pxn⁺ expression, marking differentiating cell areas respectively. We also stained for P1 to mark the terminally differentiated cells as done with progenitor-based modulations.

Intriguingly, down-regulating TCA cycle enzymes in the IZ showed no difference in Pxn^-^ or differentiating cell Pxn^+^ areas (Fig 2A-J and U). However, we observed a significant expansion of terminally differentiated P1⁺ population across all knockdowns, where we observed almost double the number of P1^+^ cells (Fig. 2A–J, U). Thus, indicating that TCA activity in IZ cells non-autonomously restrained expression of premature terminal differentiation markers in the CZ region. Growth retardation was also observed here, but upon knockdown of *mAcon1* (Fig. 2D & V), *Idh* (Fig. 2E & V), and *Kdh* (Fig. 2F & V), whose down-regulation led to marked reduction in lymph gland size. *CS* knockdown in the IZ cells did not show any change in lymph gland size (Fig. 2C & V). Crystal cell numbers while it remained unchanged in most conditions (Fig. S1C2, C6-C9 and G), it was reduced in *mAcon1^RNAi^*(Fig. S1C3 & G)*, Idh^RNAi^* (Fig. S1C4 & G) *and Kdh^RNAi^* (Fig. S1C5 & G) backgrounds. Intriguingly, perturbation of the TCA enzymes in the IZ also led to increased lamellocytes formation (Table S1). This alluded to regulation of differentiation into L1 and P1 subtype, by TCA activity in the IZ compartment.

In the CZ, *Hml^Δ^-GAL4,UAS-2xEGFP* driven knockdowns, it did not alter the overall Pxn^+^ area (Fig. 2K–T & W) or Hml⁺ area (Fig. S3A). Pxn^-^, prospective progenitor region also remained unaffected (Fig. 2W). Nevertheless, the knockdowns of TCA enzymes, excluding *CS* and *Idh*, led to a consistent increase in P1 expression (Fig. 2K-T & W). This suggested that TCA activity in mature hemocytes is largely dispensable for maintaining CZ proportions but controlled the extent of terminal differentiation. Additionally, like seen with the rest of the driver lines, a reduced gland size was also evident, and intriguingly now upon loss of *Pdha* (Fig. 2L & X)*, CS* (Fig. 2M & X)*, mAcon1* (Fig. 2N & X), and *Kdh* (Fig. 2P & X). The data revealed independence of the CZ cells from any *Idh* requirement in either regulating differentiation (Fig 2O & W) or growth (Fig. 2O and X). This observation is unlike seen in *CHIZ* cells, where *Idh* function controlled both P1 homeostasis (Fig 2E and U) and growth (Fig. 2E, and V). Crystal cell numbers were compromised in *CS^RNAi^* (Fig. S1D2 & H) *mAcon1^RNAi^* (Fig. S1D3 & H)*, and Kdh^RNAi^*(Fig. S1D5 & H), also lamellocytes were detected in all TCA knockdowns except for *Idh* and *skap* loss of function (Table S1). Overall, the data here allude to spatially distinct activity profiles of teshe TCA cycle between the IZ and differentiated CZ compartment.

### Modular architecture of the TCA cycle

Especially with respect to lymph gland growth, in the MZ progenitor and also CZ, the independence of *Idh* (Fig. 1N, 2X) in these zones, whose activity connects citrate to a-KG metabolism (Fig. 1A), revealed a disconnect in α-KG production from citrate. This establishes two nodes: a *CS-mAcon1* branch that governs citrate metabolism, while an *α-Kdh* module that controls α-ketoglutarate metabolism (Fig. 1O and S3N). Thus, to address additional routes that would facilitate α-KG production in MZ and CZ cells, we tested Glutamate dehydrogenase (*Gdh*) function in blood development. *Gdh* converts glutamate to α-KG (12, 13, 90), which prompted us to test if this mode facilitated a-KG production in blood cells (Fig. S3B). Supporting this notion, knockdown of *Gdh*, phenocopied *α-Kdh* loss in MZ progenitors leading to small lymph glands (Fig. S3C, D and G), loss of Dome⁺ progenitors, with expansion of Pxn⁺ and P1⁺ cells (Fig. S3H), and reduced crystal cell numbers (Fig. S3E, F and I). This implied *Gdh* function in the MZ facilitated its *α-*KG requirement (Fig. S3J).

However, *Gdh* knockdown in the CZ did not affect lymph gland size (Fig. S3K-M), but did lead to expansion of P1 expression (Fig. S3L and N). This confirmed that α-KG generation in differentiating cells involved *Gdh* activity, however unlike *Gdh’s* role in MZ (Fig. S3D), in the CZ it did not contribute to growth but was important for differentiation homeostasis. This revealed that the TCA in the CZ was disconnected as it functioned independently of *Idh,* but involved a modular operation via *Gdh* to control differentiation homeostasis (Fig. S3O).

Thus, within the major compartments of MZ and CZ, *CS-mAcon1* module and *Kdh* activity emerged as the dominant growth-linked enzymes, and was disengaged from the requirement for *Idh* to support growth (Fig. S3J and O). In contrast, in the differentiating IZ, *mAcon1*-*Idh-Kdh* operated as a continuous route to support LG growth (Fig. S3O).

### Spatial and Temporal Distribution of TCA Cycle Enzyme Transcripts in the Lymph Gland

Based on the dynamic nature of TCA activity detected in the genetic analysis, we set out to assess the developmental expression levels of the TCA enzymes. To do this, we investigated the spatial and temporal distribution of TCA cycle enzyme transcripts across different stages of lymph gland development by employing SABER-FISH (Signal Amplification By Exchange Reaction Fluorescence In Situ Hybridization) technique. This technique enables high-sensitivity detection of individual mRNA transcripts. Through the use of Primer Exchange Reaction (PER)-amplified concatemer probes, this allows visualization at single-cell resolution [60]. For each gene, probe sets targeting 35–60 unique sequences were designed. Methodological validation was performed using established lymph gland marker, *Hemolectin (Hml)* (differentiating cell specific). *Hml* transcripts labelled with two spectrally distinct fluorophores (488 nm and 546 nm), as expected, localized predominantly to the lymph gland periphery, consistent with its role as a cortical zone (CZ) marker. The two spectral green and red probes showed over 95% signal overlap, confirming the assay’s specificity and low rate of false-positive detection (Fig. S4A–C3).

Following validation, SABER-FISH was used to analyse transcript dynamics of key TCA cycle enzymes at 48, 72, and 96 hours after egg laying (AEL). Importantly, all TCA enzymes were seen expressed at all developmental time points, albeit at varying expression levels. We observed *Pdh* transcripts uniformly distributed across all LG cells but at lower levels (Fig. S5A–A3). Contrary to *Pdh*, *CS mRNA* expression was higher and detected ubiquitously from 48 to 72h (Fig. S5B–B1). However, at 96h, it exhibited a marked enrichment in the CZ region, but was significantly reduced in the core region (Fig. S5B2–B3). *mAcon1* and *Kdh* mRNA expression were predominantly higher in the core region across all time points (Fig. S5C–D3), with highest levels detected at 48h (Fig. S5C, D). *Gdh* expression was also confined to the inner core area throughout development (Fig. S5E–E3), with elevated levels seen at later time point (Fig. S5E2). Conversely, *Sdh* exhibited consistent expression with time (Fig. S5F–F3), with later at 96h is seen enriched in outer CZ cells (Fig. S5F2-F3). *Mdh* showed weak but uniform expression across all zones and at all-time points (Fig. S5G–G3).

These results demonstrated that, at the transcript level, most TCA cycle enzymes are expressed throughout lymph gland development, with distinct spatiotemporal expression patterns. A clear enrichment of most of the TCA enzymes in the core area was evident which corroborated with the strongest requirement in the progenitor cells.

Altogether, our genetic findings alongside the resolution of expression pattern of the respective TCA enzymes, revealed that the TCA cycle operated in a spatially resolved manner during hematopoiesis (Fig. S3J and O). While complete TCA flux preserves progenitor identity and differentiation homeostasis across zones, specific enzymatic nodes couple metabolism to lymph gland growth. This includes the requirement of *CS/mAcon1* and *α-Kdh/Gdh* modules for progenitor expansion and lymph gland growth.

The selective independence of *Idh* in MZ and CZ, outside the IZ, suggests that the TCA enzymes do not act in a uniform metabolic structure. Furthermore, the non-overlapping nature of the growth phenotypes across all driver lines support a model where TCA is flexibly assembled in the zones and zone-specific modules support the proliferative programs of the respective blood populations. With respect to differentiation, notably, the IZ population encompassing overlapping progenitor and differentiating cells, their manipulation non-autonomously restrains CZ differentiation (Fig. S3O). In contrast, CZ cells in the absence of any *Idh* function in this compartment, they appear to operate primarily with a disconnected TCA in them (Fig. S3O). These observations reveal that TCA transitions between being modular to circular within Dome^+^ progenitors and IZ cells. This modality coordinates overall lymph gland growth with homeostatic differentiation, while in the CZ, it is modular that supports both growth and also homeostatic differentiation.

### Modular TCA activity drives early hematopoietic progenitor development

Our discovery that specific enzymatic nodes of the TCA cycle, notably, *CS* (Fig. 1E &N), *mAcon1* (Fig. 1F &N), *α-Kdh*, (Fig. 1H &N), and *Gdh* (Fig. S3C-D, & G), that belong to the oxidative arm of the TCA cycle (69, 75, 91, 92) and govern lymph gland growth, in contrast to others that constitute the reductive branch (93–95), this modular rather than closed configuration, prompted us to understand how these modules contributed to early hematopoietic progenitor development.

Lymph gland growth under homeostatic conditions follows a biphasic trajectory (51, 82). Early burst of progenitor proliferation establishes the medullary zone (MZ) (Fig. 3A, B), followed by their transition in G2 arrest and slowed proliferation. At this later phase, the second proliferative wave is seen in the differentiating blood cells of the LG that establish the CZ (Fig. 3C). Together, the two proliferative waves in the respective blood cell population contribute to the overall growth of the lymph gland. Thus, we examined the roles of these enzymes in the temporal and spatial control of progenitor proliferation, maintenance, and differentiation. For comparison, we also analyzed loss-of-function conditions for *Sdh* and *Mdh*, enzymes of the reductive arm, which did not show overt growth defects but still contributed to progenitor homeostasis and P1 balance.

**Figure 3:**
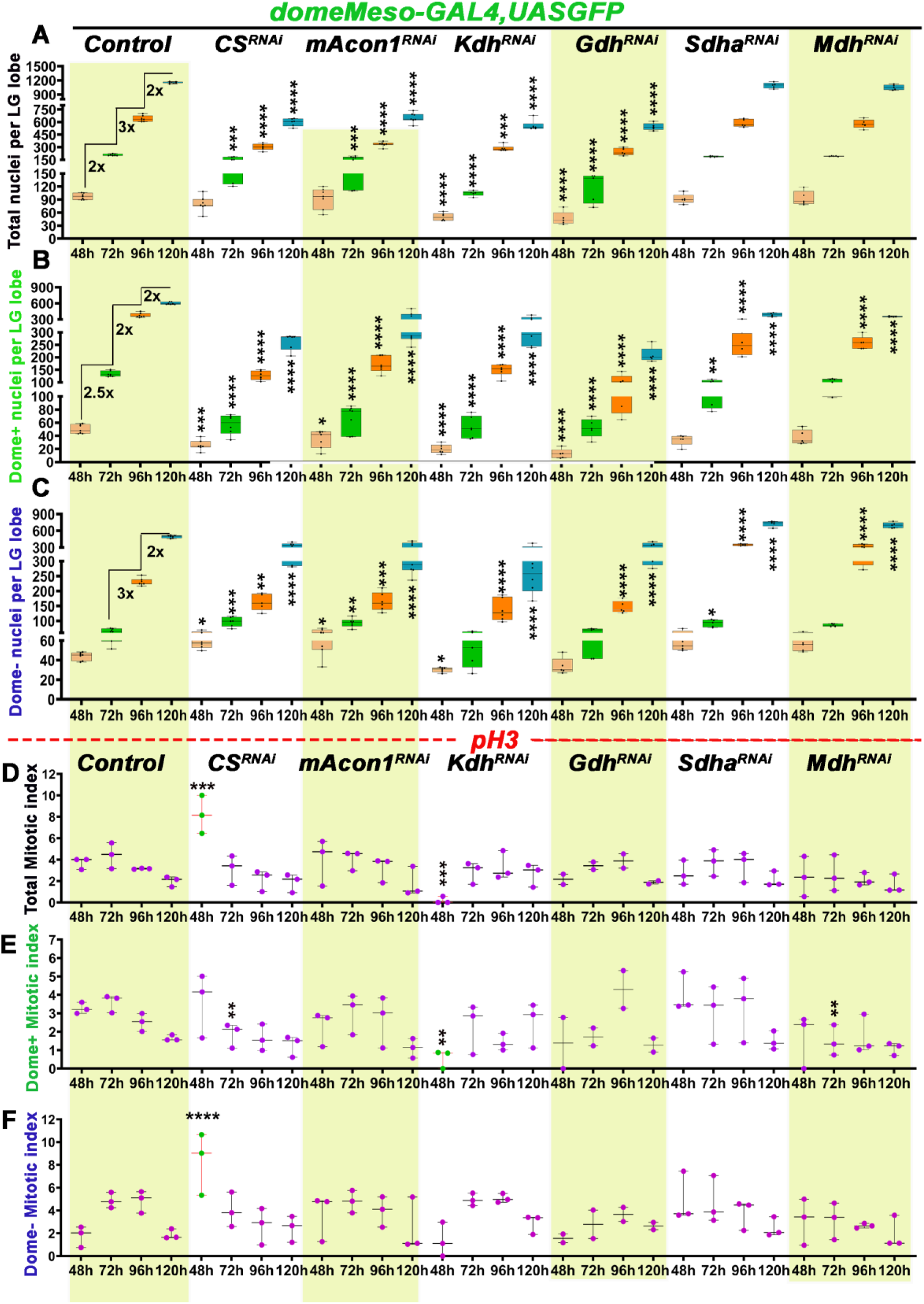
Temporal growth dynamics of a developing lymph gland. **(A)** Graph showing quantification of the total number of cells (based on nuclei count) per lymph gland lobe (LG) over developmental time. *domeMeso>GFP/+* (*control*, 48h, n=75; 72h, n=73, p<0.0001; 96h, n=53, p<0.0001; 120h, n=53, p<0.0001), *domeMeso>GFP/CS^RNAi^* (48h, n=54, p=0.1976; 72h, n=50, p=0.0006; 96h, n=55, p<0.0001; 120h, n=41, p<0.0001), *domeMeso>GFP/mAcon1^RNAi^*(48h, n=69, p=0.9904; 72h, n=69, p=0.0009; 96h, n=65, p<0.0001; 120h, n=66, p<0.0001), *domeMeso>GFP/α-Kdh^RNAi^* (48h, n=37, p<0.0001; 72h, n=48, p<0.0001; 96h, n=63, p<0.0001; 120h, n=49, p<0.0001), and *domeMeso>GFP/Gdh^RNAi^*(48h, n=34, p<0.0001; 72h, n=45, p<0.0001; 96h, n=39, p<0.0001; 120h, n=55, p<0.0001), *domeMeso>GFP/Sdha^RNAi^*(48h, n=38, p=0.9775; 72h, n=49, p=0.4901; 96h, n=43, p=0.1051; 120h, n=39, p=0.2029), *domeMeso>GFP/Mdh^RNAi^* (48h, n=37, p=0.9950; 72h, n=50, p=0.7326; 96h, n=56, p=0.0483; 120h, n=41, p0.0087), (**B)** Graph showing quantification for the number of progenitor cells (Dome⁺, green) per lymph gland (LG) at successive developmental time points. *domeMeso>GFP/+* (control, 48h, n=75; 72h, n=73, p<0.0001; 96h, n=53, p<0.0001; 120h, n=53, p<0.0001), *domeMeso>GFP/CS^RNAi^* (48h, n=54, p=0.0001; 72h, n=50, p<0.0001; 96h, n=55, p<0.0001; 120h, n=41, p<0.0001), *domeMeso>GFP/mAcon1^RNAi^*(48h, n=69, p=0.0205; 72h, n=69, p<0.0001; 96h, n=65, p<0.0001; 120h, n=66, p<0.0001), *domeMeso>GFP/α-Kdh^RNAi^* (48h, n=37, p<0.0001; 72h, n=48, p<0.0001; 96h, n=63, p<0.0001; 120h, n=49, p<0.0001), and *domeMeso>GFP/Gdh^RNAi^* (48h, n=34, p<0.0001; 72h, n=45, p<0.0001; 96h, n=39, p<0.0001; 120h, n=55, p<0.0001), *domeMeso>GFP/Sdha^RNAi^* (48h, n=38, p=0.0220; 72h, n=49, p=0.0011; 96h, n=43, p<0.0001; 120h, n=39, p<0.0001), *domeMeso>GFP/Mdh^RNAi^* (48h, n=37, p=0.1261; 72h, n=50, p=0.0287; 96h, n=56, p<0.0001; 120h, n=41, p<0.0001), **(C)** Graph showing quantification for the number of non-progenitor cells (Dome⁻, blue) per lymph gland (LG) across developmental time. *domeMeso>GFP/+* (control, 48h, n=75; 72h, n=73, p=0.0269; 96h, n=53, p<0.0001; 12h, n=53, p<0.0001), *domeMeso>GFP/CS^RNAi^* (48h, n=54, p=0.0234; 72h, n=50, p=0.0008; 96h, n=55, p=0.0011; 120h, n=41, p<0.0001), *domeMeso>GFP/mAcon1^RNAi^*(48h, n=69, p=0.0191; 72h, n=69, p=0.0030; 96h, n=65, p=0.0008; 120h, n=66, p<0.0001), *domeMeso>GFP/α-Kdh^RNAi^* (48h, n=37, p=0.0248; 72h, n=48, p=0.1963; 96h, n=63, p<0.0001; 120h, n=49, p<0.0001), and *domeMeso>GFP/Gdh^RNAi^* (48h, n=34, p=0.1828; 72h, n=45, p=0.9597; 96h, n=39, p<0.0001; 120h, n=55, p<0.0001), *domeMeso>GFP/Sdha^RNAi^*(48h, n=38, p=0.0374; 72h, n=49, p=0.0096; 96h, n=43, p<0.0001; 120h, n=39, p<0.0001), *domeMeso>GFP/Mdh^RNAi^* (48h, n=37, p=0.1252; 72h, n=50, p=0.0613; 96h, n=56, p<0.0001; 120h, n=41, p<0.0001). **(D)** Graph representing total proliferative capacity (mitotic index; based on total pH3^+^ nuclei count) per lymph gland, *domeMeso>GFP/+* (control, 48h, n=38; 72h, n=37, p=0.5880; 96h, n=28, p=0.8180; 120h, n=18, p=0.0773), *domeMeso>GFP/CS^RNAi^* (48h, n=28, p=0.0007; 72h, n=32, p=0.3856; 96h, n=32, p=0.1159; 120h, n=23, p>0.9999), *domeMeso>GFP/mAcon1^RNAi^* (48h, n=30, p>0.9999; 72h, n=38, p>0.9999; 96h, n=36, p>0.9999; 120h, n=31, p>0.9999), *domeMeso>GFP/α-Kdh^RNAi^* (48h, n=17, p=0.0002; 72h, n=28, p=0.6385; 96h, n=34, p>0.9999; 120h, n=19, p=0.9771), and *domeMeso>GFP/Gdh^RNAi^*(48h, n=11, p>0.9999; 72h, n=9, p>0.9999; 96h, n=18, p>0.9999; 120h, n=22, p>0.9999), *domeMeso>GFP/Sdha^RNAi^* (48h, n=26, p>0.9999; 72h, n=32, p>0.9999; 96h, n=34, p>0.9999; 120h, n=23, p>0.9999), *domeMeso>GFP/Mdh^RNAi^* (48h, n=20, p>0.9999; 72h, n=31, p=0.0477; 96h, n=27, p=0.1400; 120h, n=24, p>0.9999). **(E)** Graph representing progenitors (Dome^+^) proliferative capacity (mitotic index; based on Dome^+^ pH3^+^ nuclei count) per lymph gland, *domeMeso>GFP/+* (control, 48h, n=34; 72h, n=34, p>0.9999; 96h, n=41, p=0.7624; 120h, n=24, p=0.0730), *domeMeso>GFP/CS^RNAi^* (48h, n=25, p>0.9999; 72h, n=32, p=0.0045; 96h, n=32, p=0.4477; 120h, n=23, p>0.9999), *domeMeso>GFP/mAcon1^RNAi^* (48h, n=32, p=0.5978; 72h, n=38, p>0.9999; 96h, n=36, p>0.9999; 120h, n=31, p=0.2410), *domeMeso>GFP/α-Kdh^RNAi^* (48h, n=19, p=0.0021; 72h, n=29, p=0.2811; 96h, n=30, p=0.0353; 120h, n=19, p>0.9999), and *domeMeso>GFP/Gdh^RNAi^* (48h, n=10, p=0.1504; 72h, n=9, p=0.2641; 96h, n=18, p>0.9999; 120h, n=22, p=0.4473), *domeMeso>GFP/Sdha^RNAi^* (48h, n=33, p>0.9999; 72h, n=32, p=0.8935; 96h, n=34, p>0.9999; 120h, n=23, p=0.8963), *domeMeso>GFP/Mdh^RNAi^* (48h, n=24, p=0.3691; 72h, n=30, p=0.0062; 96h, n=27, p>0.9999; 120h, n=24, p=0.1891). **(F)** Graph representing non-progenitor (Dome^-^) proliferative capacity (mitotic index; based on Dome^-^ pH3^+^ nuclei count) per lymph gland, *domeMeso>GFP/+* (control, 48h, n=36; 72h, n=36, p<0.0001; 96h, n=32, p<0.0001; 120h, n=18, p>0.9999), *domeMeso>GFP/CS^RNAi^* (48h, n=29, p<0.0001; 72h, n=32, p>0.9999; 96h, n=32, p=0.0017; 120h, n=23, p>0.9999), *domeMeso>GFP/mAcon1^RNAi^* (48h, n=34, p=0.6504; 72h, n=38, p>0.9999; 96h, n=36, p=0.2196; 120h, n=30, p>0.9999), *domeMeso>GFP/α-Kdh^RNAi^* (48h, n=15, p>0.9999; 72h, n=32, p>0.9999; 96h, n=30, p>0.9999; 120h, n=22, p=0.8701), *domeMeso>GFP/Gdh^RNAi^* (48h, n=11, p>0.9999; 72h, n=8, p=0.9154; 96h, n=18, p=0.7298; 120h, n=22, p>0.9999), *domeMeso>GFP/Sdha^RNAi^*(48h, n=34, p=0.0582; 72h, n=32, p>0.9999; 96h, n=34, p=0.2718; 120h, n=23, p>0.9999), *domeMeso>GFP/Mdh^RNAi^* (48h, n=24, p>0.9999; 72h, n=31, p=0.1452; 96h, n=27, p=0.0216; 120h, n=24, p>0.9999). Data is presented as median plots (*p<0.05; **p<0.01; ***p<0.001; ****p<0.0001), ordinary one-way ANOVA. ‘n’=lymph gland lobes. Comparisons for significance are with control values.

**Figure 4:**
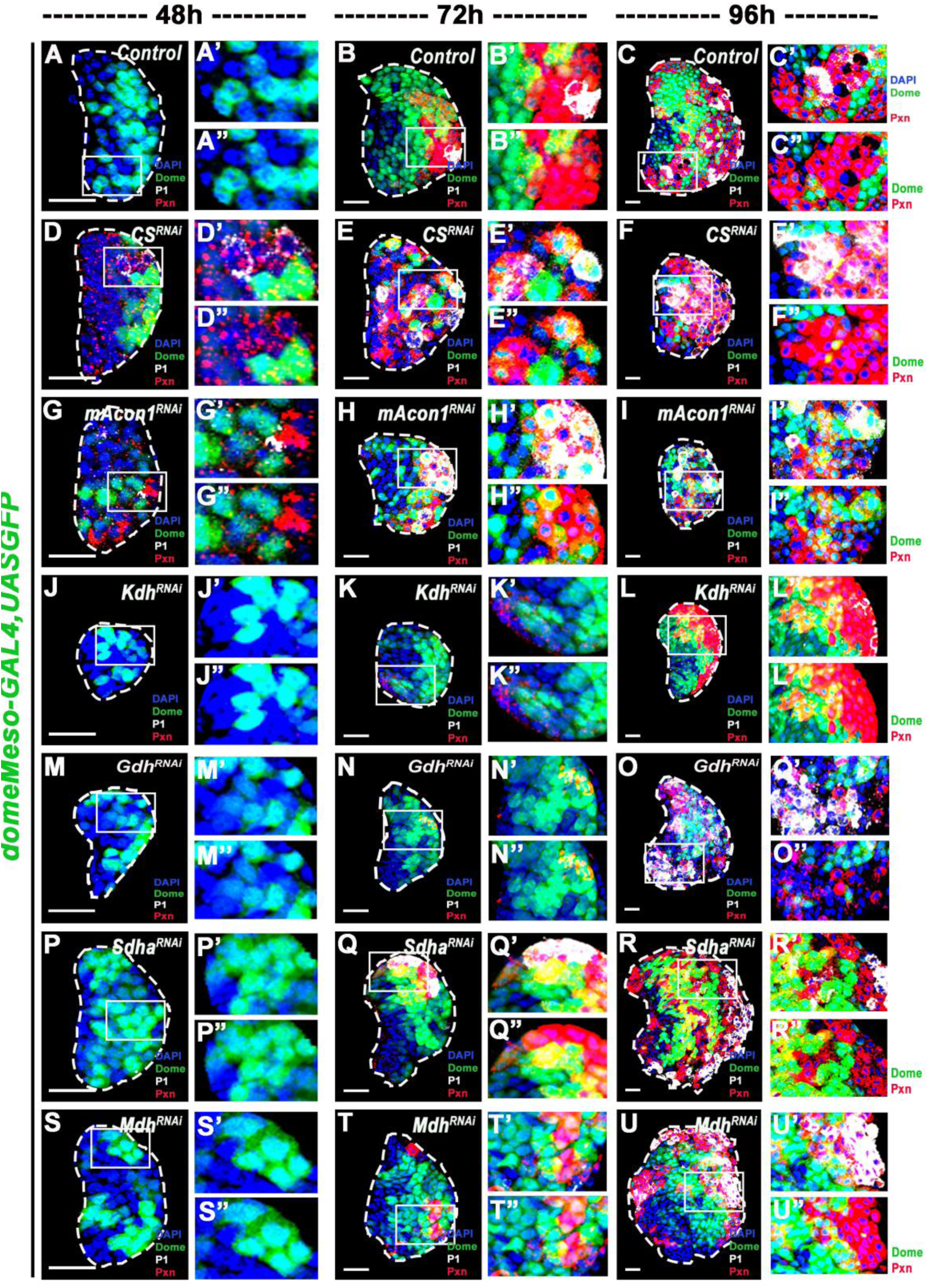
Modular control of lymph gland growth and progenitor identity by TCA enzymes. **(A-C’)** Lymph gland differentiation profile across development with differentiation marker (Pxn+, red; P1+ white) distribution in the Dome+ (green) and Dome− cells conducted temporally (48, 72 and 96 h AEL) for **(A-C”)** *domeMeso>GFP/+* (*control*) and upon loss of **(D-F”)** *domeMeso>GFP/CS^RNAi^*, **(G-I”)** *domeMeso>GFP/mAcon1^RNAi^*, **(J-L”)** *domeMeso>GFP/α-Kdh^RNAi^*, **(M-O”)***domeMeso>GFP/Gdh^RNAi^*, **(P-R”)** *domeMeso>GFP/Sdha^RNAi^*, and **(S-U”)** *domeMeso>GFP/Mdh^RNAi^*. At 48h (A-A") control lymph glands show no expression of differentiation markers (Pxn, red; P1, white), at 72h (B-B”) lymph glands show expression of differentiation marker with a few Dome+ cells expressing Pxn+ (yellow) representing the differentiating progenitor cells and Dome-cells expressing Pxn (red) and P1 (white) (see inset in B’ and B”) and (C-C”) by 96h, the lymph gland maintains a homeostatic MZ core of Dome+ progenitor population (green) with differentiating Dome-population (red and white) forming the outer CZ layer. Expression of (E-F”) *CS^RNAi^* and (G-I”) *mAcon1^RNAi^*, at (D-D” and G-G”) 48h, leads to precocious expression of differentiation markers in Dome-cells (see D’ and G’) and importantly in Dome+ cells (green cells with Pxn+ red dots in D” and G”), which (E-E” and H-H”) by 72h and (F-F” and I-I”) 96h increases progressively leading to loss of MZ progenitor pool (F and I, show reduced green population) and overall expansion of differentiated cells. Expression of (J-L”) *Kdh^RNAi^* and (M-O”) *Gdh^RNAi^*, at (J-J” and M-M”) 48h, shows no precocious expression of differentiation markers in Dome-cells or Dome+ cells, (K-K” and N-N”) by 72h, differentiation is seen comparable to controls (B-B”) but by (L-L” and O-O”) 96h differentiation is heightened showing reduction in MZ progenitor pool (L and O, show reduced green population) with increased number of differentiated cells (yellow cells in L’-L” and more white cells in O’, O”). Expression of (P-R”) *Sdh^RNAi^* and (S-U”) *Mdh^RNAi^*, at (P-P” and S-S”) 48h, shows no precocious expression of differentiation markers in Dome-cells or Dome+ cells, (Q-Q” and T-T”) by 72h, differentiation is seen comparable to controls (B-B”) but by (R-R” and U-U”) 96h differentiation is heightened showing reduction in MZ progenitor pool (R and U, show reduced green population) with increased number of differentiated cells (yellow cells in R”, U” and more white cells in R’, U”). DAPI marks DNA, Progenitors (Dome+; Green), Double positive (Dome+ Pxn+; Yellow), Differentiating population (Pxn+; Red), terminally differentiated population (P1+; white). Comparisons for significance are with control values. LG lobes are outlined with white border along with removal of other accessory tissues for clear demarcation.

Between 48-72 h after egg laying (AEL), the lymph gland doubles its cell numbers (Fig. 3A) from ∼100 cells to ∼210 cells (Fig. 3A). This increase is largely contributed by the proliferation of Dome⁺ progenitor cells, which rise from ∼60 cells at 48h to ∼150 cells at 72h (Fig. 3B). Between 72–96 h, as progenitors gradually transition to G2-phase of the cell cycle (96), the differentiating (Dome⁻) population expands in number (Fig. 3C) contributing to rapid growth of the lymph gland contributing to ∼3-fold increase in size. By 120 h, the total cell numbers in the lymph gland reaches ∼1100 cells (Fig. 3A) and the hematopoietic organ achieves its final size. At all-time points, the overall mitotic index remains stable within 3–5% as also reported in other studies (Fig. 3D, (82)), underscoring a controlled proliferation rate. This defined temporal framework enabled us to ask how the respective TCA modules aligned with the phases of blood progenitor proliferation and differentiation.

To test how knockdown of *CS* or *mAcon1* affected LG growth and progenitor proliferation we conducted temporal analysis of growth (Fig. 3) and proliferative index (Fig. S6) from 48–120 h AEL. Between, 48–72 h, in knockdown of *CS* or *mAcon1* condition, total LG cell numbers remained comparable to controls (Fig. 3A), but by 72h, growth sharply declined (Fig. 3A) as evident by comparatively fewer total cell numbers seen in these *RNAi* condition. Interestingly, when assessed for cell numbers across different cellular pools, at early 48-72h stages, Dome⁺ progenitors were significantly reduced (Fig. 3B), but the Dome⁻ cells showed a compensatory increase (Fig. 3C). This increase masked the early growth defect. However, the compensation collapsed after this and by 96 h, Dome^-^ cells could no longer demonstrate a rise in their numbers (Fig. 3C) leading to smaller LGs (Fig. 3A). Quantifications of mitotic indices as undertaken by counting the number of phosphorylated-histone H3 positive nuclei (pH3^+^. (82), Fig. S6), it revealed that while overall proliferative index was unchanged (Fig. 3D, & S6A-I’), compared to Dome^+^ cells (Fig, 3E), the Dome⁻ cells exhibited a transient spike in division at 48h (Fig. 3F), but later was normalized back to control levels (Fig. 3D and F). Compared to lymph glands from controls (Fig. S6A-A’), the increase in pH3^+^ nuclei were dramatic in *CS^RNAi^* condition (Fig. S6D, D’), where a significant spike was seen, while in *mAcon1^RNAi^* condition, the increased trend in mitotic competency was although evident in majority of lymph glands (Fig. S6G & G’), but it failed to achieve statistically significance (Fig. 3F).

Next, we analyzed *α-Kdh/Gdh* axis from 48–120 h AEL. Expressing *α-Kdh^RNAi^*or *Gdh^RNAi^*under *domeMESO-GAL4* revealed entirely different growth dynamics. Across all stages, LGs in these loss of function conditions were significantly smaller and contained fewer total cells than controls (Fig. 3A, Fig. S6J-O’), with a disproportionate reduction in Dome⁺ progenitor cell numbers (Fig. 3B). Especially at 48h, when in control LGs the progenitor numbers are ∼60 (Fig. S6A), this was limited to only ∼20 cells in *α-Kdh^RNAi^* (Fig. S6J) and *Gdh^RNAi^* (Fig. S6M) condition. The number of Dome⁻ cells however remained unaffected (FIG. 3C, Fig, S6J and M). Importantly, pH3 expression analysis revealed a significant reduction in mitotic index at 48h in *α-Kdh^RNAi^* condition, arising from reduced proliferative capacity of Dome^+^ cells (FIG 3D, E, S6J and M). Dome⁻ cells showed no change in mitotic competency (FIG 3F, Fig. S6J and M) and remained comparable to control. These results indicated that early reduction in Dome⁺ cell number precedes any depletion of Dome⁻ cells, demonstrating defective progenitor proliferation. This limits overall tissue expansion from early stages itself, indicating that the *α-Kdh/Gdh* module in progenitor cells was necessary to autonomously support their proliferation. Thus, the *α-Kdh/Gdh* axis defines a growth dedicated metabolic node, required to sustain the early proliferative burst in the progenitor cells that establishes their quorum. These phenotypes were distinct from *CS/mAcon1* activity in progenitors. While the *CS/mAcon1* activity autonomously regulated progenitor identity (Fig. 3B), the axis also controlled non-autonomously the early proliferation of neighboring Dome^-^ cells (Fig. 3F). These Dome^-^ cells are presumably the core progenitors described previously as pre-prohemocytes residing adjacent to the dorsal vessel (51) which gradually initiate expression of Dome-Gal4 marker and subsequently mature into prohemocytes.

Our findings reveal that the *CS/mAcon1* module serves dual functions: autonomously preserving progenitor/prohemocytes cell number and non-autonomously restraining the expansion of the Dome^-^ core progenitor population. Thereby ensuring a balanced growth between distinct blood-progenitor populations, necessary for proper lymph gland growth.

To further examine the impact of *CS*/*mAcon1 and α-Kdh/Gdh* loss on progenitor population, we stained early LGs (48 and 72 h AEL) for differentiation markers Pxn and P1. In controls, neither marker was detectable at 48 h (Fig. 4A-A’’), but both were detected at 72 h (Fig. 4B-B”) and were restricted to the developing CZ (Fig. 4B). From 72h onwards the Pxn^+^ cells and P1^+^ cells expanded (Fig.4B-C”) and reached almost 40% of the total LG area by 96h (Fig. S7A & B). The distinct spatial restriction of these differentiating markers to the outer layer indicated a rigorous control on progenitor homeostasis alongside initiation of the differentiation program and formation of the CZ.

In *CS^RNAi^* (Fig. 4D-F”) and *mAcon1^RNAi^*(Fig. 4G-I”), LGs displayed precocious expression of Pxn as early as 48 h on Dome^+^ cells, with subsets of Dome^-^ cells also initiating the expression of Pxn and also P1 markers (Fig. 4D-D”, G-G”). This indicated premature entry of LG blood progenitor cells into differentiation and also attainment of terminal differentiation. By 72 h, Dome⁺ progenitor numbers declined sharply and there was a significant rise in differentiating cells expressing Pxn and P1 markers (Fig. 4E-E”, H-H”, Fig. S7A, B), compared to control (Fig. 4B-B”). This pattern of extensive differentiation persisted at 96h both in *CS* and m*Acon1* loss of function conditions (Fig. 4F-F”, I-I”, Fig. S7A, B). This pattern revealed an early loss of progenitor identity with the onset of differentiation trajectory. The data showed that Dome^+^ cells lacking *CS* and *mAcon1*, prematurely exited the progenitor state and adopted Pxn⁺ identity and subsequently become P1⁺ prematurely. Moreover, the core-progenitor state (Dome^-^ cells) was also impaired as they too embarked on a differentiation trajectory. These data are consistent with impaired maintenance of progenitor identity and we speculate the rise in proliferation at 48h seen in the Dome^-^ pre-prohemocytes/core progenitor compartment (Fig. S6D & G) as the underlying cause for their precocious entry into the differentiation program. Thereby, leading us to propose a model where *CS*/*mAcon1* loss in the progenitors disrupts early progenitor-to-differentiation cell cycle checkpoint and causes premature exhaustion of progenitor cells leading to LG growth defect.

Conversely, progenitor specific loss of *α-Kdh* and *Gdh,* even though led to failure of progenitor cells to proliferate early, these cells retained Dome-GFP progenitor marker expression and remained devoid of differentiation markers at 48 h (Fig. 4J-J” & M-M”) indicating an arrest in their development while retaining undifferentiated states and progenitor identity. At later stages (72–96 h), *α-Kdh^RNAi^* and *Gdh^RNAi^* expressing LGs showed increase in P1 expression (Fig. 4K-L” and N-O”, Fig. S7A, B), suggesting that there was no change in temporal protocol for initiation of differentiation. However later in progenitor development, the loss of these enzymes led to rapid rise in differentiation, indicating loss of progenitor homeostasis.

When we assessed for progenitor homeostasis in *Sdh^RNAi^* and *Mdh^RNAi^* expressing LGs (Fig. 3A-F, Fig. S6Q-V’ and Fig. 4P-U”, Fig. S7A, B). We observed no change in overall size as total cell numbers remained unaffected (Fig. 3A-C, Fig. S6Q-V’). Also, across populations, from 48-72h, Dome^+^ and Dome^-^ cells demonstrated profiles comparable to control (Fig. S6Q-R’, and S6T-U’, compared to control in S6A-B’), but later at 96-120 h Dome^+^ cells were significantly reduced (Fig. 3B) but a corresponding rise in Dome^-^ CZ population was evident (Fig. 3C). Temporal assessment of differentiation marks in these LGs displayed no change in differentiation profile until 72 h (Fig. 4P-Q” & S-T”), but at 96h, the loss of these enzymatic steps exhibited an increase in differentiation markers (Fig. 4R-R”, & U-U”) which gets exaggerated with time and heightened differentiated is seen at later stages of development (Fig. 1M). Thus, loss of *Sdh* and *Mdh* although did not disrupt the normal temporal onset of maturation, their loss impaired later stages of blood progenitor homeostasis and led to expansion in the expression of Pxn in Dome^+^ progenitor cells. Thus, leading to dysregulation in balance between progenitor homeostasis and differentiation. This finding explains the reduction in Dome^+^ cell counts detected at 96h, as described earlier (Fig. 3B).

Together, these results delineate three distinct metabolic logics governing progenitor development. The *α-Kdh/Gdh* axis, that acts autonomously within progenitors to sustain their proliferation all throughout development. The *CS/mAcon1* axis contrarily preserves progenitor identity and prevents their premature differentiation early in LG development. Additionally, it non-autonomously restrains the proliferation of core-progenitor cells thereby enabling a framework that restricts their precocious differentiation. Together, the dual TCA enzymatic node, establishes a competent pool of long-term proliferative progenitor cells that contribute to overall LG dynamics (Fig. S7C). Later, as development progresses, and differentiation is initiated, and when progenitor cells enter homeostasis and maintain MZ quorum, the modular enzymatic activities of the TCA enzymes gradually transitions into a unified, continuous TCA loop through the engagement of *Idh*, at later 96h onwards. This circular mode of TCA is driven by *Pdh* alluding to an enhanced oxidative state, that supports progenitor homeostasis (Fig. S7C). Altogether, the modular to circular TCA transition supports proportional progenitor proliferation, growth and homeostasis.

### Citrate supplementation rescues premature differentiation and reveals a metabolic transition from modular to complete TCA cycling

Next, we wanted to confirm the functional entity of the *CS/mAcon1* module in maintaining progenitor state. To do this, we performed dietary rescue assays using citrate and isocitrate the direct metabolic end products of *CS* and m*Acon1*, respectively (Fig. 5A). Larvae expressing *CS^RNAi^* were fed either 1% citrate or isocitrate and lymph gland development was assessed (Fig. S8A). Continuous feeding throughout larval life further reduced LG size (Fig. S8D compared to C and B), suggesting possible metabolic stress due to systemic metabolite overload. Therefore, we employed a stage-specific feeding regimen of supplementing larvae from first instar until 60 h AEL, the developmental window when *CS^RNAi^* phenotypes first appear and subsequently returning them to normal diet (Fig. S8A). This early and transient supplementation was remarkably effective: citrate feeding significantly rescued the size and differentiation defects seen in *CS^RNAi^* LGs (Fig. S8E). When temporally analysed with this feeding regime (Fig. 5A and B), at 48 h, compared to control (Fig. 5C), the reduction seen in Dome⁺ progenitor numbers in *CS^RNAi^* condition (Fig. 5D) was restored on citrate diet (Fig. 5E and O). Differentiation markers (Pxn and P1) also normalized (Fig. 5D-E’ and O), and the gland growth matched control size (Fig. 5C-E’ and P). Notably, this rescue persisted after withdrawal of citrate, at 72h (Fig 5G-I’ and Q, R) and later at 120h (Fig. 5K-M’, S, and T) as well. Importantly, larvae exposed to citrate later, post 60 h of development, these LGs showed no rescue rather elevated differentiation (Fig. S8F), identifying a critical early window (< 60 h AEL) when citrate availability was necessary and sufficient to maintain progenitor identity and establish their long-term proliferative potential. These results demonstrated that once progenitor balance was re-established, it remained stable, most likely via epigenetic modes of regulation. We also tested early 1%CF supplementation on *mAcon1^RNAi^* expressing condition and failed to see any rescue of either progenitor homeostasis or LG growth (Fig S8B, G & H). This highlighted the requirement of *mAcon1* function downstream of *CS*, and also alluded to specificity of citrate rescue, and that it did not non-specifically rescue any differentiation phenotype.

**Figure 5:**
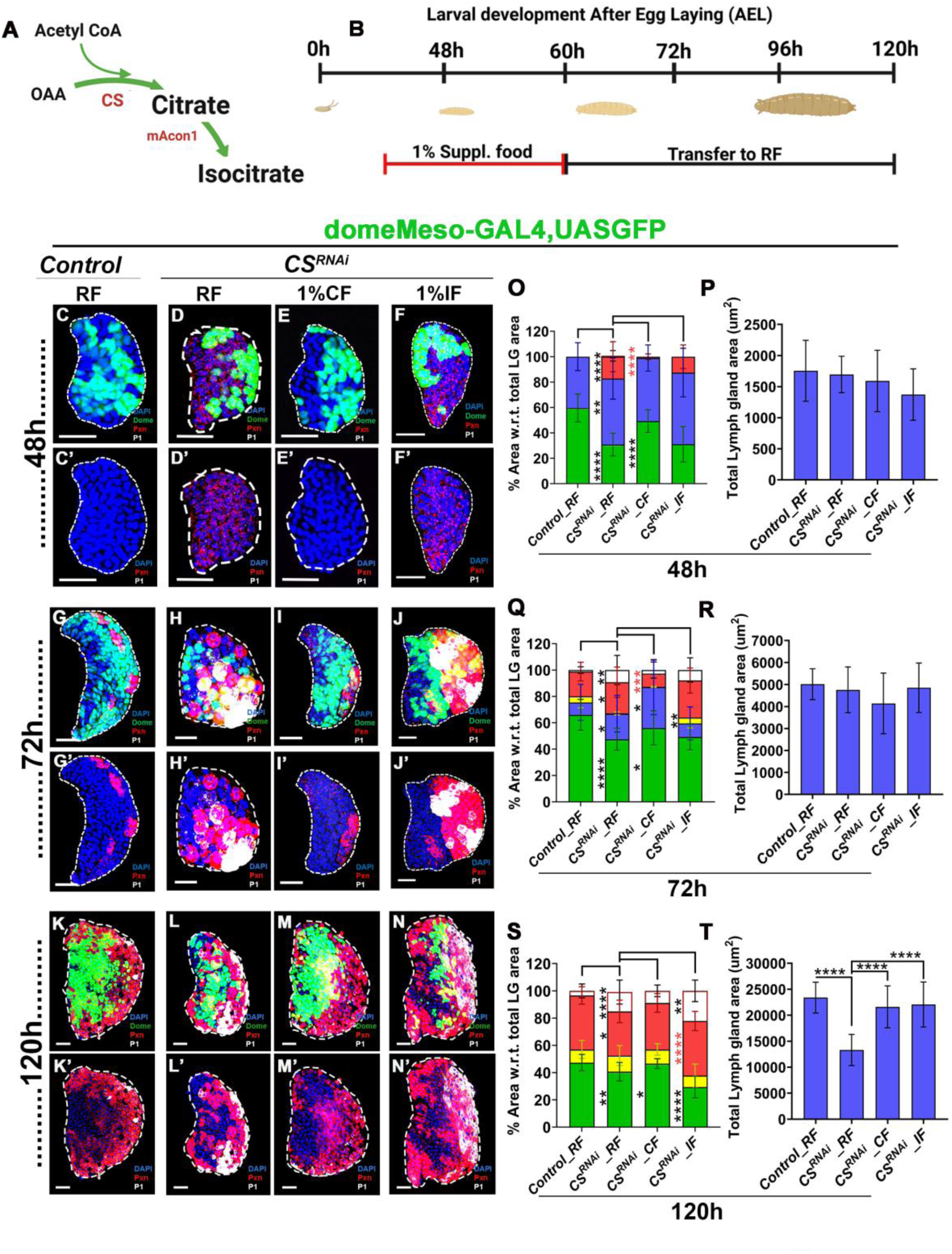
Citrate is important for early progenitor identity and development of lymph glands. **(A)** Schematic representing the the enzymatic step catalyzed by citrate synthase (CS) enzyme leading to the formation of citrate from Oxalo-Acetic Acid (OAA) and acetyl CoA and its conversion to Isocitrate by aconitase (mAcon1) enzyme. **(B)** Developmental timeline of dietary rescue experiments of *domeMeso>GFP CS^RNAi^*, where the red line represents the time frame when first instar larvae (24h AEL) of *domeMeso>GFP/CS^RNAi^*were exposed to food supplemented with 1% citrate food (CF) or 1% isocitrate food (IF), and grown (60hAEL), after which they were transferred to regular food (RF, black line). **(C-F’)** Representative lymph glands images at 48h of **(C-C’)** *Control* (domeMeso-Gal4,UAS-GFP/+) LGs on regular food (RF) with progenitors (Dome+; green) and non-progenitors (Dome⁻; blue) areas, with no detectable Pxn staining, **(D-D’)** *CS^RNAi^ (domeMeso-Gal4,UAS-GFP;UAS-CS^RNAi^*), on RF at 48h showing pronounced differentiation, where progenitor reduced, non-progenitors (Dome⁻; blue) and differentiating (Pxn+; red) cell area, are subsequently increase in comparison to **(C-C’)** *control* **(E-E’)** *CS^RNAi^ (domeMeso-Gal4,UAS-GFP;UAS-CS^RNAi^*), on 1% CF at 48 h, showing rescue of differentiation phenotype with progenitors (Dome^+^; green) area increased and non-progenitors (Dome⁻; blue) with negligible Pxn+ cells and **(F-F’)** *CS^RNAi^ (domeMeso-Gal4,UAS-GFP;UAS-CS^RNAi^*), on 1% IF showing no recovery of differentiation phenotype, where progenitor remain reduced and non-progenitors (Dome⁻; blue) continued to express differentiating markers (Pxn+; red), in comparison to (D-D’) *CS^RNAi^*on RF. **(G-J’)** Representative lymph gland Images at 72h from **(G-G’)** control *(domeMeso-Gal4,UAS-GFP/+*), on RF, **(H-H’)** *CS^RNAi^ (domeMeso-Gal4,UAS-GFP;UAS-CS^RNAi^*), on RF at 72h showed pronounced differentiation, where progenitor were reduced, which is complemented by increase in non-progenitors (Dome⁻; blue) and differentiating (Pxn+; red) cell area, was along with a decent increase in terminally differentiation (P1+; white), in comparison to **(G-G’)** control on RF. **(I-I’)** *CS^RNAi^ (domeMeso-Gal4,UAS-GFP;UAS-CS^RNAi^*), on 1% CF with dramatic recovery of differentiation phenotype even when moved to RF, with progenitors (Dome+; green) as well as non-progenitors (Dome⁻; blue) area increased in lieu of differentiating (Pxn^+^; red) cell area, was along with a notable reduction in terminally differentiated (P1^+^; white) and **(J-J’)** *CS^RNAi^ (domeMeso-Gal4,UAS-GFP;UAS-CS^RNAi^*), on 1% IF, excessive differentiation did not rescue, in comparison to (G-G’) *CS^RNAi^* on RF. **(K-N’)** Representative LG images at 120h in development, from **(K-K’)** control *(domeMeso-Gal4,UAS-GFP/+*), on RF showing progenitors (Dome^+^; green), vs Pxn^+^ (red) area which can further classified into Dome^+^Pxn^+^ (yellow), and P1^+^ (white), **(L-L’)** *CS^RNAi^ (domeMeso-Gal4,UAS-GFP;UAS-CS^RNAi^*), on RF showed enhanced differentiation with progenitors (Dome+; green) reduction due to expansion of Pxn^+^ (red) area of which there is Dome^+^Pxn^+^ (yellow), and P1^+^ (white); with also a LG growth defect, (compare with **(K-K’)** *control*), **(M-M’)** *CS^RNAi^ (domeMeso-Gal4,UAS-GFP;UAS-CS^RNAi^*), on 1% CF, restored the progenitors to differentiating population homeostasis ratio and no LG size defect is detected and **(N-N’)** *CS^RNAi^ (domeMeso-Gal4,UAS-GFP;UAS-CS^RNAi^*), on 1% IF, differentiation homeostasis could not be restored as Dome^+^ progenitors are further reduced, with a concomitant increase of Pxn^+^ (red) area, Dome^+^Pxn^+^ (yellow), and P1^+^ (white); but LG growth is restored in comparison to small LGs seen In **(L-L’)** *CS^RNAi^* on RF at 120h. **(O)** Graphs showing quantification of differentiation profile based on relative % area with respect to total lymph gland (LG) area at 48h, *domeMeso>GFP/+* (*control*, 48h, RF, n=36), *domeMeso>GFP/CS^RNAi^* (48h, RF, n=31, green, p<0.0001; blue, p=0.0019; red, p<0.0001), *domeMeso>GFP/CS^RNAi^* (48h, 1%CF, n=30, green, p<0.0001; blue, p=0.7756; red, p<0.0001), *domeMeso>GFP/CS^RNAi^* (48h, 1%IF, n=11, green, p=0.9958; blue, p=0.6082; red, p=0.2246), **(P)** graph representing the quantification of total LG area, *domeMeso>GFP/+* (control, 48h, RF, n=26), *domeMeso>GFP/CS^RNAi^* (48h, RF, n=29, p=0.9263), *domeMeso>GFP/CS^RNAi^* (48h, 1%CF, n=25, p=0.5503), *domeMeso>GFP/CS^RNAi^* (48h, 1%IF, n=11, p=0.0497), **(Q)** graph showing quantification of differentiation profile based on relative % area with respect to total lymph gland (LG) area at 72h, *domeMeso>GFP/+* (control, 72h, RF, n=26), *domeMeso>GFP/CS^RNAi^*(72h, RF, n=26, green, p<0.0001; blue, p=0.0472; yellow, p=0.0338; red, p=0.0454; white, p=0.0016), *domeMeso>GFP/CS^RNAi^* (72h, 1%CF, n=11, green, p=0.0384; blue, p=0.0269; yellow, p=0.9947; red, p=0.0002; white, p=0.1270), *domeMeso>GFP/CS^RNAi^* (72h, 1%IF, n=13, green, p=0.8237; blue, p=0.2911; yellow, p=0.0080; red, p=0.1877; white, p=0.9053), **(R)** graph representing the quantification of total LG area, *domeMeso>GFP/+* (control, 72h, RF, n=22), *domeMeso>GFP/CS^RNAi^* (72h, RF, n=19, p=0.7741), *domeMeso>GFP/CS^RNAi^* (72h, 1%CF, n=10, p=0.3107), *domeMeso>GFP/CS^RNAi^* (72h, 1%IF, n=11, p=0.9683), **(S)** graph showing quantification of differentiation profile based on relative % area with respect to total lymph gland (LG) area at 120h, *domeMeso>GFP/+* (control, 120h, RF, n=17), *domeMeso>GFP/CS^RNAi^* (120h, RF, n=33, green, p=0.0039; yellow, p=0.6186; red, p=0.0312; white, p<0.0001), *domeMeso>GFP/CS^RNAi^* (120, 1%CF, n=10, green, p=0.0357; yellow, p=0.8070; red, p=0.1384; white, p=0.1307), *domeMeso>GFP/CS^RNAi^* (120h, 1%CF, n=20, green, p<0.0001; yellow, p=0.1934; red, p<0.0001; white, p=0.0018), **(T)** Graphical represention of quantification of total LG areas, in *domeMeso>GFP/+* (control, 120h, RF, n=25), *domeMeso>GFP/CS^RNAi^*(120h, RF, n=33, p<0.0001), *domeMeso>GFP/CS^RNAi^* (120h, 1%CF, n=19, p<0.0001), *domeMeso>GFP/CS^RNAi^* (120h, 1%IF, n=18, p<0.0001). Data is presented as median lots (*p<0.05; **p<0.01; ***p<0.001; ****p<0.0001), ordinary one-way ANOVA. Scale bar: 20µm. ‘n’=lymph gland lobes. DAPI marks DNA, Progenitors (Dome+; Green), non-progenitor (blue) Double positive (Dome^+^ Pxn^+^; Yellow), Differentiating population (Pxn^+^; Red), terminally differentiated population (P1^+^; white). Comparisons for significance are with control values for RF while for supplement food (rescues) it was done with *CS^RNAi^* on RF. LG lobes are outlined with white border along with removal of other accessory tissues for clear demarcation.

**Figure 6:**
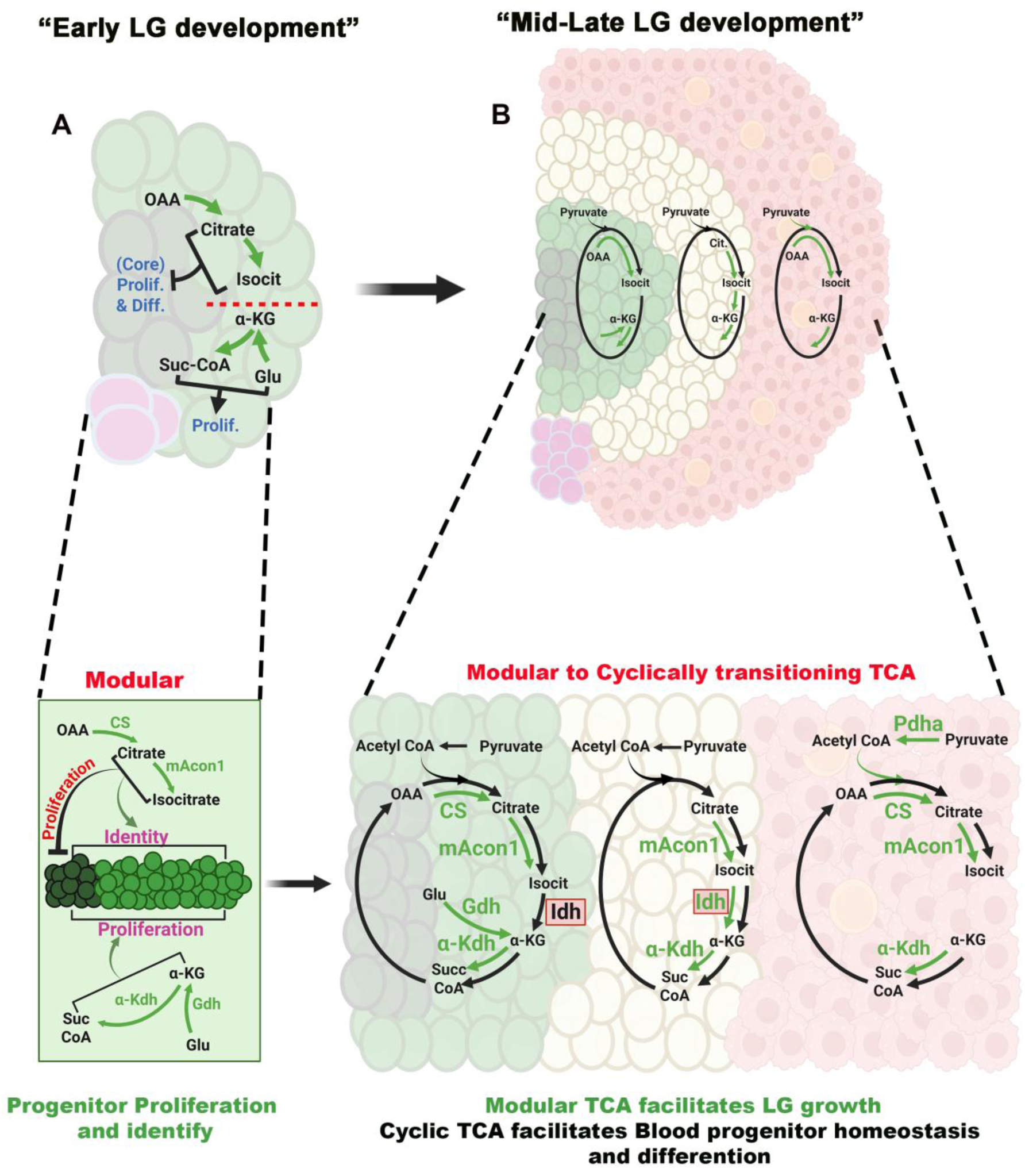
Metabolic reorganization from modular to cyclic TCA configuration during lymph gland development (A) Early stage: (Modular configuration), At early stage in lymph gland (LG) development, progenitors exhibit a modular metabolic organization of TCA cycle. Key metabolites (OAA, citrate, isocitrate, α-ketoglutarate, glutamate, and succinyl-CoA) are locally produced and utilized without completing the full oxidative TCA cycle. Citrate and isocitrate keeps a check on the proliferative capacity of the non-progenitor population (blue). Distinct enzymatic modules *citrate synthase (CS), Aconitase (mAcon1), α-Ketoglutarate dehydrogenase (α-Kdh)*, and *Glutamate dehydrogenase (Gdh)* operates in semi-independent manner to sustain progenitor identity and proliferative capacity. **(B) Mid-Late stage:** (cyclic configuration), as development progress, the LG transition towards cyclic TCA organization. The induction of Pyruvate dehydrogenase (*Pdh*) and Isocitrate dehydrogenase (*Idh*) activities drives the conversion of pyruvate to acetyl CoA and enhance citrate to α-KG flux, establishing a fully functional oxidative cycle. This metabolic reorganization integrates metabolic modules across spatial zones; progenitors (green), intermediary progenitors (yellow), and differentiating hemocytes (red) to coordinate homeostatic differentiation with zonal proliferation. Magnified views of the three metabolic zones are shown. **Left:** Progenitor-enriched zone retains the modular TCA configuration characterized by high *CS*, *mAcon1*, *Gdh*, and *α-Kdh* expression, enabling α-KG-centered metabolism essential for progenitor proliferation and identity. **Middle:** Intermediate zone shows a transition state, where increasing pyruvate entry through *Pdh* expands metabolic flux. *Idh* becomes active, supporting redox balance and enabling metabolic flexibility during differentiation initiation. **Right:** Differentiated cell zone displays a more complete oxidative cycle, with active *Pdh* and full progression of TCA intermediates, supporting the energetic and biosynthetic demands of mature blood cells. Overall this model proposes that the developing LG transition from an early modular to a late cyclic metabolic architecture. Such a shift enables a progressive coupling between anabolic progenitor programs and catabolic differentiation processes, ensuring metabolic integration and tissue level homeostasis during LG maturation.

Isocitrate feeding of *CS^RNAi^* condition on the other hand, restored the phenotypes partially. While, progenitor imbalance and premature differentiation were still retained at 48h (Fig. 5F-F’ & O) and continued to be seen later at 72h (Fig 5J, J’ & Q) and 96h, (Fig 5N, N’ & S), LG growth restoration was seen at 120h (Fig. 5T). No change in LG size was however detected, at early 48h (Fig. 5P) or 72h (Fig. 5R), indicating that the rescues were not a consequence of any early increase in lymph gland size. Rather, early isocitrate feeding was able to resolve the late lymph gland proliferative defect in *CS^RNAi^* condition (Fig. 3A and C).

Overall, the supplementation data support complete sufficiency of citrate in moderating early progenitor development, both their identity and building proliferative capacity, however isocitrate only restored their proliferative capacity, without correcting their identity. While this may imply differential non-overlapping modes of action of the two metabolites, we speculate that the intracellular thresholds of isocitrate needed to revert identity were not met in our dietary rescues. Unfortunately, when supplemented with higher doses of isocitrate, it led to developmental delays. However, based on our genetic data that demonstrate a requirement for both CS and mAcon1 activity, we conclude that citrate and isocitrate production by the *CS/mAcon1* module sustains progenitor identity and prevents their precocious differentiation. The node also supports the proliferative competency of the blood progenitor cells in the longer term as well.

Importantly we also conducted similar rescues with succinate food (68, 69, 97), given it is a downstream metabolic outcome of α-KG metabolism and asked if its availability would restore the growth defect seen in *α-Kdh* mutants (Fig. S8B, I & J). Interestingly, we did not observe any rescue indicating that the proliferation defect was not a consequence of unavailability of succinate (Fig. S8I-L). The data strengthen the unique early requirement supported by *α-Kdh* in succinyl-CoA generation as opposed to driving succinate oxidation via the TCA. Lack of any LG size defect in *Sdh* and *Mdh* knockdowns, further strengthened this notion (Fig 3A).

### *Idh*, a metabolic buffer to sustain progenitor homeostasis by completing the TCA cycle

As demonstrated by our results, summarised in Sup FIG 7C, beyond the stage of a modular TCA configuration, to sustain long-term progenitor homeostasis, the cycle transitions into a complete loop. The involvement of *Idh*, positioned between the *CS/mAcon1* and *α-Kdh* modules, emerges as a pivotal step linking these two metabolic nodes, and transitions the TCA from modular to cyclical in governing progenitor homeostasis. Moreover, the involvement of Pdh at the later time point driving pyruvate oxidation demonstrates a rise in TCA flux and indicative of a distinct metabolic framework for progenitor maintenance. In this case, the specific, loss of *Idh* in Dome⁺ progenitor cells leading to excessive differentiation could arise as a consequence of isocitrate accumulation. With respect to this notion, we therefore asked if raising TCA metabolites like citrate or isocitrate perturbed progenitor homeostasis.

We undertook the same feeding strategy as done previously, but here control larvae were fed with citrate or isocitrate early (<60h) (Fig. S9A-A’) as the means to raise the levels of these metabolites beyond the baseline. As we raised citrate/isocitrate availability in the system early (<60h), we analysed lymph glands temporally at 48h, 72h and 120h AEL for any changes. We observed that citrate feeding early at 48h mildly reduced progenitor cell population (Fig. S9C, C’ & K), but surprisingly, in the case of isocitrate feeding, progenitor reduction with a stark increase in differentiation marker Pxn was evident (Fig. S9D-D’ & K). At 72h, while citrate feeding was comparable to control (Fig. S9 F, F’& M), on isocitrate diet the progenitor (green) proportion continued to show reduction with more dramatic increase in non-progenitor population (blue) (Fig. S9G, G’ & M, in comparison to control Fig. S9E & E’). Importantly, at this stage, isocitrate feeding also showed a strong LG growth retardation, whereas no significant change was observed on citrate diet at 72h (Fig. S9N). Later, at 120h, even when the animals were no longer raised on the supplemented diet, progenitor homeostasis emerged compromised now in both citrate (Fig. S9I, I’ & O) and isocitrate (Fig. S9J, J’ & O) condition. A reduction in progenitor proportion with subsequent increment in differentiating (Pxn+) population was evident (Fig. S9H & H’) and isocitrate condition continued to be growth retarded but no changes on growth was however detected on citrate diet (Fig. S9P). These results indicated deleterious effects of excessive isocitrate and citrate on blood progenitor state and proliferation. The observations alluded us to growth limiting and pro-differentiation properties of these metabolites, when presented in excess. Leading us to propose a model where, maintaining homeostatic levels of TCA metabolites in development was also necessary for proper blood progenitor development.

## Discussion

Hematopoietic systems across species must coordinate stem cell identity, proliferative expansion, lineage commitment, and tissue scaling with developmental and systemic cues. Our findings reveal that the *Drosophila* lymph gland achieves this coordination through a two phase TCA metabolic architecture. During early larval stages, when the lymph gland expands predominantly through stem like progenitors, the TCA operates in a modular, low-oxidative configuration. This state stabilizes progenitor identity and restrains premature differentiation. As larvae enter mid second instar, these stem like progenitors generate intermediate, lineage-primed progenitors, reminiscent of transitional multipotent progenitors (MPPs) in vertebrate hematopoiesis. These IZ cells proliferate and differentiate while maintaining the progenitor pool. It is during this maturation period that the lymph gland undergoes a major metabolic transition: the previously independent *CS/mAcon1* and *Kdh/Gdh* nodes become connected by *Pdh* and *Idh*-dependent oxidative flux, forming a fully cyclic TCA network across the MZ and IZ. This integrated TCA mode supports both differentiation and progenitor homeostasis. The developmental transitions seen in the lymph gland underscore a broader principle: TCA topology itself acts as a regulatory layer that times hematopoietic progression. Early hematopoiesis requires metabolic strategies that stabilize identity and support controlled expansion with minimal oxidative burden, whereas later stages engage oxidative TCA flux to fuel differentiation, biosynthesis, and tissue maturation. The lymph gland therefore provides a developmental blueprint for how metabolic architecture synchronizes stemness, lineage priming, and differentiation with organismal physiology.

These findings parallel key features of mammalian hematopoietic stem cell (HSC) regulation. The early modular state is analogous to quiescent or early-activated HSCs, which also rely on node-specific anaplerosis and minimized oxidative load to preserve stemness (98–101). The later phases, mirror the HSC-to-MPP transition where oxidative metabolism becomes increasingly important for lineage output while maintaining regulatory control over progenitor equilibrium (99, 102–104).

A central feature of the early modular TCA architecture is the no or low *Pdh* activity in progenitor cells but elevated *CS* activity leading to citrate/isocitrate production and establishment of progenitor identity. Early citrate supplementation mediated long lasting recovery of *CS*-deficient progenitor differentiation and growth phenotypes indicates a narrow developmental window during which citrate availability “sets” progenitor identity in a durable manner. These rescues by brief citrate exposure suggest the establishment of an epigenetic memory by *CS/mAcon1* node. In this context, early progenitor cells maintain a state of low *Pdh* activity but elevated *CS/mAcon1* activity. This state will rapidly channel available acetyl-CoA and oxaloacetate into citrate. Thus, this *CS*-driven node may be a way to partition acetyl-CoA toward mitochondrial use, limiting its export for cytosolic/nuclear acetyl-CoA production and availability for histone acetylation, whereby progenitors can avoid hyper-acetylation driven chromatin opening (21–23, 105, 106) and preserve a stem-like transcriptional landscape. Moreover, cytosolic citrate can inhibit PFK (107) which can further restrain glycolysis mediated acetyl-CoA generation. Together, these mechanisms would create a metabolically enforced identity state, where early progenitor cells rather than being acetyl-CoA rich, they actively buffer and compartmentalize acetyl-CoA flux, ensuring a metabolically constrained state that stabilizes progenitor identity.

In parallel, α-KG produced by the *Gdh/Kdh* node serves as the main driver of early progenitor proliferation. Our data clearly separate α-KG function from its downstream metabolite succinate. α-KG supports proliferation through anaplerosis, nitrogen handling, mTORC1 signaling, and as a cofactor for α-KG–dependent histone demethylases mechanisms central to proliferative competence and identity regulation in vertebrate HSCs (1, 108–113). Importantly, what the system requires downstream of α-KG is its regulated oxidation to succinyl-CoA, not succinate. Succinyl-CoA lies at a metabolic–epigenetic intersection: it donates succinyl groups for lysine succinylation, modulating chromatin and mitochondrial enzyme function; it fuels heme biosynthesis essential for proliferative competence; and it contributes to anaplerotic buffering that stabilizes TCA continuity during rapid growth. In contrast, succinate neither supports these functions nor substitutes for α-KG, instead, it competitively inhibits α-KG dependent dioxygenases and can antagonize chromatin programs required for progenitor expansion.

The transition to oxidative TCA operation is accompanied by increased *Pdh* activity, and *Idh* engagement. The emergence of physiological ROS in the MZ (42), that is mitochondria derived are in alignment with our findings. The data also parallel the appearance of ROS as a differentiation-sensitizing cue in mammalian HSCs and MPPs (114–117). Balanced ROS is essential to sensitize progenitor cells to differentiation cues, however its excess drives premature differentiation, whereas insufficient oxidative throughput destabilizes progenitor maturation potential. Mitochondrial Ca²⁺, which regulates *Pdh*, *Idh,* and *Kdh* function, likely tunes this oxidative window, and is consistent with the fact that progenitor cells of the MZ maintain elevated cytosolic Ca²⁺ that is necessary to maintain their identity and also proliferation (118). Interestingly, our recent findings show systemically derived GABA, whose metabolism in the progenitor cells acting through the GABA shunt restrains *Pdh* activity and buffer ROS. Together with our current data, the findings, echo the role of systemic inputs and extrinsic metabolites also niche-derived factors in fine tuning blood-progenitor TCA activity.

Interestingly, within this spatially patterned metabolic landscape, the intermediate zone (IZ) progenitor cells comparable to transitional progenitors in vertebrate bone marrow(46, 70, 81, 119–121), they emerge as a metabolic relay compartment. Perturbations of TCA nodes in the IZ affect growth and differentiation output, positioning this population as the rate-limiting integrator of metabolic, epigenetic, and lineage signals. IZ cells perhaps integrate metabolite fluxes from the MZ and CZ and engage full TCA cycling to restore metabolic balance when fluxes or ROS levels deviate from optimal ranges (Fig. S10). The loss of specific TCA nodes, may disrupt this integration leading to dysfunctional hematopoiesis. This architecture of transitional cells as integrator of metabolic fluxes, provides a centralized mechanism for maintaining proportional lineage output and progenitor homeostasis.

While modular TCA architectures have been described in bacteria, plants, tumors, and immune cells, the lymph gland model represents a developmentally programmed form of TCA branching. In tumors or macrophages, truncated cycles arise from adaptive stress responses (12, 31, 122), whereas in the lymph gland, modularity is embedded in the developmental program. The switch from modular to complete cycling occurs predictably with developmental time and is spatially encoded across hematopoietic zones. This makes the lymph gland the first example of a temporally orchestrated metabolic architecture, where the pathway trajectory itself acts as a morphogenetic regulator. We speculate this modality may give *Drosophila* hematopoietic system the means to be sensitive to extrinsic modulations. The development of the blood progenitor cells is deeply reliant on systemic nutrition and PSC-derived niche cues (47, 70, 79, 79, 123–129). The metabolic checkpoints and framework uncovered here are likely to operate downstream of insulin/TOR or GABA metabolic signaling (68, 70, 97), thereby aligning hematopoietic expansion with larval growth and nutrient availability. Thus, the modular TCA represents a cell-intrinsic layer of a larger, organism-wide nutrient-sensing circuit and better integration with systemic and niche signals.

The modular-to-loop transition in the lymph gland may reflect an evolutionarily conserved metabolic logic of stemness. Across metazoans, early progenitors often rely on incomplete TCA activity, favoring flexibility and redox control, while differentiated cells employ full oxidative cycling for stability (130–132). Our findings extend this logic by showing that a tissue can reconstruct the TCA topology developmentally to couple proliferation, differentiation, and organ scaling, a potential ancestral feature of metabolic control during organogenesis.

While RNA*i* based perturbations establish functional modules, quantitative flux analysis and metabolite imaging will be needed to resolve the dynamics of citrate and α-KG across zones. Single-cell metabolomics or isotope tracing approaches could reveal how metabolite gradients evolve and whether inter-zonal shuttling occurs through transporters such as Indy/SLC13A5. Integrating chromatin accessibility and metabolic flux measurements will test how TCA topology directly interfaces with transcriptional and epigenetic programs during progenitor maturation.

Collectively, our data support a model where zonal communication through metabolites coordinates developmental progression: The MZ generates citrate/isocitrate to sustain progenitor identity and provides them with a long term proliferative capacity, α-KG to drive early proliferation. This modular later transitions to a complete TCA cycle operation to maintain metabolic balance and progenitor homeostasis. The CZ exhibits partial TCA activity, and the IZ integrates metabolic signals from both zones, toggling between modular and complete TCA operation to maintain metabolic balance and developmental timing between progenitor maintenance and differentiation (Fig. S10). This zonal interplay establishes the IZ as a rate-limiting node for hematopoietic homeostasis sensing systemic metabolite buildup and activating full TCA flux to stabilize progenitor differentiation balance. Such a model underscores how spatially distributed metabolic programs can coordinate organ-level development (Fig. 6).

By demonstrating how a hematopoietic tissue developmentally rewires the TCA from modular to cyclic and by linking specific nodes such as citrate, α-KG, and succinyl-CoA to progenitor identity, proliferation, and epigenetic control, this study offers an insight into HSC biology that may be largely relevant not only in mammalian HSCs, but across stem/progenitor cell systems. The study provides a conceptual platform and elucidates how TCA architecture governs stemness, lineage commitment, and organ-level homeostasis.

## Materials and Methods

### *Drosophila* husbandry, stocks and genetics

The following *Drosophila* stocks were received as gifts from Banerjee lab, UCLA, USA: *w^1118^* (control), d*omeMESO-Gal4,UAS-GFP, ChIZ-GAL4, Hml-GAL4,UAS-2X EGFP, and Tep4-GAL4,UASmcherry*. The *RNAi* stocks used in this study were obtained from Vienna *Drosophila* Resource Centre (VDRC) and Bloomington *Drosophila* Stock Centre (BDSC), these stocks are: *kdn^RNAi^/CS^RNAi^*(BDSC 60900), *mAcon1^RNAi^* (BDSC 34028), *Idh^RNAi^* (BDSC 41708, 56203)*, α-Kdh^RNAi^ (*BDSC 34101, 33686)*, skap^RNAi^* (BDSC 55168, 50939), *SdhA^RNAi^* (VDRC 330053), *Fum4^RNAi^* (BDSC 65195)*, Mdh2^RNAi^* (BDSC 62230, 62228; VDRC 101551*) Pdh^RNAi^* (BDSC 55345)*, Gdh^RNAi^* (BDSC 53255). All fly stocks were reared on corn meal agar medium with yeast supplementation at 25°C incubator unless specified. The crosses involving RNA*i* lines were maintained at 29°C to maximize the efficacy of the *Gal4/UAS* RNA*i* system. Controls refers to either *w^1118^* (wild type) or Gal4 drivers crossed with *w^1118^*.

### Immunostaining and immunohistochemistry

For staining blood cell progenitors, 3rd instar larval lymph glands were dissected out in 1X PBS and fixed in 4% formaldehyde solution, for 15 min. Post-fixation the tissues were permeabilized by 3 washes of 15 min each in 0.3% PBST (0.3% Triton-X in 1X PBS). Tissues were then blocked in 5% NGS (Normal Goat Serum), followed by overnight primary antibody treatment. Next day tissues were washed (as earlier) and incubated in secondary antibody for 2-3 hours, followed by washing (as earlier) and mounting in mounting media.

Immunohistochemistry on lymph gland was performed with the following primary antibodies: mouse α-P1 (1:30, I. Ando), rabbit α-Pxn (1:1000, J. Shim), mouse α-Myo (1:100, DSHB), rabbit α-PPO (1:200, H. M. Müller), and α-DHE (1:1000, Life technologies), rabbit α-pH3 (1:100; CST 3642S), rabbit α-Caspase 3 (1:200; CST 9661S). All the secondary antibodies were used at 1:500 dilutions, those are FITC, Cy3 and Cy5 (Jackson Immuno Research Laboratories) and Alexa Fluor 488, 546, 647 (Invitrogen). Nuclei were visualized using DAPI/Hoechst-33342 (Sigma B2261). Samples were mounted in mounting media (Vectashield, Vector Laboratories).

### Imaging and quantification of lymph gland phenotypes

Immuno-stained images of lymph glands were acquired using Olympus FV3000 confocal microscopy on 40X and 60X oil-immersion objective. Image was scanned with frame size of 800 X 800 with 0.5/1/2um thickness for Z-stack, depending upon the experiment.

All images were quantified using ImageJ software (NIH, USA). The details of the quantifications carried out are as under:

### Lymph gland and Zone area analysis

Roughly, middle two confocal Z-stacks were merged and area was selected according to the respective stainings and measured with the help of freehand selection tool of ImageJ software, followed by using the “measure tool” under “analysis tool” to get the area values. This was done for respective zones, where Tep4 or Dome area represents the progenitor zone, Dome/Pxn area represent differentiating or intermediary zone, Pxn/P1 and Hml/P1 area represent differentiated zone. The areas were represented in percent values with respect to the total area of the respective lobe. Controls were analysed in parallel to the tests every time. A minimum of 10-15 animals were analysed each time and the experiment was repeated at least three times. The quantifications represent the mean (% area) of the independent experimental sets.

### Crystals cells and lamellocytes

Crystal cell population were analysed by counting total number of cell across the Z-stacks by merging all the stacks of lymph gland. These cell counts are represented as total number of cells per lymph gland area. LG lobes with lamellocytes induction were counted against the total number of LG lobes, for quantification.

### Total, Dome^+^, and Dome^-^ nuclei analysis

To count the total nuclei, we used the ImageJ software (NIH, USA). All the stacks of the lymph gland were merged with the help of “Z stack tool” and the primary lobe was outlined by the freehand selection tool, followed by “clear outside tool” under “edit option” to remove the other tissues (e.g. ring gland, secondary, tertiary lobes, dorsal vessel etc.). The nuclei of the remaining primary lobe were then thresholded using “threshold tool” under “Image-adjust option” to obtain 8-bit image in single channel. The image obtained is then subjected to “watershed tool” under “Process-Binary option” followed by “particle analysis” with particle size 3-infinity and circularity of 0.04-1.00. This gave the total number of particles (nuclei) which were marked by Hoechst dye, in blue channel. Similarly, for Dome+ nuclei only green channel was subjected to this mentioned processing and for the Dome-nuclei Dome+ nuclei were subtracted from total nuclei for the respective lobes.

### Mitotic index for Dome^+^ and Dome^-^ population

Total pH3+ cells were counted by merging all the Z-stack together to get total number of pH3+ cells in the entire lobe. For assessing pH3^+^ population in Dome^+^ zone, *domeGFP* channel was overlaid onto pH3 channel and for the Dome^-^ pH3^+^ cells Dome^+^ pH3^+^ cells were subtracted from total pH3^+^ cells. For calculating mitotic index pH3^+^ cells were divided by total number of cells per LG lobe and same is done for Dome^+^ and Dome^-^ mitotic index.

### SABER FISH

Reagents: Quick-load 100bp DNA ladder (N0467S; NEB), SYBR Gold Nucleic Acid Gel Stain (S11494; Invitrogen), *Bst* DNA Polymerase, Large fragment (M0275L; NEB), Deoxynucleotide (dNTP) Solution set (N0446S; NEB), Formamide (AM9342; Invitrogen), Triton X-100 (T8787-50ML; Sigma Aldrich), Dextran sulfate sodium (D8906-10G; Sigma Aldrich), TWEEN^®^ 20 (P9416-50ML; Sigma Aldrich), Propyl gallate (P3130-100G; Sigma Aldrich), Magnesium Sulfate (MgSO4) Solution (B1003S; NEB), UltraPure DNase/RNase-Free Distilled Water, PBS -Phosphate-Buffered Saline (10X) pH 7.4, RNase-free (AM9624; Invitrogen), UltraPure SSC, 20X (15557044; Invitrogen).

The protocol was adapted from Kishi and coworkers, with slight modification wherever needed according to the *Drosophila* tissue (133). The protocol is as under:

#### DAY1

Setting up of PER reaction: Add all the reagents to the tube, EXCEPT for probes or primers and keep the tube at 37°C for enzyme activation for 15min. Add specific probes to the tubes and incubate for 2h/3h depending upon the hairpin in use. Terminate the reaction at 80°C for 20min and then the extended probes can be stored at 4°C for years. Confirm the extension by 1% agarose gel

#### DAY2

Dissected the LG of 48h, 72h and 96h in PBS in NUNC 4 well dish. Fixed the tissues with 4% PFA (Formaldehyde does not work very well in this protocol) for 15 min at room temperature (RT). Tissue permeabalization: 0.3% PBST (Triton X) was for 15 min at RT. Washing: 0.1% PBSTw (Tween-20) (3 X 5min at RT). Primary Hybridisation: previous day prepared/extended probes along with hybridisation solution, in a humid chamber for overnight (16h) at 45°C.

#### DAY3

Wash with Tissue wash buffer-A (40% Formamide) (2 X 20min at 37°C). Wash with Tissue wash buffer-B (25% Formamide) (2 X 30min at 37°C). Wash with 2X SSCT buffer (2 X 15min at 37°C). Developing: Add Hybridisation solution-2 (fluorophores) and incubate for 2-3h at 37°C. Wash with PBSTw (3 X 15 min at 37°C). Mount the samples in vectasheild and image.

### Metabolite supplementation

Supplemented food was prepared by adding metabolites (1% citrate (Sodium citrate tribasic dihydrate, Sigma, C7254) and 3% succinate (Sodium succinate dibasic hexahydrate, Sigma, S9637; weight/volume)) to the regular fly food. Eggs or larvae of respective stages were transferred to the supplemented food according to the experimental requirements for lymph gland analysis. In early feeding only the larvae were transferred back to the regular food after rearing on supplemented food for further development. Controls for the respective experiments were always treated similarly, un-supplemented controls were always kept alongside.

### Software, sample size, and Statistical analyses

All the images were processed in Imagej (Fiji) software and the figure panels were prepared in Microsoft power point 2010 and Adobe Photoshop. Data post-calculation was stored in Microsoft excel 2010. All models were prepared in Biorender software. OpenAI (ChatGPT) was used for the grammatical corrections and text rearranging. In all experiments, *n* implies the total number of samples analyzed that were obtained from multiple independent experimental repeats and ‘*N*’ represents the number of independent experimental repeats. All experiments have been repeated a minimum of three times; in each experimental setup, at least 10-15 animals were analyzed. *Drosophila* availability is not limiting; therefore, no power calculations were used to predetermine sample size. All statistical analyses were performed using GraphPad Prism 10 software. The means were analysed using ordinary one-way ANNOVA or unpaired Student t-test (Mann-Whitney test). For all the experiments batch effect were plotted. Plots, test applied and sample size is mentioned separately in the figure legends section.

## Supporting information

Supplementary images and supplementary table

## Acknowledgments

We thank Prof. I. Ando for the NimC1 (P1) antibody, Prof. H. M. Müller for the PPO antibody, and Prof. J. Shim for the Pxn antibody. The Bloomington *Drosophila* Stock Center (BDSC) and Vienna *Drosophila* Resource Center (VDRC) for fly stocks, the FlyBase for the literature. We acknowledge National Centre for Biological Sciences (NCBS), Centre for Cellular and Molecular Platforms (C CAMP) for Central Imaging & Flow Cytometry Facility (CIFF) and the fly facility. Owing to space limitations, we apologize to our colleagues whose work is not cited. Schematics/models were created with BioRender.com and PowerPoint 2016. Graphs were made in GraphPad Prism 10 software and OpenAI (ChatGPT) was used for the grammatical corrections and text rearranging. All the image quantification was done on ImageJ and figure panels were made in Adobe Photoshop. This study was supported by DBT/Wellcome Trust India Alliance Senior Research Fellowship (Grant number IA/S/22/1/506259), CNRS-International Research Project (IRP) MACHUB awarded to T.M. This study was partly also supported by SERB-ANRF (Grant no: SPR/2021/000486) awarded to S.C. A.T. is a Graduate Student at inStem, in the Mukherjee lab, and is supported by University Grant Commission (UGC)-Fellowship.

## Author Contributions

Conceptualization: A.T., T.M.; Methodology: A.T.; Validation: A.T.; Formal analysis: A.T.; Investigation: A.T.; Resources: A.T., S.C.; Data curation: A.T.; Writing -original draft: A.T., T.M.; Writing -review & editing: M.G., T.M.; Visualization: M.G.; Supervision: T.M.; Project administration: T.M.; Funding acquisition: S.C., T.M.

## Classification

Major Classification: Biological Sciences

Minor Classification: Developmental Biology; Immunology

## References

1. B. W. Carey, L. W. S. Finley, J. R. Cross, C. D. Allis, C. B. Thompson, Intracellular α-ketoglutarate maintains the pluripotency of embryonic stem cells. Nature 518, 413–416 (2015).

2. G. M. Tannahill, et al., Succinate is an inflammatory signal that induces IL-1β through HIF-1α. Nature 496, 238–242 (2013).

3. K. E. Wellen, et al., ATP-citrate lyase links cellular metabolism to histone acetylation. Science 324, 1076–1080 (2009).

4. M. Xiao, et al., Inhibition of α-KG-dependent histone and DNA demethylases by fumarate and succinate that are accumulated in mutations of FH and SDH tumor suppressors. Genes Dev. 26, 1326–1338 (2012).

5. H. Gu, et al., MDH1-mediated malate-aspartate NADH shuttle maintains the activity levels of fetal liver hematopoietic stem cells. Blood 136, 553–571 (2020).

6. C. B. Edwards, N. Copes, A. G. Brito, J. Canfield, P. C. Bradshaw, Malate and Fumarate Extend Lifespan in Caenorhabditis elegans. PLOS ONE 8, e58345 (2013).

7. M. Yang, T. Soga, P. J. Pollard, J. Adam, The emerging role of fumarate as an oncometabolite. Front. Oncol. 2 (2012).

8. I. Martínez-Reyes, N. S. Chandel, Mitochondrial TCA cycle metabolites control physiology and disease. Nat. Commun. 11, 102 (2020).

9. H. M. Wilkins, et al., Oxaloacetate activates brain mitochondrial biogenesis, enhances the insulin pathway, reduces inflammation and stimulates neurogenesis. Hum. Mol. Genet. 23, 6528–6541 (2014).

10. D. S. Williams, A. Cash, L. Hamadani, T. Diemer, Oxaloacetate supplementation increases lifespan in Caenorhabditis elegans through an AMPK/FOXO-dependent pathway. Aging Cell 8, 765–768 (2009).

11. C. Lussey-Lepoutre, et al., Loss of succinate dehydrogenase activity results in dependency on pyruvate carboxylation for cellular anabolism. Nat. Commun. 6, 8784 (2015).

12. C. M. Metallo, et al., Reductive glutamine metabolism by IDH1 mediates lipogenesis under hypoxia. Nature 481, 380–384 (2011).

13. A. R. Mullen, et al., Reductive carboxylation supports growth in tumour cells with defective mitochondria. Nature 481, 385–388 (2011).

14. P. K. Arnold, et al., A non-canonical tricarboxylic acid cycle underlies cellular identity. Nature 603, 477–481 (2022).

15. N. Assmann, et al., Srebp-controlled glucose metabolism is essential for NK cell functional responses. Nat. Immunol. 18, 1197–1206 (2017).

16. P. Borst, “Hydrogen transport and transport metabolites” in Funktionelle und Morphologische Organisation der Zelle, (Springer Berlin Heidelberg, 1963), pp. 137–162.

17. R. A. Parlo, P. S. Coleman, Enhanced rate of citrate export from cholesterol-rich hepatoma mitochondria. The truncated Krebs cycle and other metabolic ramifications of mitochondrial membrane cholesterol. J. Biol. Chem. 259, 9997– 10003 (1984).

18. V. Lampropoulou, et al., Itaconate Links Inhibition of Succinate Dehydrogenase with Macrophage Metabolic Remodeling and Regulation of Inflammation. Cell Metab. 24, 158–166 (2016).

19. M. P. Murphy, L. A. J. O’Neill, Krebs Cycle Reimagined: The Emerging Roles of Succinate and Itaconate as Signal Transducers. Cell 174, 780–784 (2018).

20. V. Infantino, et al., The mitochondrial citrate carrier: a new player in inflammation. Biochem. J. 438, 433–436 (2011).

21. V. Infantino, V. Iacobazzi, F. Palmieri, A. Menga, ATP-citrate lyase is essential for macrophage inflammatory response. Biochem. Biophys. Res. Commun. 440, 105– 111 (2013).

22. L. Shi, B. P. Tu, Acetyl-CoA and the regulation of metabolism: mechanisms and consequences. Curr. Opin. Cell Biol. 33, 125–131 (2015).

23. S. Sivanand, I. Viney, K. E. Wellen, Spatiotemporal Control of Acetyl-CoA Metabolism in Chromatin Regulation. Trends Biochem. Sci. 43, 61–74 (2018).

24. J. V. Lee, et al., Akt-dependent metabolic reprogramming regulates tumor cell histone acetylation. Cell Metab. 20, 306–319 (2014).

25. P.-S. Liu, et al., α-ketoglutarate orchestrates macrophage activation through metabolic and epigenetic reprogramming. Nat. Immunol. 18, 985–994 (2017).

26. D. G. Ryan, L. A. J. O’Neill, Krebs cycle rewired for macrophage and dendritic cell effector functions. FEBS Lett. 591, 2992–3006 (2017).

27. P. Gao, et al., Capillary electrophoresis -Mass spectrometry metabolomics analysis revealed enrichment of hypotaurine in rat glioma tissues. Anal. Biochem. 537, 1–7 (2017).

28. K. J. Harber, et al., Succinate Is an Inflammation-Induced Immunoregulatory Metabolite in Macrophages. Metabolites 10, 372 (2020).

29. D. G. Ryan, L. A. J. O’Neill, Krebs Cycle Reborn in Macrophage Immunometabolism. Annu. Rev. Immunol. 38, 289–313 (2020).

30. G. L. Semenza, Hypoxia-inducible factors in physiology and medicine. Cell 148, 399–408 (2012).

31. E. L. Mills, et al., Succinate Dehydrogenase Supports Metabolic Repurposing of Mitochondria to Drive Inflammatory Macrophages. Cell 167, 457–470.e13 (2016).

32. J.-Y. Wu, et al., Cancer-Derived Succinate Promotes Macrophage Polarization and Cancer Metastasis via Succinate Receptor. Mol. Cell 77, 213–227.e5 (2020).

33. M. A. Selak, et al., Succinate links TCA cycle dysfunction to oncogenesis by inhibiting HIF-alpha prolyl hydroxylase. Cancer Cell 7, 77–85 (2005).

34. A. K. Jha, et al., Network integration of parallel metabolic and transcriptional data reveals metabolic modules that regulate macrophage polarization. Immunity 42, 419–430 (2015).

35. B. Kelly, L. A. O’Neill, Metabolic reprogramming in macrophages and dendritic cells in innate immunity. Cell Res. 25, 771–784 (2015).

36. N. K. Patil, J. K. Bohannon, A. Hernandez, T. K. Patil, E. R. Sherwood, Regulation of leukocyte function by citric acid cycle intermediates. J. Leukoc. Biol. 106, 105– 117 (2019).

37. R. J. W. Arts, et al., Glutaminolysis and Fumarate Accumulation Integrate Immunometabolic and Epigenetic Programs in Trained Immunity. Cell Metab. 24, 807–819 (2016).

38. N. P. Riksen, M. G. Netea, Immunometabolic control of trained immunity. Mol. Aspects Med. 77, 100897 (2021).

39. C. D. C. C. van der Heijden, et al., Epigenetics and Trained Immunity. Antioxid. Redox Signal. 29, 1023–1040 (2018).

40. N. S. Chandel, et al., Mitochondrial reactive oxygen species trigger hypoxia-induced transcription. Proc. Natl. Acad. Sci. U. S. A. 95, 11715–11720 (1998).

41. N. S. Chandel, et al., Reactive oxygen species generated at mitochondrial complex III stabilize hypoxia-inducible factor-1alpha during hypoxia: a mechanism of O2 sensing. J. Biol. Chem. 275, 25130–25138 (2000).

42. E. Owusu-Ansah, U. Banerjee, Reactive oxygen species prime Drosophila haematopoietic progenitors for differentiation. Nature 461, 537–541 (2009).

43. W. G. Kaelin, P. J. Ratcliffe, Oxygen sensing by metazoans: the central role of the HIF hydroxylase pathway. Mol. Cell 30, 393–402 (2008).

44. D. A. Patten, et al., Hypoxia-inducible factor-1 activation in nonhypoxic conditions: the essential role of mitochondrial-derived reactive oxygen species. Mol. Biol. Cell 21, 3247–3257 (2010).

45. E. T. Chouchani, et al., Ischaemic accumulation of succinate controls reperfusion injury through mitochondrial ROS. Nature 515, 431–435 (2014).

46. C. M. Spratford, et al., Intermediate progenitor cells provide a transition between hematopoietic progenitors and their differentiated descendants. Development 148, dev200216 (2021).

47. S.-H. Jung, C. J. Evans, C. Uemura, U. Banerjee, The *Drosophila* lymph gland as a developmental model of hematopoiesis. Development 132, 2521–2533 (2005).

48. J. Krzemien, J. Oyallon, M. Crozatier, A. Vincent, Hematopoietic progenitors and hemocyte lineages in the Drosophila lymph gland. Dev. Biol. 346, 310–319 (2010).

49. H. Asha, et al., Analysis of Ras-Induced Overproliferation in Drosophila Hemocytes. Genetics 163, 203–215 (2003).

50. I. Anderl, et al., Transdifferentiation and Proliferation in Two Distinct Hemocyte Lineages in Drosophila melanogaster Larvae after Wasp Infection. PLoS Pathog. 12, e1005746 (2016).

51. C. J. Evans, T. Liu, U. Banerjee, *Drosophila* hematopoiesis: Markers and methods for molecular genetic analysis. Methods 68, 242–251 (2014).

52. L. I. Fessler, R. E. Nelson, J. H. Fessler, Drosophila extracellular matrix. Methods Enzymol. 245, 271–294 (1994).

53. A. Goto, et al., A Drosophila haemocyte-specific protein, hemolectin, similar to human von Willebrand factor. Biochem. J. 359, 99–108 (2001).

54. A. Goto, T. Kadowaki, Y. Kitagawa, Drosophila hemolectin gene is expressed in embryonic and larval hemocytes and its knock down causes bleeding defects. Dev. Biol. 264, 582–591 (2003).

55. R. E. Nelson, et al., Peroxidasin: a novel enzyme-matrix protein of Drosophila development. EMBO J. 13, 3438–3447 (1994).

56. S. Yasothornsrikul, W. J. Davis, G. Cramer, D. A. Kimbrell, C. R. Dearolf, viking: identification and characterization of a second type IV collagen in Drosophila. Gene 198, 17–25 (1997).

57. P. Bangs, K. White, Regulation and execution of apoptosis during Drosophila development. Dev. Dyn. 218, 68–79 (2000).

58. T. M. Rizki, R. M. Rizki, “The Cellular Defense System of Drosophila melanogaster” in Insect Ultrastructure*: Volume 2*, R. C. King, H. Akai, Eds. (Springer US, 1984), pp. 579–604.

59. U. Tepass, L. I. Fessler, A. Aziz, V. Hartenstein, Embryonic origin of hemocytes and their relationship to cell death in Drosophila. Dev. Camb. Engl. 120, 1829–1837 (1994).

60. R. Lanot, D. Zachary, F. Holder, M. Meister, Postembryonic hematopoiesis in Drosophila. Dev. Biol. 230, 243–257 (2001).

61. M. T. M. Rizki, Alterations in the haemocyte population of Drosophila melanogaster. J. Morphol. 100, 437–458 (1957).

62. O. Binggeli, C. Neyen, M. Poidevin, B. Lemaitre, Prophenoloxidase Activation Is Required for Survival to Microbial Infections in Drosophila. PLOS Pathog. 10, e1004067 (2014).

63. J. P. Dudzic, S. Kondo, R. Ueda, C. M. Bergman, B. Lemaitre, Drosophila innate immunity: regional and functional specialization of prophenoloxidases. BMC Biol. 13, 81 (2015).

64. P. Irving, et al., New insights into Drosophila larval haemocyte functions through genome-wide analysis. Cell. Microbiol. 7, 335–350 (2005).

65. H. Nam, I. Jang, H. You, K. Lee, W. Lee, Genetic evidence of a redox-dependent systemic wound response via Hayan Protease-Phenoloxidase system in Drosophila | The EMBO Journal. (2012). Available at: https://www.embopress.org/doi/full/10.1038/emboj.2011.476 [Accessed 1 May 2025].

66. A. J. Nappi, E. Vass, F. Frey, Y. Carton, Superoxide anion generation in Drosophila during melanotic encapsulation of parasites. Eur. J. Cell Biol. 68, 450–456 (1995).

67. T. M. Rizki, R. M. Rizki, Lamellocyte differentiation in Drosophila larvae parasitized by Leptopilina. Dev. Comp. Immunol. 16, 103–110 (1992).

68. M. Goyal, A. Tomar, S. Madhwal, T. Mukherjee, Blood progenitor redox homeostasis through olfaction-derived systemic GABA in hematopoietic growth control in Drosophila. Development 149, dev199550 (2021).

69. M. Goyal, et al., Metabolic coupling of ROS generation and antioxidant synthesis by the GABA shunt pathway in myeloid-like blood progenitor cells of Drosophila. PLOS Genet. 21, e1011602 (2025).

70. J. Shim, T. Mukherjee, U. Banerjee, Direct sensing of systemic and nutritional signals by hematopoietic progenitors in Drosophila. Nat. Cell Biol. 14, 394–400 (2012).

71. S. A. Sinenko, L. Mandal, J. A. Martinez-Agosto, U. Banerjee, Dual Role of Wingless Signaling in Stem-like Hematopoietic Precursor Maintenance in Drosophila. Dev. Cell 16, 756–763 (2009).

72. A. Prakash, M. S. Inamdar, Mitochondrial-activity-driven hematopoietic stem cell fate and lineage choice is first established in the aorta-gonad-mesonephros. Cell Rep. 44, 116296 (2025).

73. H. Daud, S. Browne, R. Al-Majmaie, W. Murphy, M. Al-Rubeai, Metabolic profiling of hematopoietic stem and progenitor cells during proliferation and differentiation into red blood cells. New Biotechnol. 33, 179–186 (2016).

74. A. Sinha, R. J. Khadilkar, V. K. S., A. RoyChowdhury Sinha, M. S. Inamdar, Conserved Regulation of the JAK/STAT Pathway by the Endosomal Protein Asrij Maintains Stem Cell Potency. Cell Rep. 4, 649–658 (2013).

75. S. K. Tiwari, A. G. Toshniwal, S. Mandal, L. Mandal, Fatty acid β-oxidation is required for the differentiation of larval hematopoietic progenitors in Drosophila. eLife 9, e53247 (2020).

76. M.-D. Filippi, S. Ghaffari, Mitochondria in the maintenance of hematopoietic stem cells: new perspectives and opportunities. Blood 133, 1943–1952 (2019).

77. M. Xiong, et al., Proteomics reveals dynamic metabolic changes in human hematopoietic stem progenitor cells from fetal to adulthood. Stem Cell Res. Ther. 15, 303 (2024).

78. P. Irving, et al., New insights into Drosophila larval haemocyte functions through genome-wide analysis. Cell. Microbiol. 7, 335–350 (2005).

79. J. Krzemień, et al., Control of blood cell homeostasis in Drosophila larvae by the posterior signalling centre. Nature 446, 325–328 (2007).

80. J. R. Girard, et al., Paths and pathways that generate cell-type heterogeneity and developmental progression in hematopoiesis. eLife 10, e67516 (2021).

81. U. Banerjee, J. R. Girard, L. M. Goins, C. M. Spratford, *Drosophila* as a Genetic Model for Hematopoiesis. Genetics 211, 367–417 (2019).

82. A. Kapoor, A. Padmavathi, S. Madhwal, T. Mukherjee, Dual control of dopamine in Drosophila myeloid-like progenitor cell proliferation and regulation of lymph gland growth. EMBO Rep. 23, e52951 (2022).

83. I. Louradour, et al., Reactive oxygen species-dependent Toll/NF-κB activation in the Drosophila hematopoietic niche confers resistance to wasp parasitism. eLife 6, e25496 (2017).

84. T. Tokusumi, et al., Screening and Analysis of Janelia FlyLight Project Enhancer-Gal4 Strains Identifies Multiple Gene Enhancers Active During Hematopoiesis in Normal and Wasp-Challenged Drosophila Larvae. G3 GenesGenomesGenetics 7, 437–448 (2017).

85. S. Korkes, J. R. Stern, I. C. Gunsalus, S. Ochoa, Enzymatic Synthesis of Citrate from Pyruvate and Oxalacetate. Nature 166, 439–440 (1950).

86. R. C. Pletcher, et al., A Genetic Screen Using the Drosophila melanogaster TRiP RNAi Collection To Identify Metabolic Enzymes Required for Eye Development. G3 Bethesda Md 9, 2061–2070 (2019).

87. R. J. DeBerardinis, J. J. Lum, G. Hatzivassiliou, C. B. Thompson, The Biology of Cancer: Metabolic Reprogramming Fuels Cell Growth and Proliferation. Cell Metab. 7, 11–20 (2008).

88. O. E. Owen, S. C. Kalhan, R. W. Hanson, The key role of anaplerosis and cataplerosis for citric acid cycle function. J. Biol. Chem. 277, 30409–30412 (2002).

89. P. K. Arnold, L. W. S. Finley, Regulation and function of the mammalian tricarboxylic acid cycle. J. Biol. Chem. 299, 102838 (2023).

90. H. Yoo, M. R. Antoniewicz, G. Stephanopoulos, J. K. Kelleher, Quantifying Reductive Carboxylation Flux of Glutamine to Lipid in a Brown Adipocyte Cell Line. J. Biol. Chem. 283, 20621–20627 (2008).

91. B. Cho, et al., Single-cell transcriptome maps of myeloid blood cell lineages in Drosophila. Nat. Commun. 11, 4483 (2020).

92. A. Ray, K. Kamat, M. S. Inamdar, A Conserved Role for Asrij/OCIAD1 in Progenitor Differentiation and Lineage Specification Through Functional Interaction With the Regulators of Mitochondrial Dynamics. Front. Cell Dev. Biol. 9 (2021).

93. Y. S. Cho, et al., Discovery and Evaluation of Clinical Candidate IDH305, a Brain Penetrant Mutant IDH1 Inhibitor. ACS Med. Chem. Lett. 8, 1116–1121 (2017).

94. D. C. Singleton, et al., Pyruvate anaplerosis is a mechanism of resistance to pharmacological glutaminase inhibition in triple-receptor negative breast cancer. BMC Cancer 20, 470 (2020).

95. R. Leonardi, C. Subramanian, S. Jackowski, C. O. Rock, Cancer-associated Isocitrate Dehydrogenase Mutations Inactivate NADPH-dependent Reductive Carboxylation*. J. Biol. Chem. 287, 14615–14620 (2012).

96. S. K. Sharma, S. Ghosh, A. R. Geetha, S. Mandal, L. Mandal, Cell Adhesion-Mediated Actomyosin Assembly Regulates the Activity of Cubitus Interruptus for Hematopoietic Progenitor Maintenance in Drosophila. Genetics 212, 1279–1300 (2019).

97. S. Madhwal, et al., Metabolic control of cellular immune-competency by odors in Drosophila. eLife 9, e60376 (2020).

98. F. Kocabas, et al., Meis1 regulates the metabolic phenotype and oxidant defense of hematopoietic stem cells. Blood 120, 4963–4972 (2012).

99. M. Maryanovich, et al., An MTCH2 pathway repressing mitochondria metabolism regulates haematopoietic stem cell fate. Nat. Commun. 6, 7901 (2015).

100. L. Papa, M. Djedaini, R. Hoffman, Mitochondrial Role in Stemness and Differentiation of Hematopoietic Stem Cells. Stem Cells Int. 2019, 4067162 (2019).

101. C. Morganti, N. Cabezas-Wallscheid, K. Ito, Metabolic Regulation of Hematopoietic Stem Cells. HemaSphere 6, e740 (2022).

102. G. Hajishengallis, X. Li, T. Chavakis, Immunometabolic control of hematopoiesis. Mol. Aspects Med. 77, 100923 (2021).

103. S. Watanuki, et al., Context-dependent modification of PFKFB3 in hematopoietic stem cells promotes anaerobic glycolysis and ensures stress hematopoiesis. eLife 12, RP87674 (2024).

104. M. Xiong, et al., Proteomics reveals dynamic metabolic changes in human hematopoietic stem progenitor cells from fetal to adulthood. Stem Cell Res. Ther. 15, 303 (2024).

105. G. Hatzivassiliou, et al., ATP citrate lyase inhibition can suppress tumor cell growth. Cancer Cell 8, 311–321 (2005).

106. M. Peng, et al., Aerobic glycolysis promotes T helper 1 cell differentiation through an epigenetic mechanism. Science 354, 481–484 (2016).

107. A. Yalcin, S. Telang, B. Clem, J. Chesney, Regulation of glucose metabolism by 6-phosphofructo-2-kinase/fructose-2,6-bisphosphatases in cancer. Exp. Mol. Pathol. 86, 174–179 (2009).

108. R. V. Durán, et al., Glutaminolysis Activates Rag-mTORC1 Signaling. Mol. Cell 47, 349–358 (2012).

109. J. G. Moloughney, et al., mTORC2 Responds to Glutamine Catabolite Levels to Modulate the Hexosamine Biosynthesis Enzyme GFAT1. Mol. Cell 63, 811–826 (2016).

110. Y. Sekita, et al., AKT signaling is associated with epigenetic reprogramming via the upregulation of TET and its cofactor, alpha-ketoglutarate during iPSC generation. Stem Cell Res. Ther. 12, 510 (2021).

111. Y. W. Zhang, K. Schönberger, N. Cabezas-Wallscheid, Bidirectional interplay between metabolism and epigenetics in hematopoietic stem cells and leukemia. EMBO J. 42, e112348 (2023).

112. S. C. Baksh, L. W. S. Finley, Metabolic Coordination of Cell Fate by α-Ketoglutarate-Dependent Dioxygenases. Trends Cell Biol. 31, 24–36 (2021).

113. K. Liu, J. Cao, X. Shi, L. Wang, T. Zhao, Cellular metabolism and homeostasis in pluripotency regulation. Protein Cell 11, 630–640 (2020).

114. A. Ludin, et al., Reactive Oxygen Species Regulate Hematopoietic Stem Cell Self-Renewal, Migration and Development, As Well As Their Bone Marrow Microenvironment. Antioxid. Redox Signal. 21, 1605–1619 (2014).

115. Y.-Y. Jang, S. J. Sharkis, A low level of reactive oxygen species selects for primitive hematopoietic stem cells that may reside in the low-oxygenic niche. Blood 110, 3056–3063 (2007).

116. C. Piccoli, et al., Bone-marrow derived hematopoietic stem/progenitor cells express multiple isoforms of NADPH oxidase and produce constitutively reactive oxygen species. Biochem. Biophys. Res. Commun. 353, 965–972 (2007).

117. Y. Cao, et al., ROS functions as an upstream trigger for autophagy to drive hematopoietic stem cell differentiation. Hematol. Amst. Neth. 21, 613–618 (2016).

118. R. M. Denton, Regulation of mitochondrial dehydrogenases by calcium ions. Biochim. Biophys. Acta 1787, 1309–1316 (2009).

119. M. Destalminil-Letourneau, I. Morin-Poulard, Y. Tian, N. Vanzo, M. Crozatier, The vascular niche controls Drosophila hematopoiesis via fibroblast growth factor signaling. eLife 10, e64672 (2021).

120. F. Koranteng, B. Cho, J. Shim, Intrinsic and Extrinsic Regulation of Hematopoiesis in *Drosophila*. Mol. Cells 45, 101–108 (2022).

121. W. Lan, S. Liu, L. Zhao, Y. Su, Regulation of Drosophila Hematopoiesis in Lymph Gland: From a Developmental Signaling Point of View. Int. J. Mol. Sci. 21, 5246 (2020).

122. L. A. J. O’Neill, E. J. Pearce, Immunometabolism governs dendritic cell and macrophage function. J. Exp. Med. 213, 15–23 (2016).

123. B. Benmimoun, C. Polesello, L. Waltzer, M. Haenlin, Dual role for Insulin/TOR signaling in the control of hematopoietic progenitor maintenance in Drosophila. Development 139, 1713–1717 (2012).

124. N. S. Dey, P. Ramesh, M. Chugh, S. Mandal, L. Mandal, Dpp dependent Hematopoietic stem cells give rise to Hh dependent blood progenitors in larval lymph gland of Drosophila. eLife 5, e18295 (2016).

125. R. J. Khadilkar, et al., ARF1–GTP regulates Asrij to provide endocytic control of *Drosophila* blood cell homeostasis. Proc. Natl. Acad. Sci. 111, 4898–4903 (2014).

126. B. C. Mondal, et al., Interaction between Differentiating Cell-and Niche-Derived Signals in Hematopoietic Progenitor Maintenance. Cell 147, 1589–1600 (2011).

127. L. Mandal, J. A. Martinez-Agosto, C. J. Evans, V. Hartenstein, U. Banerjee, A Hedgehog- and Antennapedia-dependent niche maintains Drosophila haematopoietic precursors. Nature 446, 320–324 (2007).

128. D. Pennetier, et al., Size control of the *Drosophila* hematopoietic niche by bone morphogenetic protein signaling reveals parallels with mammals. Proc. Natl. Acad. Sci. 109, 3389–3394 (2012).

129. Y. Tokusumi, T. Tokusumi, D. A. Shoue, R. A. Schulz, Gene Regulatory Networks Controlling Hematopoietic Progenitor Niche Cell Production and Differentiation in the Drosophila Lymph Gland. PLoS ONE 7, e41604 (2012).

130. K. Takubo, et al., Regulation of glycolysis by Pdk functions as a metabolic checkpoint for cell cycle quiescence in hematopoietic stem cells. Cell Stem Cell 12, 49–61 (2013).

131. M. Khacho, et al., Mitochondrial Dynamics Impacts Stem Cell Identity and Fate Decisions by Regulating a Nuclear Transcriptional Program. Cell Stem Cell 19, 232–247 (2016).

132. M. J. Rodríguez-Colman, et al., Interplay between metabolic identities in the intestinal crypt supports stem cell function. Nature 543, 424–427 (2017).

133. J. Y. Kishi, et al., SABER amplifies FISH: enhanced multiplexed imaging of RNA and DNA in cells and tissues. Nat. Methods 16, 533–544 (2019).

