## Supplementary images and supplementary table for "Modular to cyclic TCA governs hematopoiesis in *Drosophila*"

Supplementary figure 1:

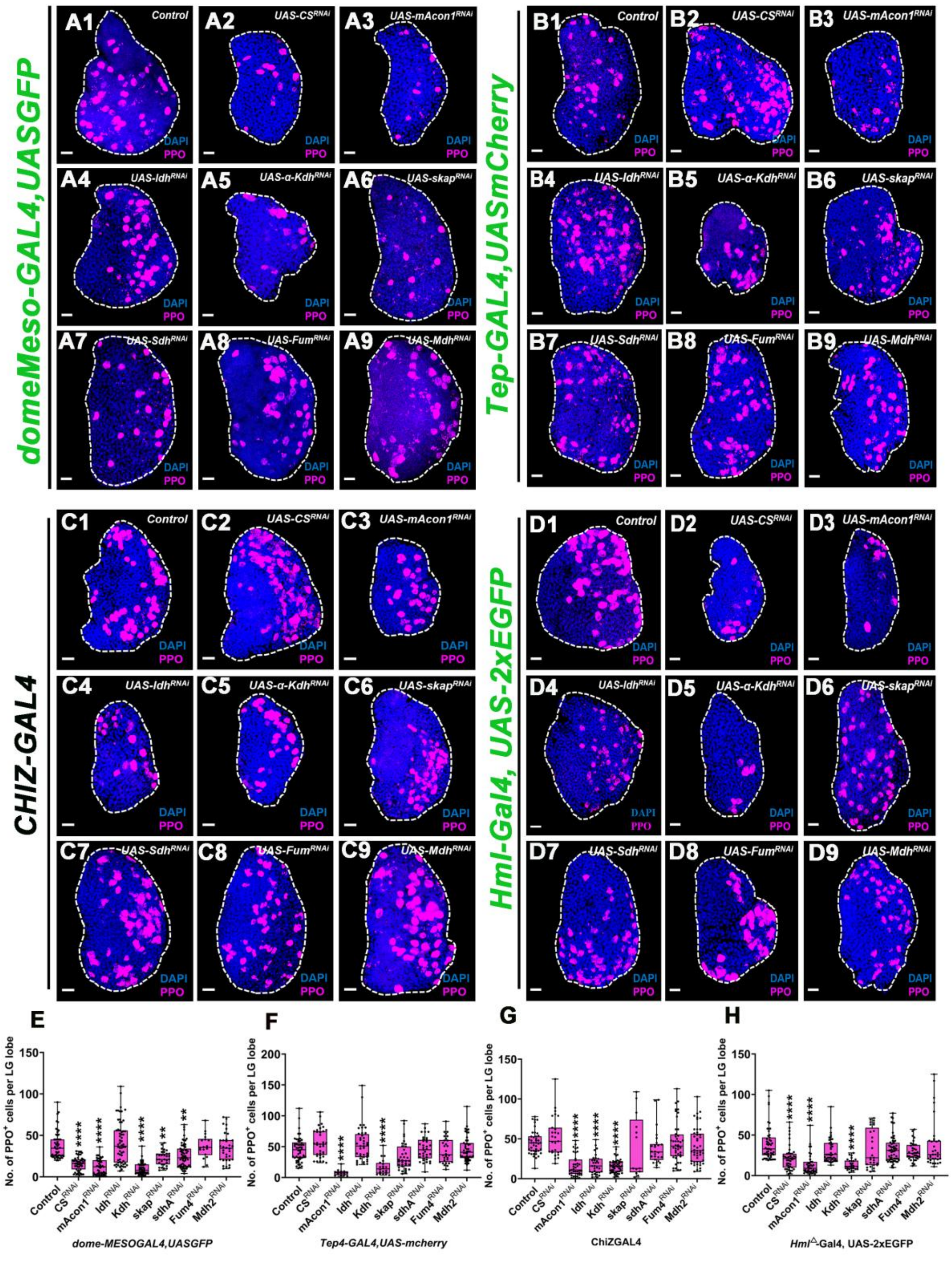

Supplementary Figure 1:

**(A1-A9)** Representative images showing crystal cells in the lymph gland of 3$^{rd}$ instar *Drosophila* larva, when TCA cycle was modulated in progenitor population (MZ) with the help of *domeMeso-Gal4,UAS-GFP.* **(A1)** *Control* (*domeMeso-Gal4,UAS-GFP/+*) lymph gland showing general distribution of mature crystal cells (PPO+; magenta) population at 3rd instar larval stage, **(A2)** expressing *CS*$^{RNAi}$ (*domeMeso-Gal4,UAS-GFP;UAS-CS*$^{RNAi}$), **(A3)** *mAcon1RNAi (domeMeso-Gal4,UAS-GFP;UAS-mAcon1*$^{RNAi}$), **(A4)** *α-Kdh*$^{RNAi}$ (*domeMeso-Gal4,UAS-GFP;UAS-α-Kdh*$^{RNAi}$), **(A5)** *Idh*$^{RNAi}$ (*domeMeso-Gal4,UAS-GFP;UAS-Idh*$^{RNAi)}$, **(A6)** *Skap*$^{RNAi}$ (*domeMeso-Gal4,UAS-GFP;UAS-Skap*$^{RNAi}$), **(A7)** *Sdh*$^{RNAi}$ (*domeMeso-Gal4,UAS-GFP;UAS-Sdh*$^{RNAi}$), **(A8)** *Fum*$^{RNAi}$ (*domeMeso-Gal4,UAS-GFP;UAS-Fum*$^{RNAi}$),and **(A9)** *Mdh*$^{RNAi}$ (*domeMeso-Gal4,UAS-GFP;UAS-Mdh*$^{RNAi}$), **(A2, A3 and A5-A7)** lymph glands shows reduction in crystal cells (PPO$^{+}$; magenta), compare to **(A1)** control. **(E)** Quantification is total number of crystal cells (PPO+) cells in entire LG lobe, *domeMeso>GFP/+* (*control*, n=50), *domeMeso>GFP/CS*$^{RNAi}$ (n=47, p<0.0001), *domeMeso>GFP/mAcon1*$^{RNAi}$ (n=47, p<0.0001), *domeMeso>GFP/Idh*$^{RNAi}$ (n=60, p=0.3933), *domeMeso>GFP/α-Kdh*$^{RNAi}$ (n=58, p<0.0001), *domeMeso>GFP/Skap*$^{RNAi}$ (n=28, p=0.0020), *domeMeso>GFP/Sdh*$^{RNAi}$ (n=57, p=0.0063), *domeMeso>GFP/Fum*$^{RNAi}$ (n=18, p>0.9999), *domeMeso>GFP/Mdh*$^{RNAi}$ (n=34, p=0.9997). **(B1-B9)** Representative images showing crystal cells in the lymph gland of 3$^{rd}$ instar *Drosophila* larva, when TCA cycle was modulated in core-progenitor population (MZ) with the help of *Tep4-GAL4,UASmcherry.* **(B1)** *Control* (*Tep4-GAL4,UASmcherry/+*) **(B2)** CS$^{RNAi}$ (*Tep4-GAL4,UASmcherry;UAS-CS*$^{RNAi}$), **(B3)** *mAcon1*$^{RNAi}$ (*Tep4-GAL4,UASmcherry;UAS-mAcon1RNAi*), **(B4)** *Idh*$^{RNAi}$ *(Tep4-GAL4,UASmcherry;UAS-Idh*$^{RNAi}$*),* **(B5)** *α-Kdh*$^{RNAi}$ (*Tep4-GAL4,UASmcherry;UAS-α-KdhRNAi*), ***(B6)*** *Skap*$^{RNAi}$ *(Tep4-GAL4,UASmcherry; UAS-Skap*$^{RNAi}$), **(B7)** *Sdh*$^{RNAi}$ (*Tep4-GAL4,UASmcherry;UAS-Sdh*$^{RNAi}$), **(B8)** *Fum*$^{RNAi}$ (*Tep4-GAL4,UASmcherry;UAS-Fum*$^{RNAi}$*),* and **(B9)** *Mdh*$^{RNAi}$ (*Tep4-*

*GAL4,UASmcherry;UAS-Mdh*$^{RNAi}$), **(B3 & B5)** lymph glands shows reduction in crystal cells (PPO+; magenta), compare to **(B1)** control. **(F)** Quantification is total number of crystal cells (PPO+) cells in entire LG lobe, *Tep4-GAL4,UASmcherry/+* (control, n=51), *Tep4-GAL4,UASmcherry/CS*$^{RNAi}$ (n=38, p=0.0896), *Tep4-GAL4,UASmcherry/mAcon1*$^{RNAi}$ (n=14, p<0.0001), *Tep4-GAL4,UASmcherry/Idh*$^{RNAi}$ (n=43, p=0.1000), *Tep4-GAL4,UASmcherry/α-Kdh*$^{RNAi}$ (n=20, p<0.0001), *Tep4-GAL4,UASmcherry/Skap*$^{RNAi}$ (n=33, p=0.1494), *Tep4-GAL4,UASmcherry/Sdh*$^{RNAi}$ (n=35, p>0.9999), *Tep4-GAL4,UASmcherry/Fum*$^{RNAi}$ (n=30, p=0.9986), *Tep4-GAL4,UASmcherry/Mdh*$^{RNAi}$ (n=48, p>0.9999). **(C1-C9)** Representative images showing crystal cells in the lymph gland of 3$^{rd}$ instar *Drosophila* larva, when TCA cycle was modulated in intermediary progenitor population (IZ) with the help of *ChIZ-GAL4.* **(C1)** *Control* (*ChIZ-GAL4*/+), **(C2)** expressing *CS*$^{RNAi}$ (*ChIZ-GAL4*;UAS-*CS*$^{RNAi}$), **(C3)** *mAcon1*$^{RNAi}$ (*ChIZ-GAL4;UAS-mAcon1*$^{RNAi}$), **(C4)** *α-Kdh*$^{RNAi}$ (*ChIZ-GAL4*;*UAS-α-Kdh*$^{RNAi}$), **(C5)** *Idh*$^{RNAi}$ (*ChIZ-GAL4;UAS-Idh*$^{RNAi}$), **(C6)** Skap$^{RNAi}$ (*ChIZ-GAL4;UAS-Skap*$^{RNAi}$), **(C7)** *Sdh*$^{RNAi}$ (*ChIZ-GAL4;UAS-Sdh*$^{RNAi}$), **(C8)** *Fum*$^{RNAi}$ (*ChIZ-GAL4;UAS-Fum*$^{RNAi}$),and **(C9)** *Mdh*$^{RNAi}$ (*ChIZ-GAL4;UAS-Mdh*$^{RNAi}$), **(C3-C5)** lymph glands shows reduction in number of crystal cells in comparison to **(C1)** control. **(G)** Quantification is total number of crystal cells (PPO+) cells in entire LG lobe, *ChIZ-GAL4*/+ (control, n=44), *ChIZ-GAL4/CS*$^{RNAi}$ (n=27, p=0.8904), *ChIZ-GAL4/mAcon1*$^{RNAi}$ (n=37, p<0.0001), *ChIZ-GAL4/Idh*$^{RNAi}$ (n=31, p<0.0001), *ChIZ-GAL4/α-Kdh*$^{RNAi}$ (n=50, p<0.0001), *ChIZ*-GAL4/Skap$^{RNAi}$ (n=13, p=0.8805), *ChIZ-GAL4/Sdh*$^{RNAi}$ (n=26, p=0.3933), *ChIZ-GAL4/Fum*$^{RNAi}$ (n=44, p=0.9998), *ChIZ-GAL4/MdhRNAi* (n=48, p=0.5853). **(D1-D9)** Representative images showing crystal cells in the lymph gland of 3$^{rd}$ instar *Drosophila* larva, when TCA cycle was modulated in core-progenitor population (MZ) with the help of *Hml*$^{\Delta}$*-GAL4,UAS-2XEGFP.* **(D1)** *Control* (*Hml*$^{\Delta}$*-GAL4,UAS-2XEGFP*/+), **(D2)** *CS*$^{RNAi}$ (*Hml*$^{\Delta}$*-GAL4,UAS-2XEGFP*;UAS-*CS*$^{RNAi}$), **(D3)** *mAcon1*$^{RNAi}$ (*Hml*$^{\Delta}$*-GAL4,UAS-2X*

*EGFP*;*UAS-mAcon1*$^{RNAi}$*),* **(D4)** *α-Kdh*$^{RNAi}$ (*Hml△-GAL4,UAS-2X EGFP*;*UAS-α-Kdh*$^{RNAi}$ ), **(D5)** *Idh*$^{RNAi}$ (*Hml△-GAL4,UAS-2XEGFP*;*UAS-Idh*$^{RNAi}$), **(D6)** *Skap*$^{RNAi}$ (*Hml△-GAL4,UAS-2XEGFP*;*UAS-Skap*$^{RNAi}$), **(D7)** *Sdh*$^{RNAi}$ (*Hml△-GAL4,UAS-2XEGFP*;*UAS-Sdh*$^{RNAi}$), **(D8)** *Fum*$^{RNAi}$ (*Hml△-GAL4,UAS-2XEGFP*;*UAS-Fum*$^{RNAi}$),and **(D9)** *Mdh*$^{RNAi}$ (*Hml△-GAL4,UAS-2XEGFP*;*UAS-Mdh*$^{RNAi}$), **(D2, D3 & D5)** lymph glands shows decrease in crystal cell population in comparison to **(D1)** control. **(H)** Quantification is total number of crystal cells (PPO+) cells in entire LG lobe, *Hml△-GAL4,UAS-2X EGFP*/+ (control, n=48), *Hml△-GAL4,UAS-2X EGFP*/*CS*$^{RNAi}$ (n=37, p<0.0001), *Hml△-GAL4,UAS-2X EGFP* /mAcon1$^{RNAi}$ (n=36, p<0.0001), *Hml△-GAL4,UAS-2X EGFP* /Idh$^{RNAi}$ (n=46, p=0.2199), *Hml△-GAL4,UAS-2X EGFP*/α-Kdh$^{RNAi}$ (n=26, p<0.0001), *Hml△-GAL4,UAS-2X EGFP* /Skap$^{RNAi}$ (n=31, p>0.9999), *Hml△-GAL4,UAS-2X EGFP* /Sdh$^{RNAi}$ (n=61, p=0.8906), *Hml△-GAL4,UAS-2X EGFP* /Fum$^{RNAi}$ (n=44, p=0.0745), *Hml△-GAL4,UAS-2X EGFP* /Mdh$^{RNAi}$ (n=34, p=0.9962).Data is presented as median plots (*p<0.05; **p<0.01; ***p<0.001; ****p<0.0001), ordinary one-way ANOVA. Scale bar: 20μm. 'n'=lymph gland lobes. Comparisons for significance are with respective controls.

**Supplementary figure 2:**

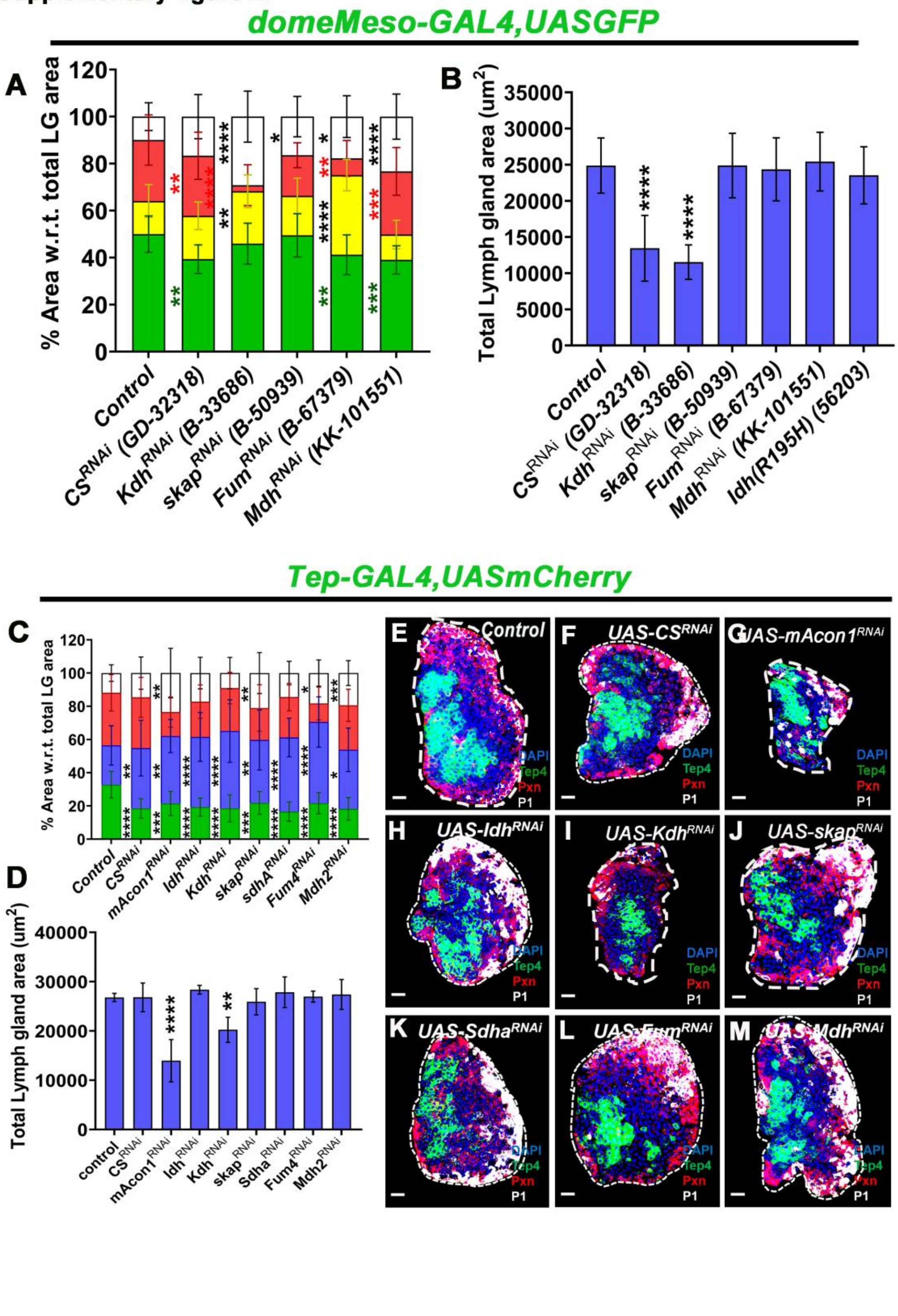

1620

**Supplementary Figure 2:**

**(A-B)** To validate the specificity and efficiency of the observed phenotypes, multiple independent RNAi lines targeting the same gene were used wherever available, to rule out potential off-target effects. **(A)** Quantification is relative % area with respect to total lymph gland area for maintenance and differentiation profile, marked for respective zones (MZ: Dome$^+$, *CZ*, IZ: Dome$^+$Pxn$^+$, *IZ*, CZ: Pxn$^+$, red and P1$^+$, white) from *domeMeso>GFP/+* (control, n=17), *domeMeso>GFP/CS*$^{RNAi}$ (GD-32318; n=12, *CZ*: p=0.0046; *IZ*: p=0.3800; *CZ*: p>0.9999, *P1 area*: p=0.2218), *domeMeso>GFP/Kdh*$^{RNAi}$ (B-33686; n=19, *CZ*: p=0.4984; *IZ*: p=0.0034; *CZ*: p<0.0001, *P1 area*: p<0.0001), *skap*$^{RNAi}$ (B-50939; n=15, *CZ*: p>0.9999; *IZ*: p=0.7693; *CZ*: p=0.0285, *P1 area*: p=0.1996), *domeMeso>GFP/Fum*$^{RNAi}$ (B-67379; n=27, *CZ*: p=0.0044; *IZ*: p<0.0001; *CZ*: p<0.0001, *P1 area*: p=0.0333), *domeMeso>GFP/Mdh*$^{RNAi}$ (KK-101551; n=23, *CZ*: p=0.0004; *IZ*: p=0.4852; *CZ*: p=0.9992, *P1 area*: p<0.0001). **(B)** Graph showing quantification in terms of total lymph gland area, *domeMeso>GFP/+* (*Control*, n=17), *domeMeso>GFP/CS*$^{RNAi}$ (GD-32318; n=14, p<0.0001), *domeMeso>GFP/Kdh*$^{RNAi}$ (B-33686; n=19, p<0.0001), *domeMeso>GFP/skap*$^{RNAi}$ (B-50939; n=14, p>0.9999), *domeMeso>GFP/Fum*$^{RNAi}$ (B-67379; n=27, p=0.9991), *domeMeso>GFP/Mdh*$^{RNAi}$ (KK-101551; n=23, p=0.9989), *domeMeso>GFP/IdhR195H* (B-56203; n=17, p=0.8988). **(C)** graph representing quantification of core-progenitor (Tep4$^+$; green), differentiating (blue and red) population. **(D)** graph representing the size in terms of total area of lymph gland. **(E)** Control (Tep4-GAL4,UASmcherry/+) lymph gland showing general distribution of core-progenitors (Tep4$^+$; green) and differentiating (blue and red) population at 3rd instar larval stage, **(F)** *CS*$^{RNAi}$ (*Tep4-GAL4,UASmcherry/UAS-CS*$^{RNAi}$), **(G)** mAcon1$^{RNAi}$ (*Tep4-GAL4,UASmcherry/UAS-mAcon1*$^{RNAi}$), **(H)** *Idh*$^{RNAi}$ (*domeMeso-Gal4,UAS-GFP/UAS-*

*Idh*$^{RNAi}$), **(I)** *α-Kdh*$^{RNAi}$ (*Tep4-GAL4,UASmcherr/UAS-α-Kdh*$^{RNAi}$), **(J)** *skap*$^{RNAi}$ (*domeMeso-Gal4,UAS-GFP/UAS-skap*$^{RNAi}$), **(K)** *Sdh*$^{RNAi}$ (*Tep4-GAL4,UASmcherr/UAS-Sdh*$^{RNAi}$), **(L)** *Fum*$^{RNAi}$ (*Tep4-GAL4,UASmcherry/UAS-Fum*$^{RNAi}$), and **(M)** *Mdh*$^{RNAi}$ (*Tep4-GAL4,UASmcherry/UAS-Mdh*$^{RNAi}$), **(F-M)** lymph glands showing reduction in core-progenitor (green) cells and concomitant increase in differentiating population (Te4-Pxn-; blue), compare to **(E)** control. **(G, I)** both lymph glands also show smaller size as compared to **(C)** control. **(C)** Quantification is relative % area with respect to total lymph gland area for maintenance and differentiation profile, *Tep4-GAL4,UASmcherry/+* (control, n=35), *Tep4-GAL4,UASmcherry/CS*$^{RNAi}$ (n=31, green: p<0.0001; blue: p=0.0076; red: p>0.9999; white: p=0.9345), *Tep4-GAL4,UASmcherry/mAcon1*$^{RNAi}$ (n=17, green: p=0.0007; blue: p=0.0027; red: p<0.0001; white: p=0.0084), *Tep4-GAL4,UASmcherry/Idh*$^{RNAi}$ (n=26, green: p<0.0001; blue: p<0.0001; red: p=0.0069; white: p=0.1003), *Tep4-GAL4,UASmcherry/α-Kdh*$^{RNAi}$ (n=15, green: p<0.0001; blue: p<0.0001; red: p>0.9999; white: p>0.9999), *Tep4-GAL4,UASmcherry/Skap*$^{RNAi}$ (n=22, green: p=0.0008; blue: p=0.0079; red: p=0.0077; white: p=0.0088), *Tep4-GAL4,UASmcherry/Sdh*$^{RNAi}$ (n=17, green: p<0.0001; blue: p<0.0001; red: p=0.3595; white: p>0.9999), *Tep4-GAL4,UASmcherry/Fum*$^{RNAi}$ (n=25, green: p<0.0001; blue: p<0.0001; red: p<0.0001; white: p=0.0134), *Tep4-GAL4,UASmcherry/Mdh*$^{RNAi}$ (n=28, green: p<0.0001; blue: p=0.0248; red: p=0.8038; white: p=0.0008), **(D)** Quantification in terms of total lymph gland area, *Tep4-GAL4,UASmcherry/+* (control, n=78), *Tep4-GAL4,UASmcherry/CS*$^{RNAi}$ (n=83, p>0.9999), *Tep4-GAL4,UASmcherry/mAcon1*$^{RNAi}$ (n=43, p<0.0001), *Tep4-GAL4,UASmcherry/Idh*$^{RNAi}$ (n=51, p=0.8798), *Tep4-GAL4,UASmcherry/α-Kdh*$^{RNAi}$ (n=34, p=0.0016), *Tep4-GAL4,UASmcherry/Skap*$^{RNAi}$ (n=28, p=0.9971), *Tep4-GAL4,UASmcherry/Sdh*$^{RNAi}$ (n=48, p=0.9845), *Tep4-GAL4,UASmcherry/Fum*$^{RNAi}$ (n=55, p>0.9999), *Tep4-GAL4,UASmcherry/Mdh*$^{RNAi}$ (n=80, p=0.9993). Data is presented as bar graph (*$p<0.05$; **$p<0.01$; ***$p<0.001$; ****$p<0.0001$), ordinary one-way ANOVA. Scale bar: 20µm. ‘n’=lymph gland lobes. DAPI marks DNA, Core Progenitors (Tep4$^+$; Green), Double negative (Tep4$^-$ Pxn$^-$; Blue), Differentiating population (Pxn$^+$; Red), terminally differentiated population (P1$^+$; White). Comparisons for significance are with control values. LG lobes are outlined with white border along with removal of other accessory tissues for clear demarcation.

### Supplementary Figure: 3

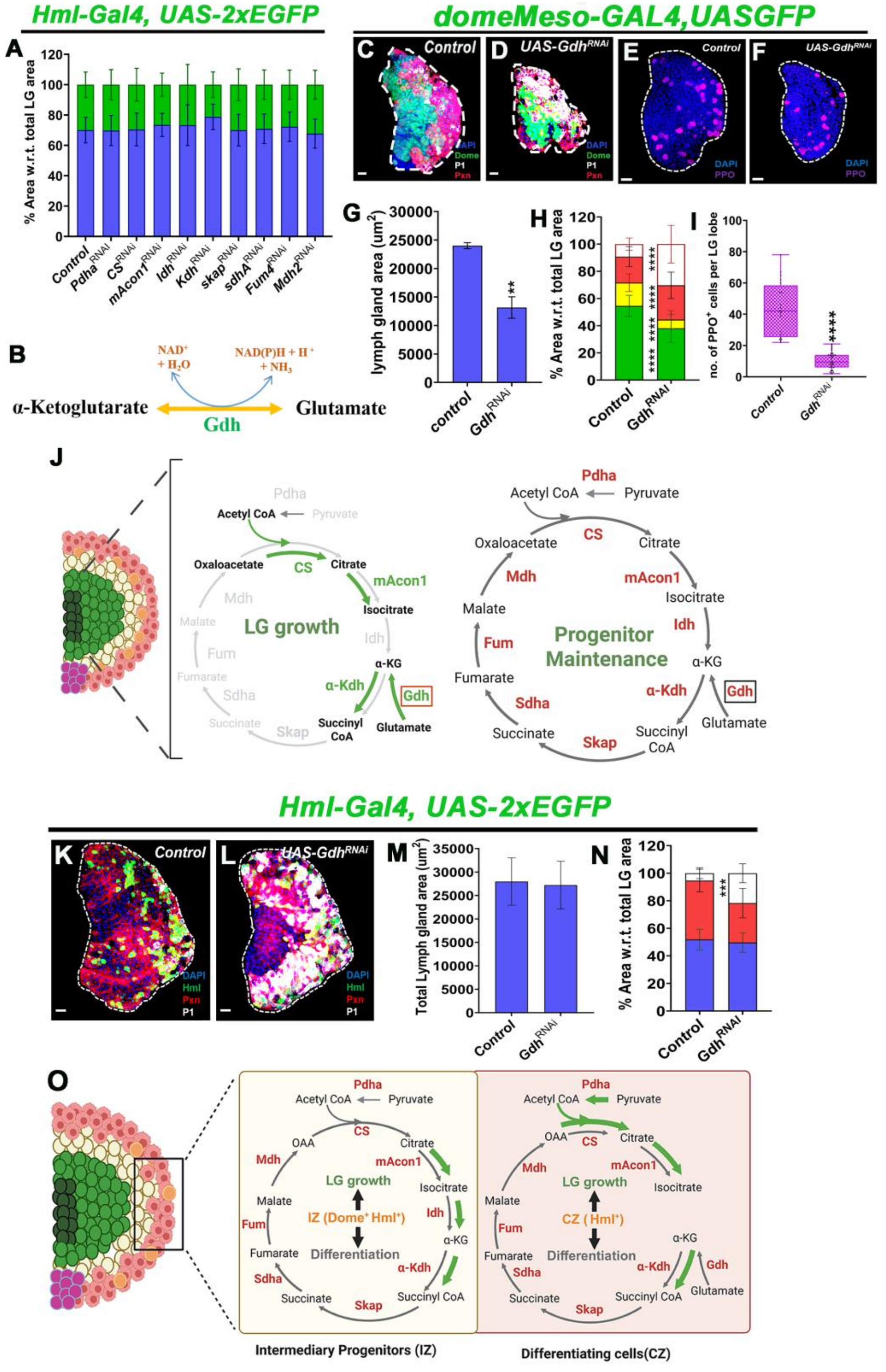

**Supplementary Figure 3:**

**(A)** Graph showing quantification is relative % area with respect to total lymph gland area for maintenance (non-Hml; blue) and differentiation profile (Hml; green), *Hml△-GAL4,UAS-2XEGFP/+* (*Control*, n=60), *Hml△-GAL4,UAS-2XEGFP/Pdha*$^{RNAi}$ (n=25, blue: p>0.9999; green: p>0.9999), *Hml△-GAL4,UAS-2XEGFP/CS*$^{RNAi}$ (n=60, blue: p>0.9999; green: p>0.9999), *Hml△-GAL4,UAS-2XEGFP/mAcon1*$^{RNAi}$ (n=48, blue: p=0.4358; green: p=0.4358), *Hml△-GAL4,UAS-2XEGFP /Idh*$^{RNAi}$ (n=47, blue: p=0.3033; green: p=0.3033), *Hml△-GAL4,UAS-2XEGFP/α-Kdh*$^{RNAi}$ (n=34, blue: p=0.0003; green: p=0.0003), *Hml△-GAL4,UAS-2XEGFP/skap*$^{RNAi}$ (n=38, blue: p>0.9999; green: p>0.9999), *Hml△-GAL4,UAS-2XEGFP/Sdh*$^{RNAi}$ (n=52, blue: p>0.9999; green: p>0.9999), *Hml△-GAL4,UAS-2X EGFP/Fum*$^{RNAi}$ (n=45, blue: p=0.9171; green: p=0.9171), *Hml△-GAL4,UAS-2X EGFP/Mdh*$^{RNAi}$ (n=47, blue: p>0.9999; green: p>0.9999). **(B)** Schematic for chemical equation showing *Glutamate dehydrogenase (Gdh)* function in conversion of α-KG to glutamate in the presence of the cofactors involved. **(C-F)** images representing differentiation profile and growth defect phenotype of the *Gdh* loss of function in progenitors. These results ascertain our hypothesis of *Gdh* as a source of α-KG for the functioning of *α-Kdh*, when cycle is modular as it phenocopied all the effects of *α-Kdh* loss of function. **(C)** *Control (domeMeso-Gal4,UAS-GFP/+)* lymph gland showing general distribution of progenitor *(Dome+;* green*),* differentiating *(Dome*$^{+}$*Pxn*$^{+}$*;* yellow *and Pxn*$^{+}$ *;* red*)* and terminally differentiated (*P1*$^{+}$; white) population at 3rd instar larval stage, **(D)** *Gdh*$^{RNAi}$ *(domeMeso-Gal4,UAS-GFP;UAS-Gdh*$^{RNAi}$*),* it shows reduction in progenitor population (*Dome*$^{+}$; green), due to expansion of Pxn$^{+}$ population (red) and mature plasmatocytes (P1$^{+}$; white) in comparison to **(C)** *Control*. **(E)** *Control (domeMeso-Gal4,UAS-GFP/+)* lymph gland showing general distribution of crystal cells *(PPO*$^{+}$*;*

magenta) population at 3rd instar larval stage, **(F)** *$Gdh^{RNAi}$ (domeMeso-Gal4,UAS-GFP;UAS-$Gdh^{RNAi}$),* it shows reduction in crystal cell population (*PPO+;* magenta) in comparison to **(E)** *Control*. **(G)** Graph representing the quantification of total LG area, *domeMeso>GFP/+* (*Control*, n=35)*, domeMeso>GFP/$Gdh^{RNAi}$* (n=48, p=0.0079)*,* **(H)** graph representing the quantification of relative % area with respect to total lymph gland area for ($Dome^+$; MZ), differentiating ($Dome^+Pxn^+$; IZ and only *Pxn+;* CZ*)* and and terminally differentiated (*$P1^+$*; white) population at 3rd instar larval stage, *domeMeso>GFP/+* (control, n=37), *domeMeso>GFP/$Gdh^{RNAi}$* (n=35, MZ: $p<0.0001$; IZ: $p<0.0001$; CZ: $p<0.0001$; P1 area: $p<0.0001$), **(I)** Graph representing the quantification of total crystal cells in LG lobe, *domeMeso>GFP/+ (control, n=13), domeMeso>GFP/$Gdh^{RNAi}$ (n=13, p=0.0051).* **(J)** During LG (lymph gland) growth, a subset of TCA cycle enzymes (shown in green) is upregulated, promoting citrate production and α-ketoglutarate (α-KG) flux. Enzymes highlighted include *CS, mAcon1, α-Kdh,* and *Gdh*, supporting anabolic metabolism required for tissue expansion. In contrast, during progenitor maintenance, a broader set of TCA cycle enzymes (shown in red) is expressed, maintaining metabolic balance and sustaining the progenitor pool. Differential enzyme usage suggests distinct metabolic states underpin growth versus homeostasis in the LG progenitor niche. **(K-L)** images representing differentiation profile of the *Gdh* loss of function in CZ. **(K)** *Control (Hml△-GAL4,UAS-2XEGFP/+)* lymph gland showing general distribution of differentiating population *(Hml+;* green, Pxn+; red*),* progenitors (blue) and terminally differentiated (*$P1^+$*; white) population at 3rd instar larval stage, **(L)** *$Gdh^{RNAi}$ (Hml△-GAL4,UAS-2XEGFP/UAS-$Gdh^{RNAi}$),* it does not show reduction in progenitor (blue) or differentiating population ($Pxn^+$; red), but we do observe expansion of mature plasmatocytes ($P1^+$; white) in comparison to **(K)** *Control*. Also no growth defect of the LG was observed here. **(M)** Graph representing the quantification of total LG area, *Hml△-GAL4,UAS-2XEGFP/+ (control, n=13), Hml△-*

*GAL4,UAS-2XEGFP/Gdh*$^{RNAi}$ *(n=13, p=0.0051)* **(N)** Graph representing the quantification of relative % area with respect to total lymph gland area for progenitors (DAPI; blue), differentiating (Pxn$^{+}$; red*)* and terminally differentiated (*P1*$^{+}$; white) population at 3rd instar larval stage. *Hml△-GAL4,UAS-2XEGFP/+* (*contro*l, n=13), *Hml△-GAL4,UAS-2X EGFP/Gdh*$^{RNAi}$ (n=7, blue: p=0.5356; red: p=0.5356; white: p=0.0001). **(O)** Model showing spatial compartmentalization and functional contribution of TCA cycle enzymes during *Drosophila* larval LG development. Enzymes acting in the intermediary zone (IZ; Dome$^{+}$Hml$^{+}$), including *CS, mAcon1, α-Kdh,* and *Gdh*, promote progenitor proliferation and maintain LG growth. Loss of these enzymes leads to either autonomous or non-autonomous regulation of progenitor maintenance and differentiation. *Pdha, CS, mAcon1,* and *α-Kdh* are also active in the cortical zone (CZ; *Hml* $^{+}$), supporting continued LG growth through metabolic activity linked to α-KG flux. *Sdha, Mdh,* and *Fum* contribute to later developmental stages and maintain LG homeostasis. Arrows indicate metabolite flow through the TCA cycle, with green arrows representing active reactions contributing to growth and later maintenance, while black arrow represent steps for regulation of differentiation. Data is presented as median plots (*p<0.05; **p<0.01; ***p<0.001; ****p<0.0001), ordinary one-way ANOVA & student unpaired t-test. Scale bar: 20µm. 'n'=lymph gland lobes. DAPI marks DNA, Progenitors (Dome+; Green for Dome> and blue for Hml>), Differentiating population (Dome$^{+}$Pxn$^{+}$; yellow and *Pxn*$^{+}$*;* red for dome> and Hml$^{+}$; green and *Pxn*$^{+}$*;* red for Hml>) and terminally differentiated (*P1*$^{+}$; white) population. Comparisons for significance are with control values. LG lobes are outlined with white border along with removal of other accessory tissues for clear demarcation.

1750

**Supplementary Figure 4:**

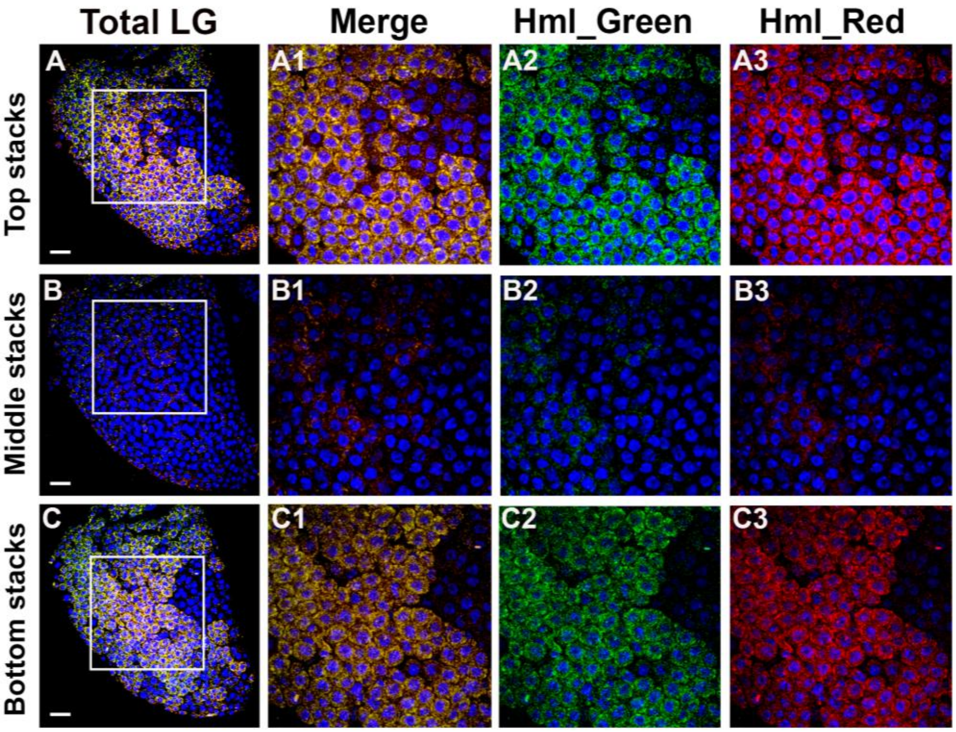

**Supplementary Figure 4:**

To validate the SABER-FISH protocol in the *Drosophila* larval LG, probes were designed according to the established protocol. We first tested probes targeting the Hml transcript, labeled with two spectrally distinct fluorophores (488 nm and 546 nm). The resulting signal showed ~95% overlap between the two fluorophores, confirming the specificity of the probes and the absence of off-target hybridization. **(A–A3)** Top optical stack showing the full LG with merged and individual channels for Hml transcripts. **(B–B3)** Middle stack showing merged and separate channels. **(C–C3)** Bottom stack showing merged and separate channels. For each region of interest (ROI), a 250 × 250 pixel area was cropped to generate zoomed-in views of the respective LG stacks. Four optical sections (0.5 µm each) were merged to generate representative top, middle, and bottom stacks. A higher concentration of Hml transcripts was detected in the top and bottom stacks relative to the middle stack, highlighting the spatial distribution pattern of Hml expression. This distribution closely mirrors that observed using the HmlΔ-GAL4, UAS-2×EGFP reporter construct (see Fig. 2). Images were acquired using a 60X oil immersion objective with a step size of 0.5 µm. Scale bar: 20 µm. DAPI marks nuclei (blue); gene transcripts, red and green channels as indicated.

**Supplementary Figure 5:**

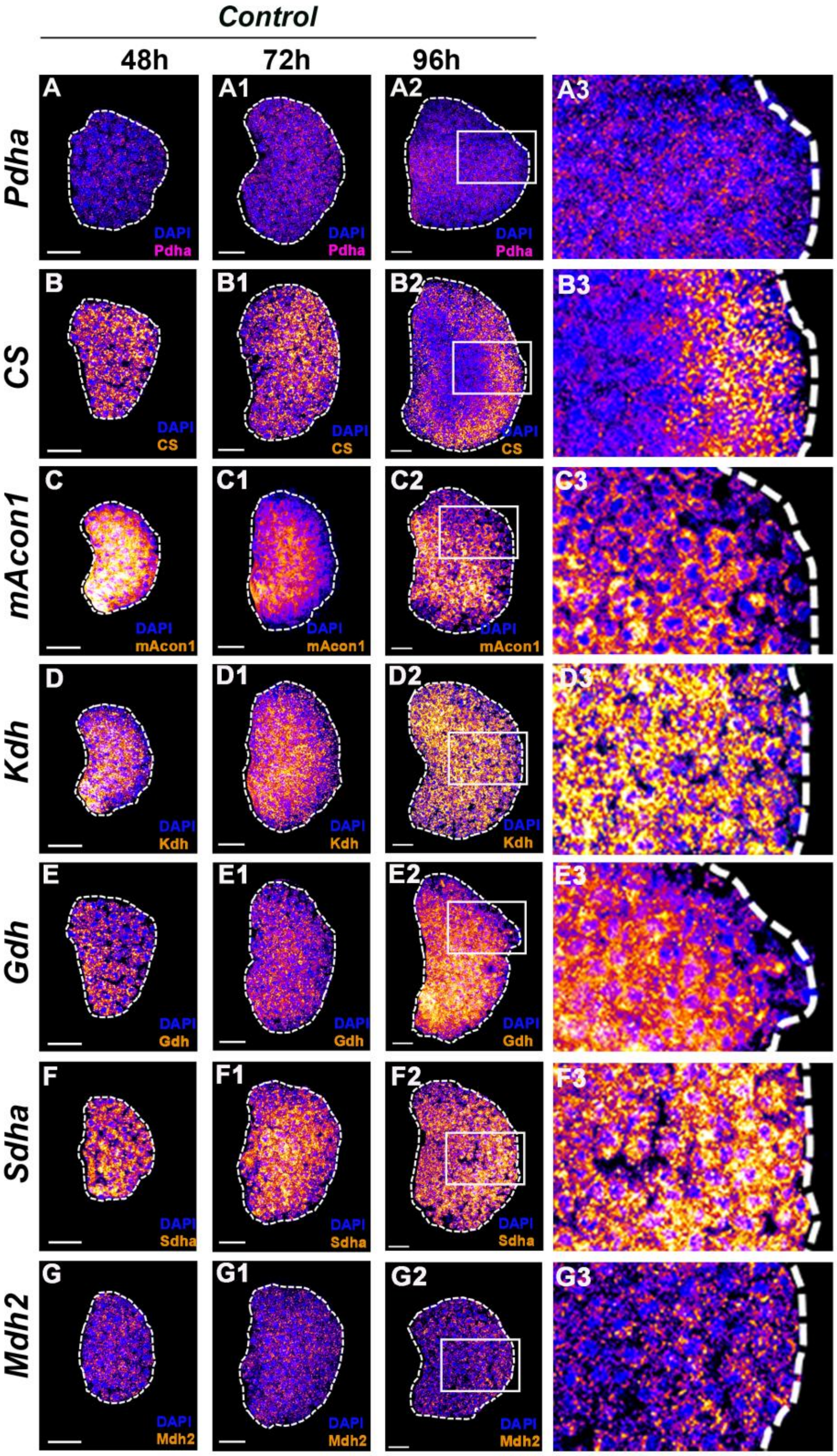

1770

**Supplementary Figure 5:**

SABER-FISH was performed in a control background ($Y^{+}W^{+} \times w^{1118}$) to examine the temporal and spatial distribution of transcripts encoding TCA cycle enzymes during LG development. For each region of interest (ROI), a 185 × 125 pixel area was cropped to generate zoomed-in views of the respective LG stacks. **(A–A3)** *Pdh* transcripts were uniformly distributed across all LG cells at early time points (48 h and 72 h AEL) but became enriched in the inner core (medullary zone, MZ) by 96 h. **(B–B3)** *CS* mRNA was initially enriched in the MZ from 48–72 h but showed a marked shift toward the cortical zone (CZ) at 96 h. **(C–C3)** mAcon1 expression was consistently higher in the core region across all time points, with peak levels observed at 48 h. **(D–D3)** *Kdh* transcripts were predominantly enriched in the core region throughout all developmental stages examined. **(E–E3)** *Gdh* expression remained restricted to the inner core region across all time points. **(F–F3)** Sdh showed progressive upregulation over time, being initially enriched in the core region and later (96 h) also observed in outer CZ cells. **(G–G3)** *Mdh* exhibited weak and uniform expression across all zones and developmental stages. Scale bar: 20 µm. DAPI marks DNA (blue); gene transcripts are shown in Fire LUT. LG lobes are outlined with white border along with removal of other accessory tissues for clear demarcation.

### Supplementary Figure 6

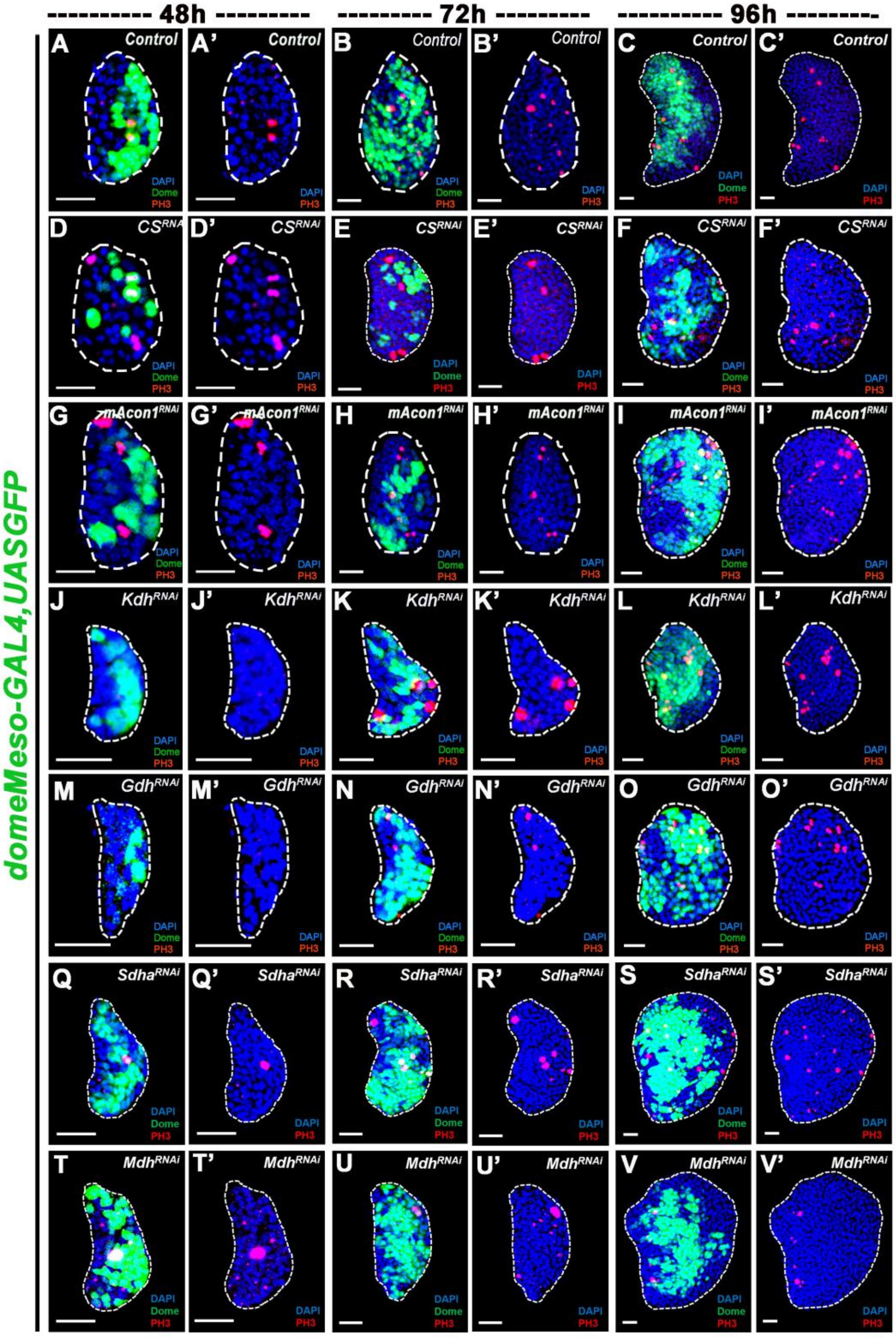

1789

**Supplementary Figure 6:**

Representative images for proliferative potential (pH3) as well as progenitor (Dome) vs non-progenitor (non-dome) blood cell population's contribution during the development of the LG (48h, 72h and 96h) in *Drosophila* 3$^{rd}$ instar larval stage. In control lymph glands (LGs), the normal trajectory of growth shows that total cell number approximately doubles between 48–72h, driven primarily by proliferating progenitors (Dome$^{+}$, green), with minimal contribution from non-progenitors (Dome$^{-}$, blue), corresponding to a mitotic index (MI) of ~4–5%. Between 72–96h, cell number triples, with contributions from both progenitor (MI ~3%) and non-progenitor (MI ~5%) populations. Growth continues between 96–120h, with total cell number doubling again, predominantly driven by non-progenitor proliferation (MI ~5%). Knockdown of CS and mAcon1 results in a growth defect evident from 72h; however, at 48h, enhanced mitotic activity in the Dome$^{-}$ population compensates for the loss of progenitors, preventing an immediate size defect. By 72h, this compensatory effect is insufficient to offset the reduced progenitor proliferation, leading to a noticeable growth defect. In contrast, loss of *Kdh* and *Gdh* causes an earlier growth reduction, apparent at 48h, primarily due to impaired mitotic activity in the progenitor (Dome$^{+}$) compartment. *Sdha* and *Mdh* loss-of-function serve as additional controls and do not display any detectable size or growth defects. **(A)** Represents image with dome and **(A')** image without dome. **(A-C')** *Control (domeMeso-Gal4,UAS-GFP/+)*, **(D-F')** *CS$^{RNAi}$ (domeMeso-Gal4,UAS-GFP;UAS-CS$^{RNAi}$)*, **(G-I')** *mAcon1$^{RNAi}$ (domeMeso-Gal4,UAS-GFP;UAS-mAcon1$^{RNAi}$)*, **(J-L')** *α-Kdh$^{RNAi}$ (domeMeso-Gal4,UAS-GFP;UAS-α-Kdh$^{RNAi}$ )*, **(M-O')** *GdhRNAi (domeMeso-Gal4,UAS-GFP;UAS-Gdh$^{RNAi}$*), **(Q-S')** *Sdh$^{RNAi}$ (domeMeso-Gal4,UAS-GFP;UAS-Sdh$^{RNAi}$)*, **(T-V')** *Mdh$^{RNAi}$ (domeMeso-Gal4,UAS-GFP;UAS-Mdh$^{RNAi}$)*. Scale bar: 20µm. DAPI marks DNA, Progenitors (Dome+; Green), non-progenitor (Dome-

; blue) population, and mitotic index (pH3+; Red). LG lobes are outlined with white border along with removal of other accessory tissues for clear demarcation.

### Supplementary Figure 7

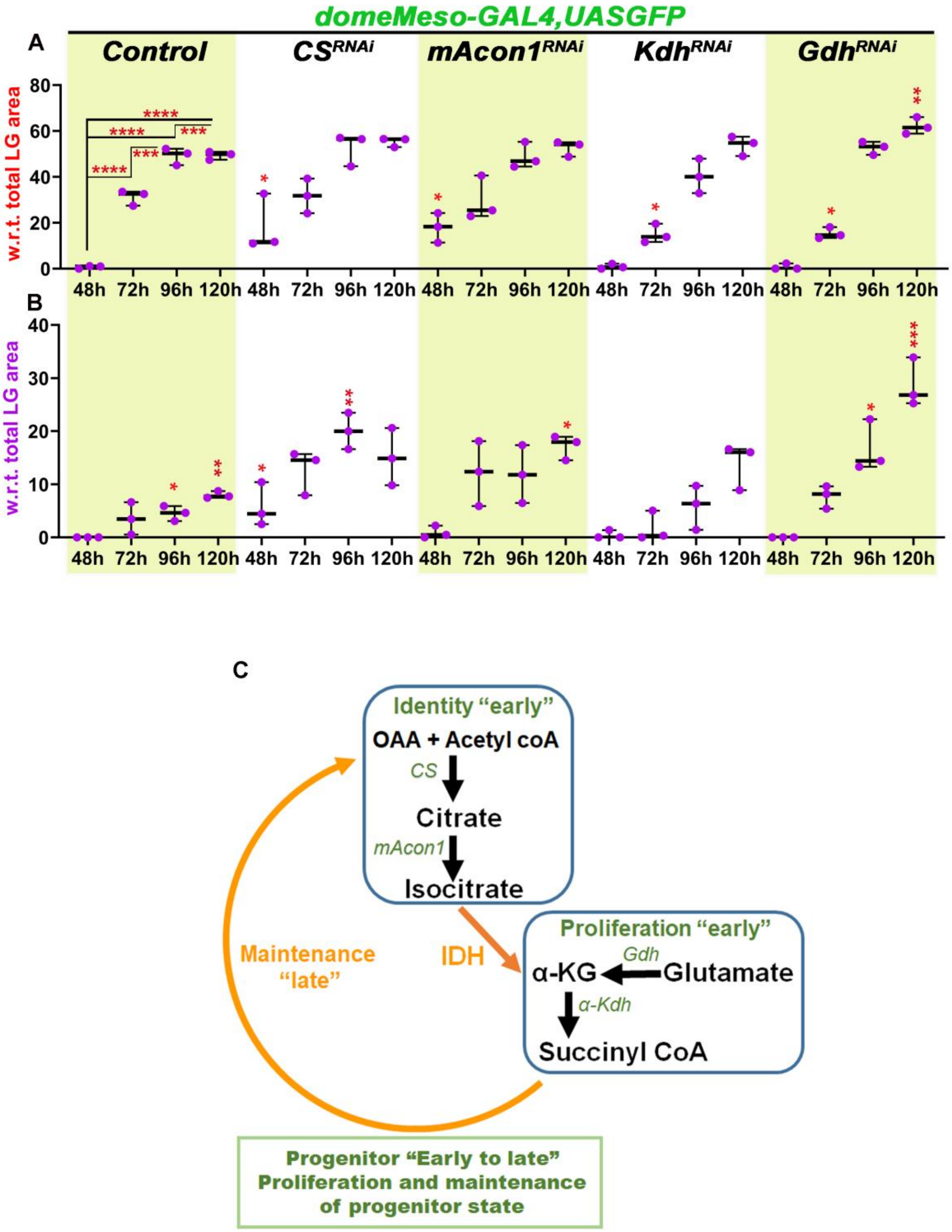

1817

**Supplementary Figure 7:**

**Temporal differentiation profile of the wandering 3$^{rd}$ instar *Drosophila* larval LG.** In control lymph glands (LGs), differentiation markers Pxn and P1 are not detected until ~60 h, with no signal observed at 48 h. By 72h, approximately the Pxn and P1 could be detected in the LG, which further increases through the development. Knockdown of *CS* and *mAcon1* leads to the premature appearance of Pxn$^+$ cells at 48h, along with early emergence of P1$^+$ cells in *CS* knockdown LGs, highlighting the essential roles of *CS* and *mAcon1* in maintaining progenitor identity, their loss resulting in precocious differentiation. In contrast, *Kdh* and *Gdh* knockdowns do not exhibit early differentiation, and the normal trajectory of Pxn/P1 expression is maintained, further supporting their primary role in promoting LG growth during early development rather than in progenitor maintenance. **(A)** Graph showing the differentiation profile (Pxn$^+$), based on relative % area with respect to total lymph gland (LG) area over developmental time, *domeMeso>GFP/+* (*control*, 48h, n=31; 72h, n=34, p<0.0001; 96h, n=24, p<0.0001; 120h, n=21, p<0.0001), *domeMeso>GFP/CS*$^{RNAi}$ (48h, n=20, p=0.0195; 72h, n=23, p=0.9999; 96h, n=18, p=0.8549; 120h, n=18, p=0.1338), *domeMeso>GFP/mAcon1*$^{RNAi}$ (48h, n=21, p=0.0228; 72h, n=30, p=0.9933; 96h, n=29, p>0.9999; 120h, n=25, p=0.5532), *domeMeso>GFP/α-Kdh*$^{RNAi}$ (48h, n=25, p>0.9999; 72h, n=39, p=0.0263; 96h, n=27, p=0.2277; 120h, n=32, p=0.3222), and *domeMeso>GFP/Gdh*$^{RNAi}$ (48h, n=20, p>0.9999; 72h, n=18, p=0.0294; 96h, n=19, p=0.8574; 120h, n=14, p=0.0021), (**B)** Graph showing the terminal differentiation profile (P1$^+$), based on relative % area with respect to total lymph gland (LG) area over developmental time. *domeMeso>GFP/+* (*control*, 48h, n=31; 72h, n=34, p=0.0840; 96h, n=24, p=0.0294; 120h, n=21, p=0.0012), *domeMeso>GFP/CS*$^{RNAi}$ (48h, n=20, p=0.0146; 72h, n=23, p=0.0517; 96h, n=18, p=0.0033; 120h, n=18, p=0.1380), *domeMeso>GFP/mAcon1*$^{RNAi}$ (48h, n=21, p=0.9413; 72h, n=30, p=0.0694; 96h, n=29, p=0.1547; 120h, n=25, p=0.0488), *domeMeso>GFP/α-Kdh*$^{RNAi}$ (48h, n=25, p=0.9948; 72h, n=39, p=0.9471; 96h, n=27, p=0.9844; 120h, n=32, p=0.2514), and *domeMeso>GFP/Gdh*$^{RNAi}$ (48h, n=20, p>0.9999; 72h, n=18, p=0.5233; 96h, n=19, p=0.0156; 120h, n=14, p=0.0002). **(C)** This schematic illustrates two early metabolic modules governing progenitor cell behavior. The "Identity early" module is characterized by *CS* and *mAcon1* enzymes, which controls the stemness of the progenitors. The "Proliferation early" module includes genes such as *Gdh* and *α-Kdh*, which support early proliferative activity. *Idh* serves as a key metabolic node connecting the identity and proliferation programs (orange arrow). A broader "Maintenance late" module (large orange arc) reinforces the transition from early to late progenitor states, supporting long-term progenitor maintenance. Together, these modules integrate metabolic signals to regulate progenitor progression from early identity through proliferation and into a maintained progenitor state. Data is presented as median plots (*p<0.05; **p<0.01; ***p<0.001; ****p<0.0001), ordinary one-way ANOVA. Comparisons for significance are with control values.

**Supplementary Figure 8:**

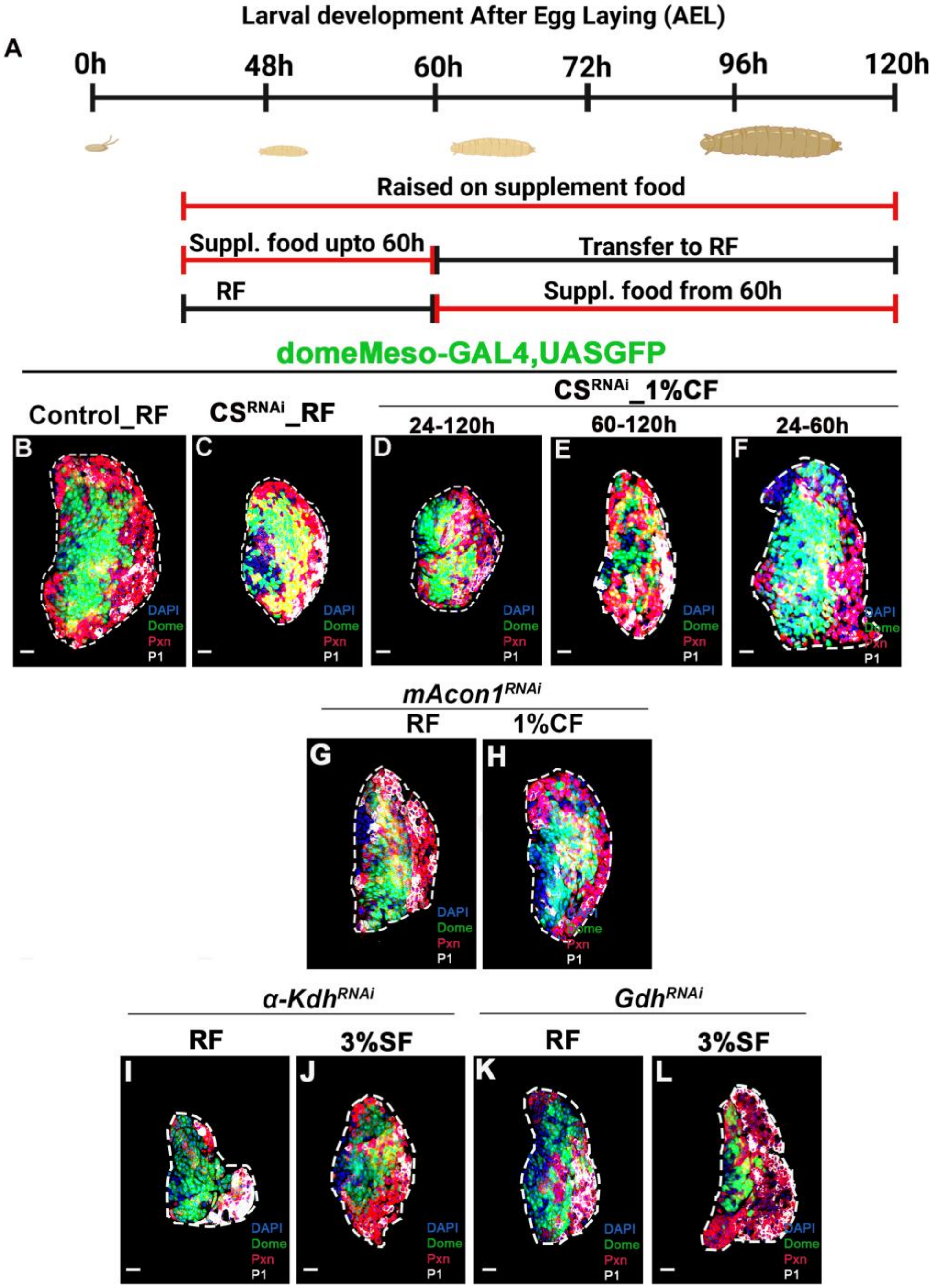

1859

**Supplementary Figure 8:**

**(A)** Developmental timeline of *Drosophila* melanogaster and experimental stages used in the study. The schematic represents the major developmental stages of *Drosophila* melanogaster from embryogenesis to pupal stage under standard culture conditions (25 °C). The black timeline indicates total larval developmental duration (~5-6 days). The red bars indicate specific developmental windows analyzed in this study. Line 1 (top red bar): the larvae were cultured on supplement food (1%CF or 3%SF) for entirety of larval stages. Line 2 (middle red bar): 1st to 2nd instar larvae were reared on supplement food and then transferred to regular food. Line 3 (bottom red bar): after mid-2nd instar larval stage larvae were reared on supplementary food. This 60h time point in the development of larvae was chosen as it is well established differentiation initiation time point. **(B-L)** Representative images showing the effect of different supplemented foods 1% citrate food (CF) and 3% succinate food (SF) on the growth and differentiation of the LG in the loss of function TCA enzymes at wandering 3$^{rd}$ instar larval stage (120h AEL) in the background of *domeMESO-GAL4,UAS-GFP*. **(B)** *Control (domeMeso-Gal4,UAS-GFP/+)* on RF showing progenitors (Dome$^+$; green: MZ), and differentiating population, which is further sub-divided into Dome$^+$Pxn$^+$ (yellow: IZ), Pxn$^+$ (red: CZ) area, including ~10% P1$^+$ (white). **(C)** *$CS^{RNAi}$ (domeMeso-Gal4,UAS-GFP/$CS^{RNAi}$)* on RF has reduced progenitors area and increased differentiation as well as LG size reduction, in comparison to **(B)** control on RF. **(D)** *$CS^{RNAi}$ (domeMeso-Gal4,UAS-GFP/$CS^{RNAi}$)* on 1%CF (from 24-120h) shown no recovery in differentiation and growth in comparison to **(C)** *$CS^{RNAi}$* on RF. **(E)** *$CS^{RNAi}$ (domeMeso-Gal4,UAS-GFP/$CS^{RNAi}$)* on 1%CF (early; from 24-60h) recovered progenitors homeostasis as well as LG size to the extent of control on RF in comparison to **(C)** *$CS^{RNAi}$* on RF. **(F)** *$CS^{RNAi}$ (domeMeso-Gal4,UAS-GFP/$CS^{RNAi}$)* on 1%CF (late; from 60-120h) shown no recovery in differentiation and growth in comparison to **(C)** *CS*$^{RNAi}$ on RF. **(G)** *mAcon1*$^{RNAi}$ *(domeMeso-Gal4,UAS-GFP/mAcon1*$^{RNAi}$*)* on RF has reduced progenitors area and increased differentiation as well as LG size reduction, in comparison to **(B)** *Control* on RF. **(H)** *mAcon1*$^{RNAi}$ *(domeMeso-Gal4,UAS-GFP/mAcon1*$^{RNAi}$*)* on 1%CF (early; from 24-60h) shown no recovery in differentiation and growth in comparison to **(G)** *mAcon1*$^{RNAi}$ on RF, indicating the specificity of citrate's involvement in rescue of differentiation and growth of LG. **(I-L)** Succinate is downstream to *α-Kdh* enzymatic step, therefore we decided to feed succinate to understand if it can rescue the phenotype. **(I)** *Kdh*$^{RNAi}$ *(domeMeso-Gal4,UAS-GFP/α-Kdh*$^{RNAi}$*)* on RF has reduced progenitors area and increased differentiation as well as LG size reduction, in comparison to **(B)** control on RF. **(J)** *α-Kdh*$^{RNAi}$ *(domeMeso-Gal4,UAS-GFP/α-Kdh*$^{RNAi}$*)* on 3%SF (early; from 24-60h) shown no recovery in differentiation and growth in comparison to **(I)** *α-Kdh*$^{RNAi}$ on RF, rather further increase in differentiation was observed, suggesting succinate as pro-differentiation cue. **(K)** *Gdh*$^{RNAi}$ *(domeMeso-Gal4,UAS-GFP/GdhRNAi)* on RF has reduced progenitors area and increased differentiation as well as LG size reduction, in comparison to **(B)** control on RF. **(L)** *Gdh*$^{RNAi}$ *(domeMeso-Gal4,UAS-GFP/Gdh*$^{RNAi}$*)* on 3%SF (early; from 24-60h) shown no recovery in differentiation and growth in comparison to **(K)** *Gdh*$^{RNAi}$ on RF, rather, further increase in differentiation was observed, suggesting succinate as pro-differentiation cue. DAPI marks DNA, Progenitors (Dome+; Green), intermediary progenitors (Dome+ Pxn+; Yellow), Differentiating population (Pxn+; Red), terminally differentiated population (P1+; white). LG lobes are outlined with white border along with removal of other accessory tissues for clear demarcation. Scale bar: 20µm.

### Supplementary Figure 9:

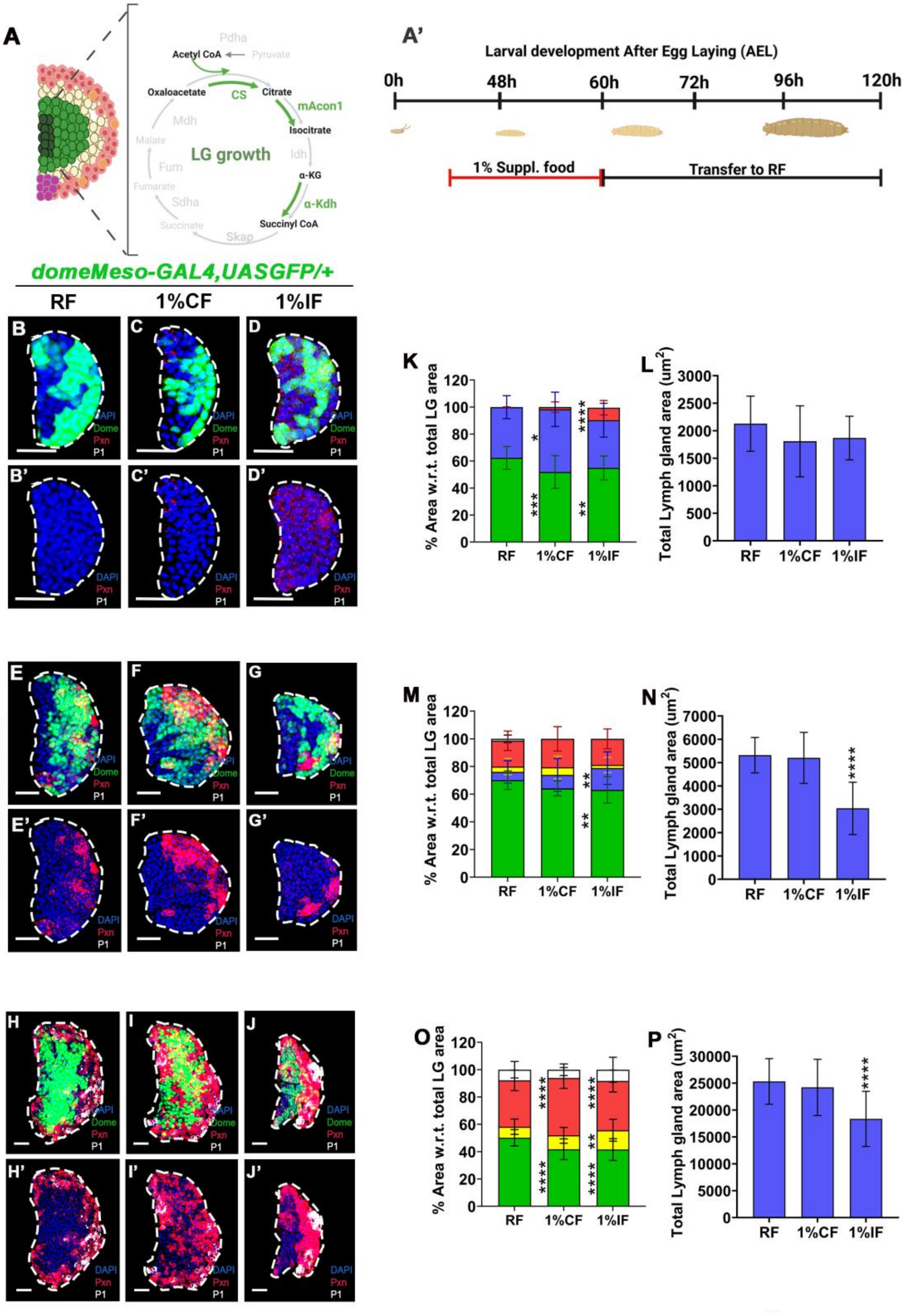

**Supplementary Figure 9:**

**(A)** Developmental timeline of *Drosophila* larvae and experimental stages used in the study. The schematic represents the major developmental stages of *Drosophila* melanogaster from embryogenesis to pupal stage under standard culture conditions (25 °C). The timeline of experiment is as follow: red line represents 1st to 2nd instar larvae were reared on supplement food, i.e., 1% citrate food (CF) or 1% isocitrate food (IF) and then transferred to regular food. **(B-J')** Representative images showing the effect of different supplemented foods 1% CF or 1% IF on the growth and differentiation of the wild-type (*Control*) larval lymph gland (LG) at different developmental stages (48 h, 72 h, and 120 h AEL) in the background of *domeMESO-GAL4,UAS-GFP*. **(B)** represent image with Dome and **(B')** represent image without dome. **(B-B')** Control LGs on regular food (RF) at 48h showing progenitor area (Dome$^{+}$; green) and non-progenitor (Dome$^{-}$; blue) area, with no detectable Pxn staining. **(C-C')** On 1% CF at 48 h, Dome$^{+}$ area was reduced, with a compensatory expansion of Dome$^{-}$ (blue) area and few Pxn$^{+}$ cells; no growth defect is observed in comparison to **(B-B')** *Control* on RF. **(D-D')** On 1% IF at 48 h, reduction in Dome$^{+}$ proportion was compensated by Pxn$^{+}$ cells expansion with non-progenitor (Dome$^{-}$; blue) population remained unchanged; no growth defect was observed in comparison to **(B-B')** *Control* on RF*.* **(E-G')** Images representing the effect on differentiation profile and growth of wild type (*Control*) larval LG on different supplement foods (1%CF or 1%IF) in comparison to regular food (RF) at 72h in development in the background of *domeMESO-GAL4,UAS-GFP*. **(E-E')** Control on RF shows progenitors (Dome$^{+}$; green), non-progenitors (Dome$^{-}$; blue), intermediary progenitors (Dome$^{+}$Pxn$^{+}$), and differentiating population (Pxn$^{+}$), including terminally differentiated (P1$^{+}$) cells. **(F-F')** No major change in differentiation or LG size is observed on 1% CF at 72h. **(G-G')** On 1% IF, progenitors

(Dome+; green) are reduced with a corresponding increase in non-progenitor (Dome$^{-}$ ;blue)population, while other populations remain unchanged; LG growth defect is evident compared to **(E-E')** *Control* on RF at 72h. **(H-J')** images representing the effect on differentiation profile and growth of wild type (control) larval LG on different supplement foods (1%CF or 1%IF) in comparison to regular food (RF) at 120h in development in the background of *domeMESO-GAL4,UAS-GFP*. **(H-H')** *Control* on RF showing progenitors (Dome$^{+}$; green: MZ), intermediary progenitors (Dome$^{+}$Pxn$^{+}$; yellow: IZ), and differentiating population (Pxn$^{+}$; red: CZ) area, including P1$^{+}$ (white). **(I-I')** On 1% CF, Dome$^{+}$ progenitors are reduced with a complementary increase in differentiating Pxn$^{+}$ population, without change in terminal differentiation or LG size. in comparison to **(H-H')** *Control* at 120h on RF. **(J-J')** On 1% IF, Dome$^{+}$ progenitors are reduced due to expansion of the Dome$^{+}$Pxn$^{+}$ fraction; no change in terminal differentiation, but significant growth defect compared to **(H-H')** control on RF at 120h. **(K)** Graph showing quantification of differentiation profile based on relative % area with respect to total lymph gland (LG) area at 48h, *domeMeso>GFP/+* (control, 48h, RF, n=26), *domeMeso>GFP/+* (48h, 1%CF, n=22; green, p=0.0008; blue, p=0.0158; red, p=0.2534), *domeMeso>GFP/+* (48h, 1%IF, n=26, green, p=0.0074; blue, p=0.7108; red, p<0.0001), **(L)** Graph representing the quantification of total LG area, *domeMeso>GFP/+* (control, 48h, RF, n=24), *domeMeso>GFP/+* (48h, 1%CF, n=21, p=0.0780), *domeMeso>GFP/+* (48h, 1%IF, n=24, p=0.1525), **(M)** Graph showing quantification of differentiation profile based on relative % area with respect to total lymph gland (LG) area at 72h, *domeMeso>GFP/+* (control, 72h, RF, n=22), *domeMeso>GFP/+* (72h, 1%CF, n=12, green, p=0.0667; blue, p=0.5133; yellow, p=0.6918; red, p=0.7945), *domeMeso>GFP/+* (72h, 1%IF, n=22, green, p=0.0087; blue, p=0.0071; yellow, p=6696; red, p=0.6119), **(N)** Graph representing the quantification of total LG area, *domeMeso>GFP/+* (control, 72h, RF, n=22), *domeMeso>GFP/+* (72h,

1%CF, n=12, p=0.9266), *domeMeso>GFP/+* (72h, 1%IF, n=20, p<0.0001), **(O)** Graph showing quantification of differentiation profile based on relative % area with respect to total lymph gland (LG) area at 120h, *domeMeso>GFP/+* (control, 120h, RF, n=36), *domeMeso>GFP/+* (120, 1%CF, n=34, green, p<0.0001; yellow, p=0.3007; red, p<0.0001; white, p=0.4120), *domeMeso>GFP/+* (120h, 1%IF, n=22, green, p<0.0001; yellow, p=0.0013; red, p<0.0001; white, p=0.9485), **(P)** Graph representing the quantification of total LG area, *domeMeso>GFP/+* (control, 120h, RF, n=34), *domeMeso>GFP/+* (120h, 1%CF, n=34, p=0.5542), *domeMeso>GFP/+* (120h, 1%IF, n=27, p<0.0001)**.** Data is presented as bar plots (*p<0.05; **p<0.01; ***p<0.001; ****p<0.0001), ordinary one-way ANOVA. Scale bar: 20µm. ‘n’= lymph gland lobes. DAPI marks DNA, Progenitors (Dome$^{+}$; Green), non-progenitors (blue) intermediary progenitors (Dome$^{+}$ Pxn$^{+}$; Yellow), Differentiating population (Pxn$^{+}$; Red), terminally differentiated population (P1$^{+}$; white). Comparisons for significance are with control on RF. LG lobes are outlined with white border along with removal of other accessory tissues for clear demarcation.

**Supplementary figure 10:**

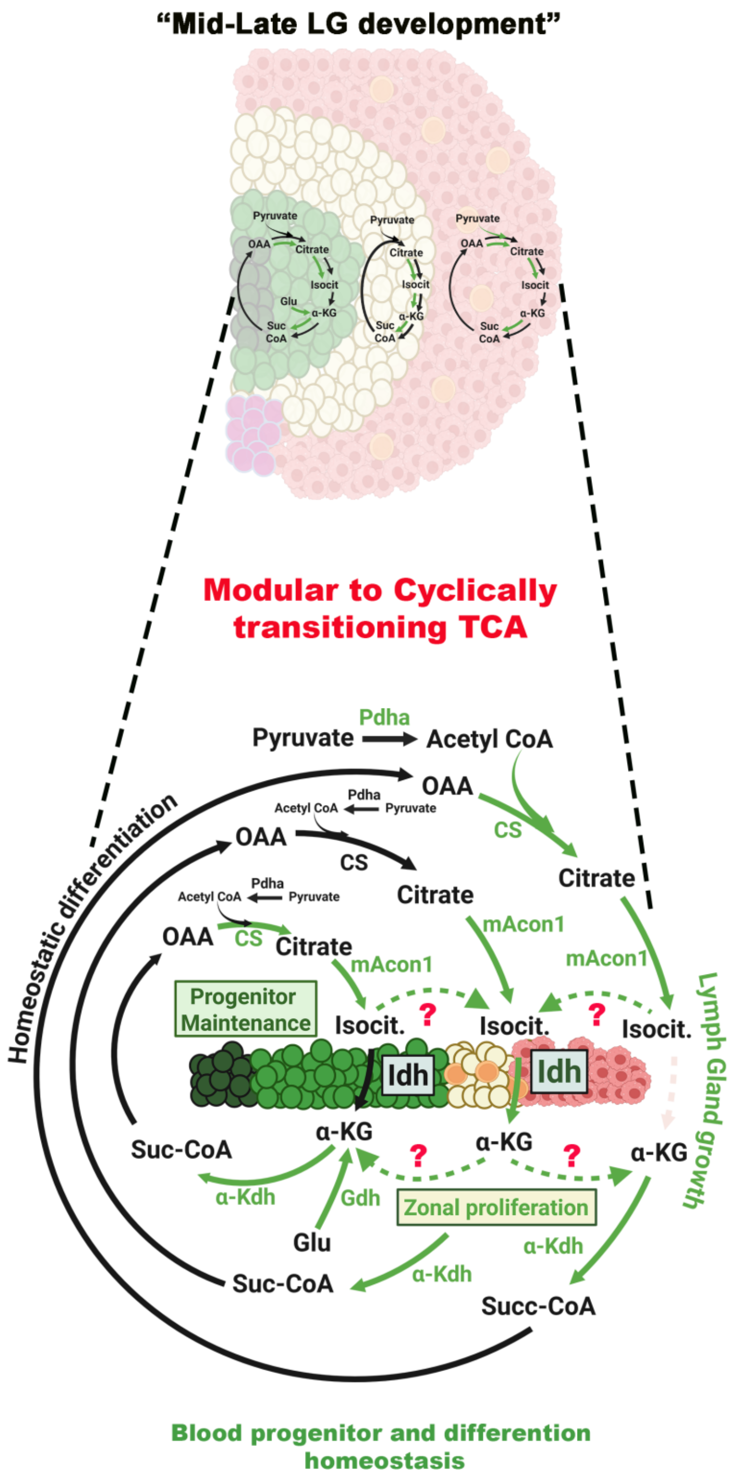

1972

**Supplementary Figure 10**

This schematic illustrates the spatial and developmental reorganization of the tricarboxylic acid (TCA) cycle across the *Drosophila* larval lymph gland during mid-late third instar in different zones, medullary zone (MZ), intermediary zone (IZ) and cortical zone (CZ). Top illustration shows lymph gland architecture with medullary zone (progenitors, green), intermediate zone (transitioning cells, yellow), and cortical zone (maturing hemocytes, red). Metabolic nodes (citrate/isocitrate/α-KG modules) are depicted as operating semi-independently within early progenitors. Bottom schematic represents the shift to a more integrated, cyclical TCA configuration. Key enzymatic steps; *Pdha* (pyruvate dehydrogenase), *CS* (citrate synthase), *mAcon1* (mitochondrial aconitase), *Idh* (isocitrate dehydrogenase), *α-Kdh* (α-ketoglutarate dehydrogenase) are shown with zone-specific activities. Green arrows denote biochemical steps contributing to progenitor proliferation and maintenance, while dashed arrows and question marks indicate unresolved or context-dependent fluxes and black arrows represent steps involved in maintenance across different zones. In differentiating zones, α-KG metabolism via *α-Kdh* and *Gdh* supports zonal proliferation and lineage progression. The overall model proposes that a modular TCA topology supports progenitor quiescence and maintenance, while a cyclic TCA topology emerges as development proceeds, coordinating growth, differentiation, and lymph gland size homeostasis. IZ population perhaps integrate metabolite fluxes from both MZ and CZ and engage full TCA cycling to restore metabolic balance when fluxes or ROS levels deviate from optimal ranges.

**Table S1**

| ***domeMESO-GAL4,UAS-GFP*** | | |
|---|---|---|
| **S.No.** | **Genotype** | **LG with Lamellocyte** |
| 1 | *Control* | 17 % |
| 2 | $CS^{RNAi}$ | 27.45 |
| 3 | $mAcon1^{RNAi}$ | 60 % |
| 4 | $Idh^{RNAi}$ | 35.71 % |
| 5 | $Kdh^{RNAi}$ | 4 % |
| 6 | $Sdh^{RNAi}$ | 32.14 % |
| 7 | $Fum^{RNAi}$ | 36.36 % |
| 8 | $Mdh^{RNAi}$ | 31.57 % |
| ***Tep4-GAL4,UASmcherry*** | | |
| 1 | *Control* | 20.58 % |
| 2 | $CS^{RNAi}$ | 60.6 % |
| 3 | $mAcon1^{RNAi}$ | 72.72 % |
| 4 | $Idh^{RNAi}$ | 80.95 % |
| 5 | $Kdh^{RNAi}$ | 50 % |
| 6 | $skap^{RNAi}$ | 44 % |
| 7 | $Sdh^{RNAi}$ | 25 % |
| 8 | $Fum^{RNAi}$ | 67.74 % |
| 9 | $Mdh^{RNAi}$ | 73.33 % |
| ***$Hml^{\Delta}$-GAL4,UAS-2XEGFP*** | | |
| 1 | *Control* | 17.18 % |
| 2 | $CS^{RNAi}$ | 61.11 % |
| 3 | $mAcon1^{RNAi}$ | 33.33 % |
| 4 | $Idh^{RNAi}$ | 5.55 % |
| 5 | $Kdh^{RNAi}$ | 50 % |
| 6 | $skap^{RNAi}$ | 15 % |
| 7 | $Sdh^{RNAi}$ | 74.35 % |
| 8 | $Fum^{RNAi}$ | 36.66 % |

| | | |
|---|---|---|
| 9 | *Mdh*$^{RNAi}$ | 44.44 % |
| ***ChIZ-GAL4*** | | |
| 1 | *Control* | 24.39 % |
| 2 | *CS*$^{RNAi}$ | 59.25 % |
| 3 | *mAcon1*$^{RNAi}$ | 59.45 % |
| 4 | *Idh*$^{RNAi}$ | 64.51% |
| 5 | *Kdh*$^{RNAi}$ | 56 % |
| 6 | *skap*$^{RNAi}$ | 37.5 % |
| 7 | *Sdh*$^{RNAi}$ | 82.35 % |
| 8 | *Fum*$^{RNAi}$ | 80.76 % |
| 9 | *Mdh*$^{RNAi}$ | 75.51 % |
| 10 | *Pdh*$^{RNAi}$ | 76.74 % |

**Table S1** Frequency of lymph glands showing lamellocytes induction following zone-specific knockdown of TCA cycle enzymes, presented as a proportion of the total lymph gland cohort. *domeMESO-GAL4,UAS-GFP* (pan-MZ progenitors), *Tep4-GAL4,UAS-mCherry* (core progenitors), *CHIZ-GAL4* (IZ), and *HmlΔ-GAL4,UAS-2xEGFP* (CZ).
